# SARS-COV-2 γ variant acquires spike P681H or P681R for improved viral fitness

**DOI:** 10.1101/2021.10.16.464641

**Authors:** Xiang-Jiao Yang

## Abstract

Severe acute respiratory syndrome coronavirus 2 (SARS-COV-2) evolves and generates different variants through a continuously branching model. Four variants of concern have been the major pandemic drivers around the world. One important question is how they may evolve and generate subvariants, some of which may be even more virulent and drive the pandemic further. While investigating how γ (or P.1) variant has been evolving, I noticed the spike substitution P681H in a group of genomes encoding a new subvariant, which has been designated P.1.7. This subvariant has become the dominant P.1 sublineage in Brazil, Italy, Spain and Peru, supporting that P681H confers evolutionary advantage to P.1.7. In Brazil and Peru, P.1.7 was still responsible for ~30% and ~40% cases, respectively, in August 2021. However, it has been competed out by δ1 (a δ subvariant) in both countries, Italy and Spain, suggesting that P.1.7 is not as virulent as δ1. In addition, 160 P.1 genomes possess a related substitution, P681R, and 120 of them encode a new subvariant, designated P.1.8. This P.1 subvariant carries two additional spike substitutions, T470N and C1235F, located at the receptor-binding pocket and cytoplasmic tail of spike protein, respectively. More P.1.8 genomes have been identified than P.1 genomes that encode P681R but not T470N and C1235F, suggesting that these two substitutions improve virulence of P.1.8 subvariant. Some P.1 genomes carry other substitutions (such as N679K, V687L and C1250F) that affect the furin cleavage site or cytoplasmic tail of spike protein. Thus, to improve viral fitness and expand its evolutionary cage, γ variant acquires mutations to finetune the furin cleavage site and cytoplasmic tail of spike protein.

## INTRODUCTION

Coronavirus disease 2019 (COVID-19) is caused by severe acute respiratory syndrome coronavirus 2 (SARS-CoV-2). Immediately after identification of the initial COVID-19 cases in December 2019, SARS-CoV-2 was identified, sequenced and reported in the first two weeks of 2020 [1–5]. The sequence indicates that the virus is homologous to SARS-COV-1, an enveloped, positive-sense and single-stranded RNA beta-coronavirus that caused the severe acute respiratory syndrome (SARS) epidemic, mainly in Asia, from 2003 to 2004 [6]. Also related is Middle-east respiratory syndrome–related coronavirus (MERS), responsible for outbreaks of flu-like illness from 2012 to 2015 in ~20 countries around the world [7]. Despite the sequence similarity, neither SARS-COV-1 nor MERS led to a pandemic. In stark contrast, just two months after its initial discovery in January 2021, SARS-COV-2 caused the pandemic in March 2020, with multiple powerful waves of COVID-19 cases afterwards. This drastic difference among these three homologous viruses raises an important question about what genomic or sequence features make SARS-COV-2 so contagious, virulent and deadly.

One striking molecular feature identified upon the initial SARS-COV-2 genomes were sequenced in January 2020 [1–5] is an obvious furin cleavage site in the spike protein (681-P**RR**A**R/**S-686, where the three arginine residues key to furin recognition are highlighted in bold and the cleave site is shown with a slash; Fig. 1A). The equivalent sites in the spike proteins from SARS-COV-1 and MERS are less optimal (663-VSSL**R/**S-668 and 747-P**R**SV**R/**S-752, respectively; Fig. 1A), with only one or two arginine residues for furin recognition. Notably, compared to the consensus sequence for furin recognition [8,9], even the site in SARS-COV-2 spike protein is still not optimal, thus leaving the potential for further improvement during virus evolution. Interestingly, spike proteins from the common cold alpha-coronaviruses HKU1 and OC43 possess optimal furin cleavage sites, but such sites are much less optimal in two other α-coronaviruses (Fig. 1A). Because both are still highly prevalent in pediatric common cold cases [10], optimal furin sites are not required for these two viruses to become highly transmissible, suggesting that furin cleavage is only one of many factors that contribute to overall viral fitness.

**Figure 1.**
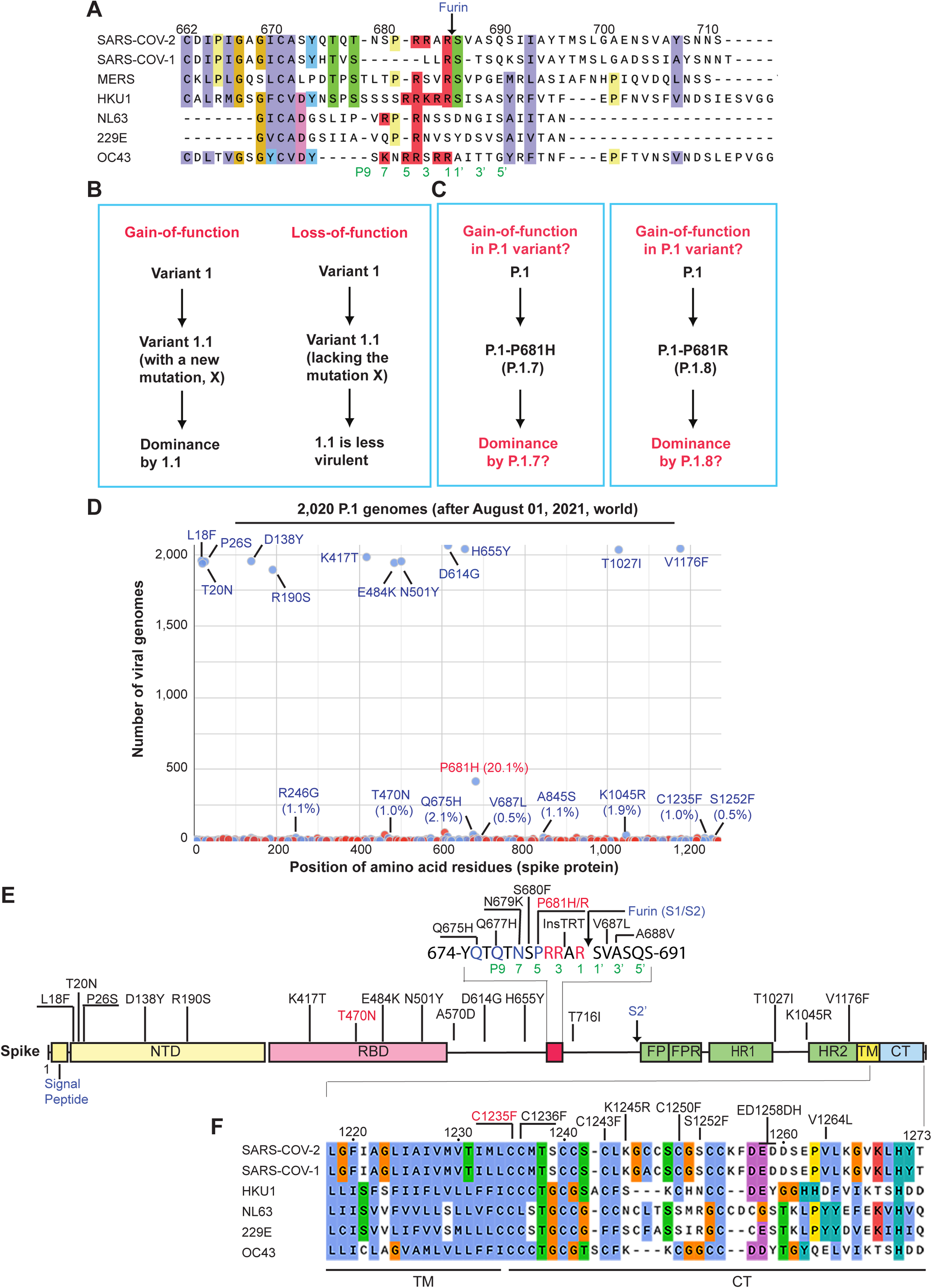
Mutation profile of P.1 genomes. (**A**) Sequence comparison of the furin cleavage site SARS-COV-2 spike protein with the corresponding regions of spike proteins from five other beta coronaviruses, SARS-COV-1, HKU1, NL63, 229E and OC43. (**B**) Schematic representation of two approaches to assess functional impact of specific viral mutations. (**C**) Schematic illustration on the potential use of the ‘gain-of-function’ approach to assess functional impact of the spike substitutions P681H and P681R. (**D**) Mutation profile of 2,020 P.1 genomes identified after August 01, 2021 in different countries around the world. The genomes were downloaded from the GISAID SARS-COV-2 genome sequence database on September 08, 2021 for mutation profiling via Coronapp. Shown here are spike substitutions. (**E**) Domain organization of spike protein and its substitutions encoded by P.1 genomes. The sequence of the furin cleavage site is shown above the domain organization. The nine residue positions before and after the cleavage site are indicated by P1 to P9 and P1’ to P5’, respectively. NTD, N-terminal domain; RBD, receptor-binding domain; S1/S2, boundary of S1 and S2 domains generated by furin cleavage; FP, fusion peptide; FPR, fusion peptide C-terminal proximal region; HR1 and HR2, heptad-repeat regions 1 and 2, respectively; S2’, a cleavage site within S2 domain; TM, trans-membrane motif; CT, C-terminal cytoplasmic tail. (**F**) Sequence comparison of the cytoplasmic tail (and a portion of the transmembrane domain) of SARS-COV-2 spike protein with the corresponding regions of spike proteins from five other beta coronaviruses, SARS-COV-1, HKU1, NL63, 229E and OC43.

SARS-CoV-2 has evolved dynamically and yielded many variants, with some being the main drivers of the pandemic and causing waves of COVD-19 cases. There are now four variants of concern, B.1.1.7 (α) [11], B.1.351 (β) [12], P.1 (γ) [13] and B.1.617.2 (δ) [14] initially identified in the United Kingdom, South Africa, Brazil and India/Japan, respectively. Among these variants, α and δ possess the spike substitutions P681H and P681R, respectively (replacing the proline residue at P5 position of the furin recognition site [8,9], Fig. 1A), but neither β nor γ variants encode such mutations, raising the possibility that these two variants may acquire new mutations to improve the furin cleavage site. This possibility is also related to the important question is how different variants may evolve and generate more virulent subvariants. To address this question, I have tracked over four million SARS-COV-2 genomes in the global initiative on sharing avian influenza data (GISAID) database [15]. While trying to elucidate how γ variant may have been evolving in Brazil and other countries, I noticed the spike substitution P681H in a large group of P.1 genomes. In Brazil, this substitution was present in 29% SARS-COV-2 genomes sequenced in August 2021. In addition, there are about 160 P.1 genomes encoding a related substitution, P681R. Here, I present mutational and phylogenetic analyses of P681H- or P681R-encoding P.1 genomes. The results suggest that acquisition of P681H or P681R confers evolutionary advantage to γ variant.

## Results AND DISCUSSION

### Spike P681H as a major new substitution acquired by γ variant

SARS-COV-2 has infected almost 240 million individuals (according to the Our World in Data website https://ourworldindata.org/, accessed on October 15, 2021). The viral genomes from over 4 million cases have been sequenced and deposited into the GISAID database [15]. This valuable resource offers an unprecedented opportunity to carry out digital genetic analysis for annotation of SARS-COV-2 mutations through the classical “gain-of-function” and “loss-of-function” approaches (Fig. 1B). I utilized the “gain-of-function” approach to assess the functional impact of the P681H and P681R substitutions (Fig. 1C). For this, 2,020 P.1 genomes identified around the world after August 01, 2021 were downloaded from the GISAID database on September 08, 2021 for analysis via Coronapp, an efficient web-based mutation annotation application [16,17]. As expected, signature substitutions such as K417T, E484K, N510Y, D614G and V1176F were detected (Fig. 1D). In addition, P681H is a prominent spike substitution present in 20% of the genomes (Fig. 1D). Other spike substitutions such as T470N, Q675H, V687L, C1235F and C1252F were also identified (Fig. 1D), but they are present in 0.5-2% genomes. While T470N is located at the receptor-binding domain, the other four substitutions alter the furin cleaveage site or the cytoplasmic tail (Fig. 1E-F). The P681H-encoding genomes correspond to a new P.1 subvariant, which has received the Pango lineage designation P.1.7 (https://cov-lineages.org/) [18]. According to the GISAID database, cases with such genomes first appeared in Italy in January 2021. This subvariant was initially described in an important preprint released on July 03, 2021 by scientists from Brazil [19,20].

Phylogenetic analysis of P.1.7 genomes identified by March 2021 revealed that those from Italy form one cluster (Fig. 2A). Many early P.1.7 genomes from Brazil and Peru form two other clusters, which are highly related to that from Italy. The fourth cluster is more heterogenous and contains P.1.7 genomes from Brazil, the USA and several other countries. These four clusters of cases may thus serve as precursors for other P.1.7 cases that have been identified after March 2021. P.1.7 subvariant has acquired additional mutations. For example, 65 of 67 P.1.7 genomes from Spain encode an additional spike substitution, Q613H (the GISAID database, accessed on September 08, 2021). This is adjacent to D614G, a key substitution that defines the entire course of the pandemic [21–23]. Moreover, D14G alters conformational state of the spike protein [24]. Thus, Q613H may synergize with D614G and be important. Indeed, after July 01, 2021, 88 P.1 genomes were sequenced and 40 of them were found to encode both P681H and Q613H, supporting that they improve the fitness (the GISAID database, accessed on September 08, 2021). This raises the question how P681H and Q613H individually contribute to the improvement. Analysis of P.1 genomes from Brazil and Italy suggests that at least P681H confers evolutionary advantage to P.1.7 subvariant (see below). Among 3,827 genomes sequenced in Spain after August 01, 2021 (GISAID database, accessed on September 08, 2021), 18 encode the P.1.7-Q613H subvariant and 3,621 are due to δ variant. In the country, δ variant drives the pandemic mainly through δ1 subvariant [25]. Thus, δ1 subvariant is still predominant whereas the P.1.7 subvariant is relatively obscure in Spain.

**Figure 2.**
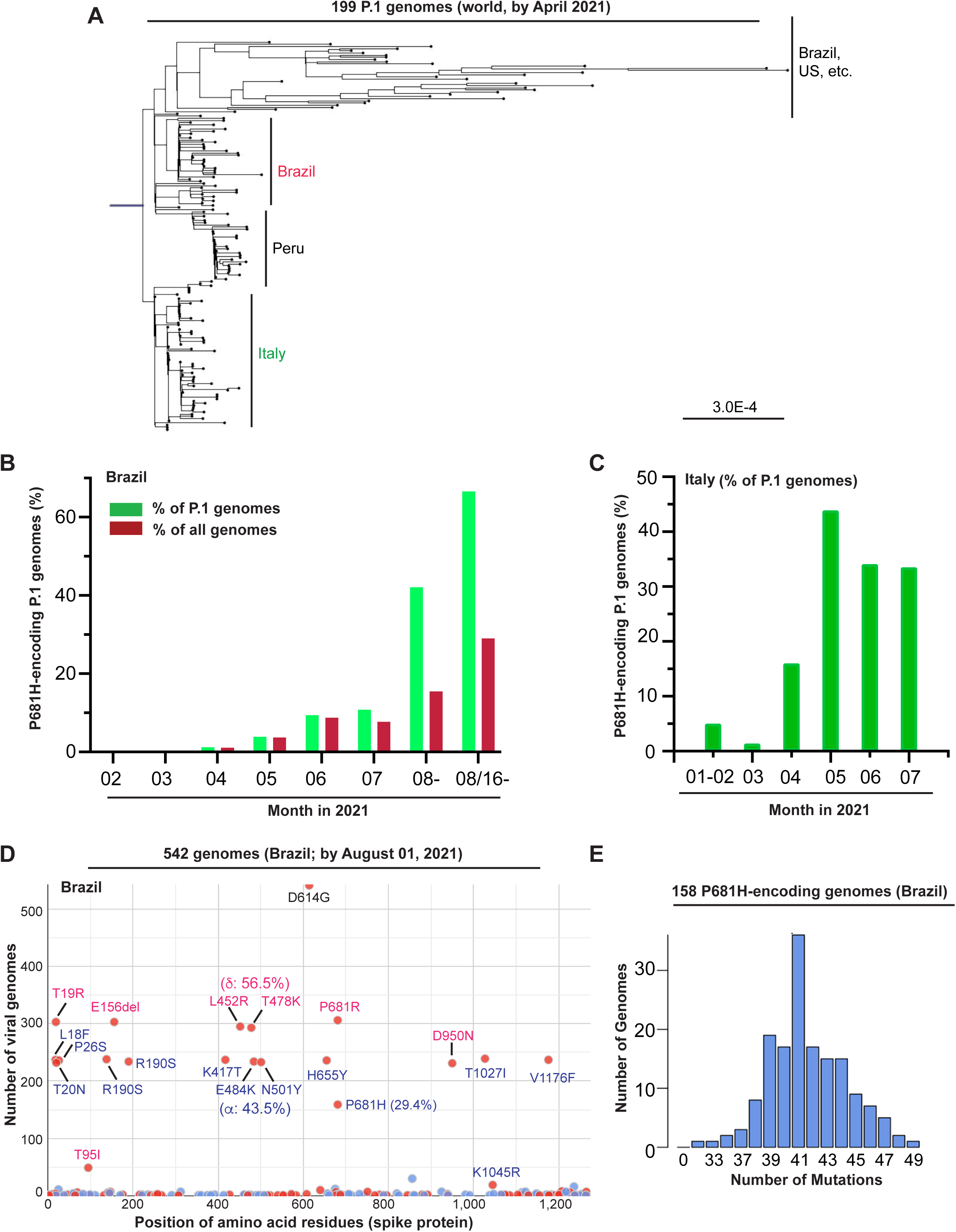
Emergence of P.1.7 as the dominant P.1 subvariant. (**A-B**) Exponential growth of P.1.7 cases in Brazil (A) and Italy (B). This subvariant was still responsible for ~29% new cases identified in Brazil during August 2021, but it had been overtaken by δ variant in Italy. (**C**) Mutation profile of SARS-COV-2 genomes corresponding 542 cases identified in Brazil from August 16 to September 01, 2021. 56.5% and 43.5% of the genomes encode δ and γ variants, respectively. The genomes were downloaded from the GISAID database on September 08, 2021 for mutation profiling via Coronapp. Shown here are spike substitutions. (**D**) Mutation load of P681H-encoding genomes. To generate the distribution, 158 P681H-encoding genomes identified in Brazil from August 16 to September 01, 2021 were used for analysis by Coronapp. (**E**) Phylogenetic analysis of 199 P.1 genomes identified by April 2021. The genomes were downloaded from the GISAID database on September 05, 2021. The phylogenetic analysis package RAxML-NG was used to generate 20 maximum likelihood trees and one bestTree. Figtree was used to display these trees for manual inspection and re-rooting. Presented here is the bestTree. The strain names and GISAID accession numbers of the genomes are provided in Fig. S1.

### Spike P681H confers evolutionary advantage to γ variant

Different from those in Spain, P.1.7 genomes identified in Brazil and Italy carry P681H but not Q613H. Thus, comparison of P.1.7 cases with those carrying parental P.1 genomes should shed light on the contribution of P681H to improvement of viral fitness. As shown in Fig. 2A, there was an exponential growth of P.1.7 cases in Brazil. P.1.7 genomes accounted for over 60% P.1 genomes identified after August 16, 2021 (Fig. 2A). This was also apparent from March to May 2021 in Italy (Fig. 2B). In Brazil, this subvariant was responsible for slightly less than 10% of all COVID-19 genomes sequenced from June-July 2021, and this increased to ~29% from August 16 to September 01, 2021 (GISAID database, accessed on September 08, 2021; Fig. 2A). These results suggest that P681H confers evolutionary advantage to P.1.7.

Compared to those from Brazil, P.1.7 genomes from Italy carry extra mutations. 71.7% P.1.7 genomes sequenced in Italy after June 01, 2021 encode A534V substitution in NSP3 (GISAID database, accessed on September 08, 2021). In addition, there are two silent mutations in the spike gene (N140N and C760C in 98% and 76% genomes, respectively). By August 01, 2021, δ1 subvariant had overtaken P.1.7 and other variants as the dominant pandemic driver in Italy [25]. This is similar to what occurred in Spain (see above). Thus, despite P681H, P.1.7 is still less virulent than δ1 subvariant.

I then utilized Coronapp [16,17] to carry out mutation profiling of 542 SARS-COV-2 genomes identified in Brazil from August 16 to September 01, 2021 (GISAID database, accessed on September 08, 2021). This profiling revealed that 56.5% and 43.5% of them encode δ and γ variants, respectively (Fig. 2D). 29.4% of all genomes identified in Brazil from August 16 to September 01, 2021 carry P681H (Fig. 2D). On the average, these P.1.7 genomes carry 41 mutations per genome (Fig. 2E). Thus, P.1.7 was still an important pandemic driver in Brazil by August 2021. But this has changed drastically afterwards. 3,747 SARS-COV-2 genomes have been sequenced for COVID-19 cases identified in the country during September 2021 (GISAID database, accessed on October 15, 2021). Among these genomes from Brazil, 3,407 and 284 correspond to δ and γ variants, respectively. 107 of the genomes (~38% of P.1 genomes) encode P.1.7 subvariant. These results support that although acquisition of P681H confers evolutionary advantage, P.1.7 is still much less virulent than δ variant.

### Spike P681R is encoded by a small group of P.1 genomes

An arginine residue is preferred at position P5 of the furin recognition site (Fig. 1A) [8,9], so I asked whether there are P.1 genomes encoding P681R. Indeed, there are over 160 such genomes (Fig. 3; GISAID database, accessed on September 08, 2021). Mutation profiling via Coronapp identified I441V as an extra NSP3 substitution in over 120 genomes (Fig. 3A). Interestingly, these genomes also encode two extra spike substitutions, T470N and C1235F (Fig. 3B), which are located at the receptor-binding pocket and the cytoplasmic tail of spike protein, respectively (Fig. 1E-F). The P.1 subvariant encoding these three extra substitutions have been designated P.1.8 (https://cov-lineages.org/) [18]. Except for a silent mutation alters the codon for S201, no extra nucleocapsid substitutions are prominent (Fig. 3C).

**Figure 3.**
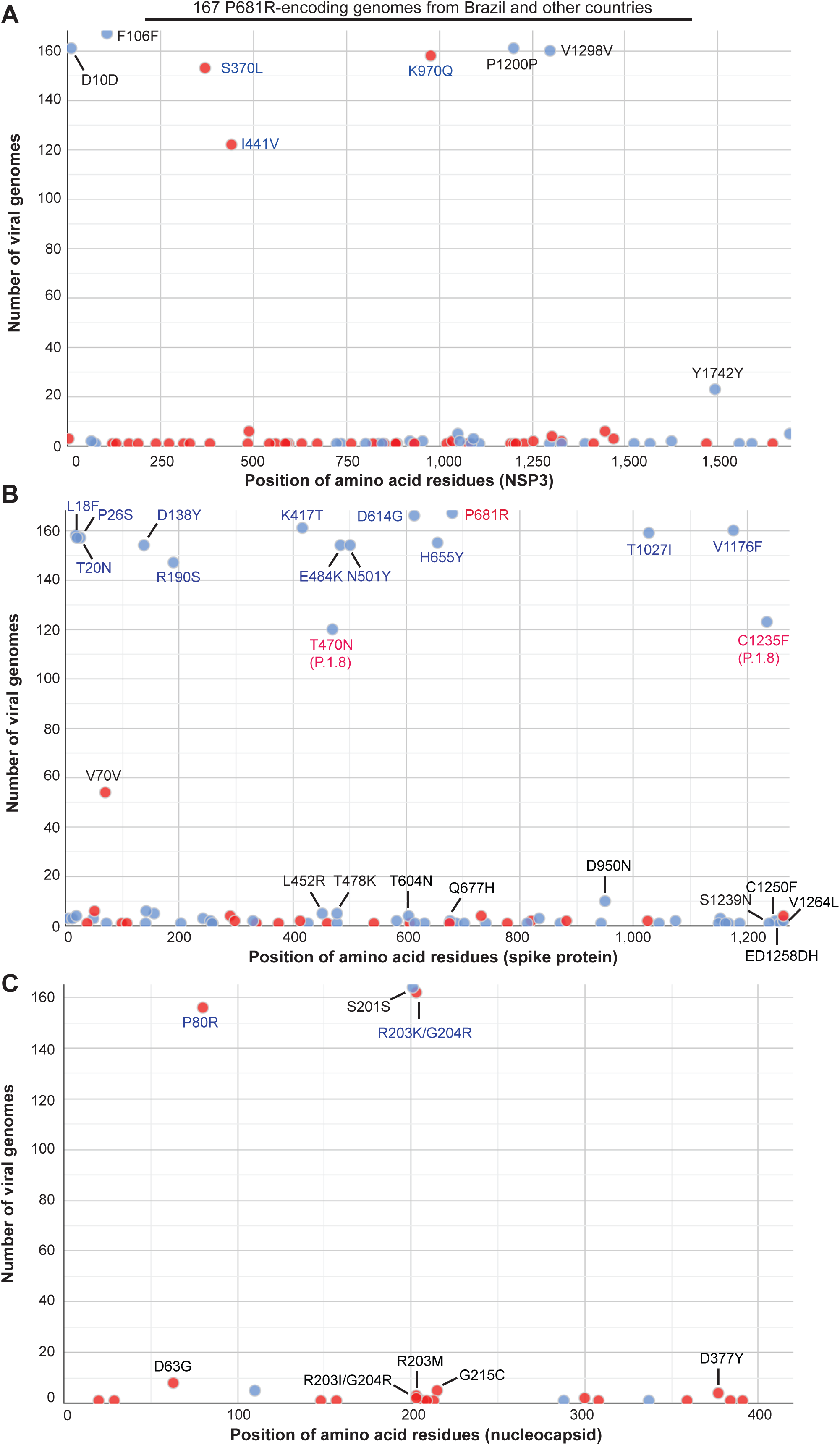
Mutation profile of 167 P681R-encoding P.1 and P.1-like genomes identified in Brazil and and other countries around the world. As in Fig. 1C, the genomes were downloaded from the GISAID SARS-COV-2 genome sequence database on September 05, 2021 for mutation profiling via Coronapp. Shown in (A), (B) and (C) are substitutions in NSP3, spike and nucleocapsid proteins, respectively.

On the average, these P.1.8 genomes carry 44 mutations per genome (Fig. 4A). This is three more than the average mutation load of 41 per P.1.7 genome (Fig. 2E) and 6 more than the average mutation load of 41 per P.1-N679K genome (Fig. 4B). On the average, P.1 variant carries 35-38 mutations per genome, so P.1.8 is the most mutated P.1 subvariant. For comparison, C.1.2 variant is a highly mutated variant and carries 46 mutations per genome [26,27]. Although δ variant typically carries about 38-41 mutations per genome, some subvariants harbor 46-50 mutations per genome [25,28,29]. Thus, P.1.8 is the most mutated P.1 subvariant but is not one of the most mutated SARS-COV-2 variants.

**Figure 4.**
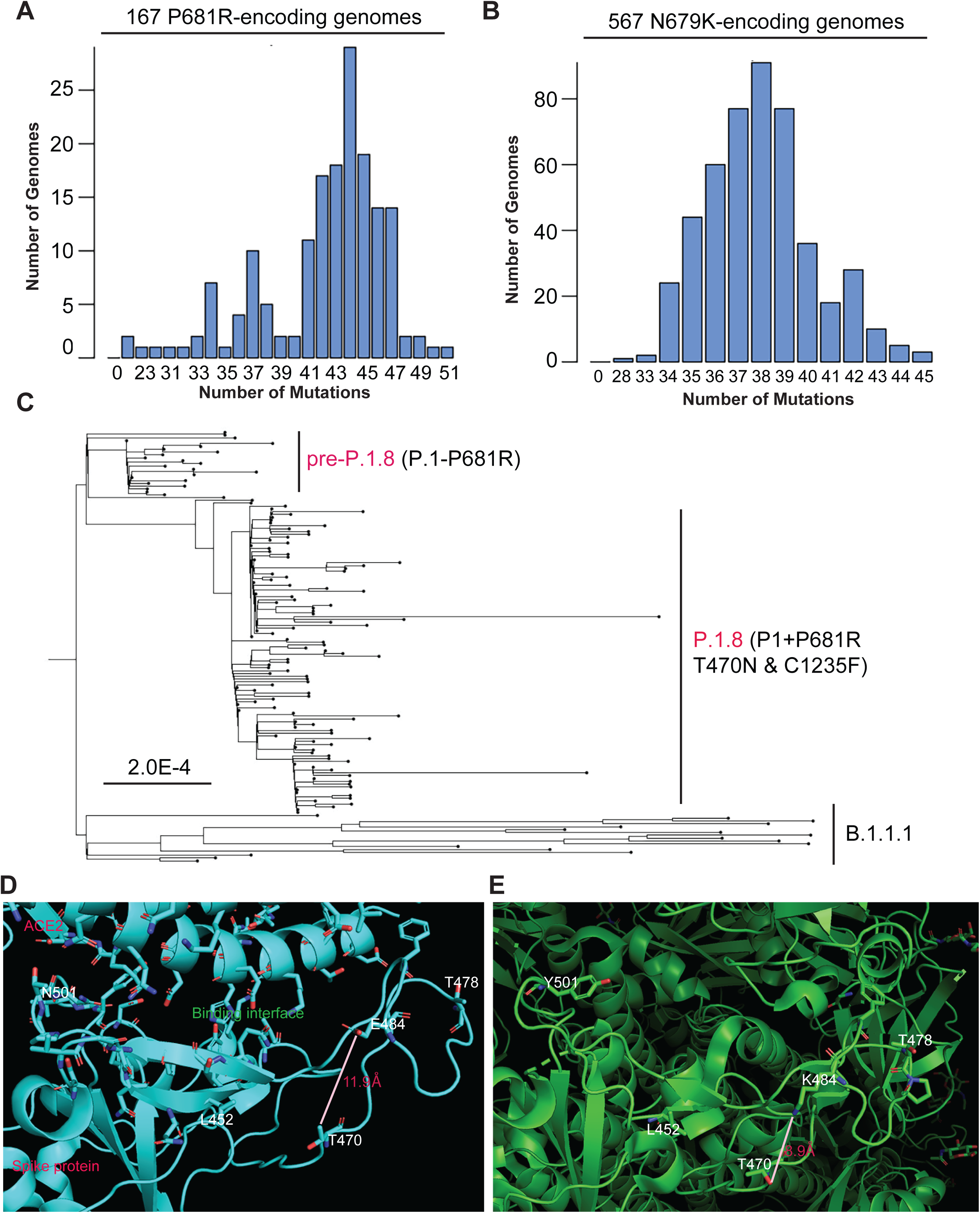
Mutation profiling and phylogenetic analysis of P681R-encoding P.1 genomes. (**A**) Mutation load of P681R-encoding genomes. To generate the distribution, 167 P681R-encoding P.1 and P.1-like genomes identified in Brazil and other countries were used for analysis by Coronapp. The genomes were downloaded from the GISAID database on September 07, 2021. (**B**) Mutation load of N679K-encoding genomes. To generate the distribution, 476 N679K-encoding P.1 genomes identified in Brazil and other countries were used for analysis by Coronapp. (**C**) Phylogenetic analysis of the 167 P.1 and P.1-like genomes. RAxML-NG was used to generate 20 maximum likelihood trees and one bestTree. Figtree was used to display these trees for manual inspection and re-rooting. Presented here is the bestTree. The strain names and GISAID accession numbers of the genomes are provided in Fig. S2. (**D**) Structural details showing the interface between ACE2 and the receptor-binding pocket of spike protein. T470 is away from the pocket and 11.9 Å from E484. Adapted from PyMol presentation of the spike protein-ACE2 complex structure 6M17 from the PDB database. (**E**) In spike protein from γ variant, T470 is 8.9 Å away from K484. T470N makes this distance even shorter. Adapted from PyMol presentation of γ variant spike protein structure 7M8K from the PDB database.

I next carried out phylogenetic analysis of the 167 P.1.8 and P.1.8-like genomes. For this, RAxML-NG was used to generate maximum likelihood trees and one bestTree. Figtree was used to display these trees for manual inspection and re-rooting. As shown in Fig. 4C, P.1-P681R and P.1.8 genomes form distinct branches, away from a group of related genomes that also encode E484K, N501Y and P681R. The P.1.8 branch is much larger than the P.1-P681 branch, so the NSP3 substitution I441V and the spike substitutions T470N and C1235F make P.1.8 more virulent.

According an NSP3 structural model (QHD43415, built by Dr. Yang Zhang’s group at University of Chicago, https://zhanglab.ccmb.med.umich.edu/COVID-19/), I441 is in vicinity with V462 and V482. These three residues thus form a hydrophobic core and I441V is expected to improve this core. As shown in Fig. 4D are structural details about the interface between ACE2 and the receptor-binding pocket of spike protein. Spike T470 is away from the pocket and 11.9Å from E484. In the spike protein from P.1 variant, T470 is 8.9Å away from K484. The substitution T470N makes this distance shorter. In the parental spike protein, T470N is unfavorable for spike expression and ACE2 binding [30]. It remains unclear how T470N exerts its impact in the context of γ variant. C1235 is located at the Cys-rich region of the cytoplasmic tail (Fig. 1E-F). This region is required for palmitoylation and membrane fusion [31], so C1235F may affect these events. In addition, to C1235F, there are other substitutions at the cytoplasmic tail (Figs 1D-F & 3B). Thus, this tail is often subject to substitutions. In support of this, V1264L (Fig. 1E-F) is a key substitution of a new δ subvariant that has been a predominant pandemic driver in Indonesia, Singapore and Malaysia [29].

### Spike P681H and P681R in improving viral fitness of SARS-COV-2 variants

One important question is relative virulence of P681H- or P681R-encoding P.1 subvariants when compared to δ and other variants. As stated above, the P.1.7 subvariants have been largely competed out by δ in Italy and Spain, indicating that they are not as virulent as δ variant. Consistent with this, the P.1.7 genome number did not increase as dramatically as the δ genome number in Brazil (Fig. 5A). With the initial 6 genomes identified in April 2021, δ variant has quickly become the dominant pandemic driver in the country. Under such an aggressive invasion by δ variant, P.1 variant receded quickly whereas P.1.7 subvariant displayed growth, still accounting for ~29% cases in August 2021 (Fig. 2B). However, among 3,737 SARS-COV-2 genomes sequenced for COVD-19 cases identified in Brazil during September 2021, 91% and 7.6% correspond to δ and γ variants, respectively, with only 2.9% genomes encoding P.1.7 subvariant (GISAID database, accessed on October 15, 2021). These results support that although acquisition of P681H confers evolutionary advantage, P.1.7 is still much less virulent than δ variant (Fig. 5B).

**Figure 5.**
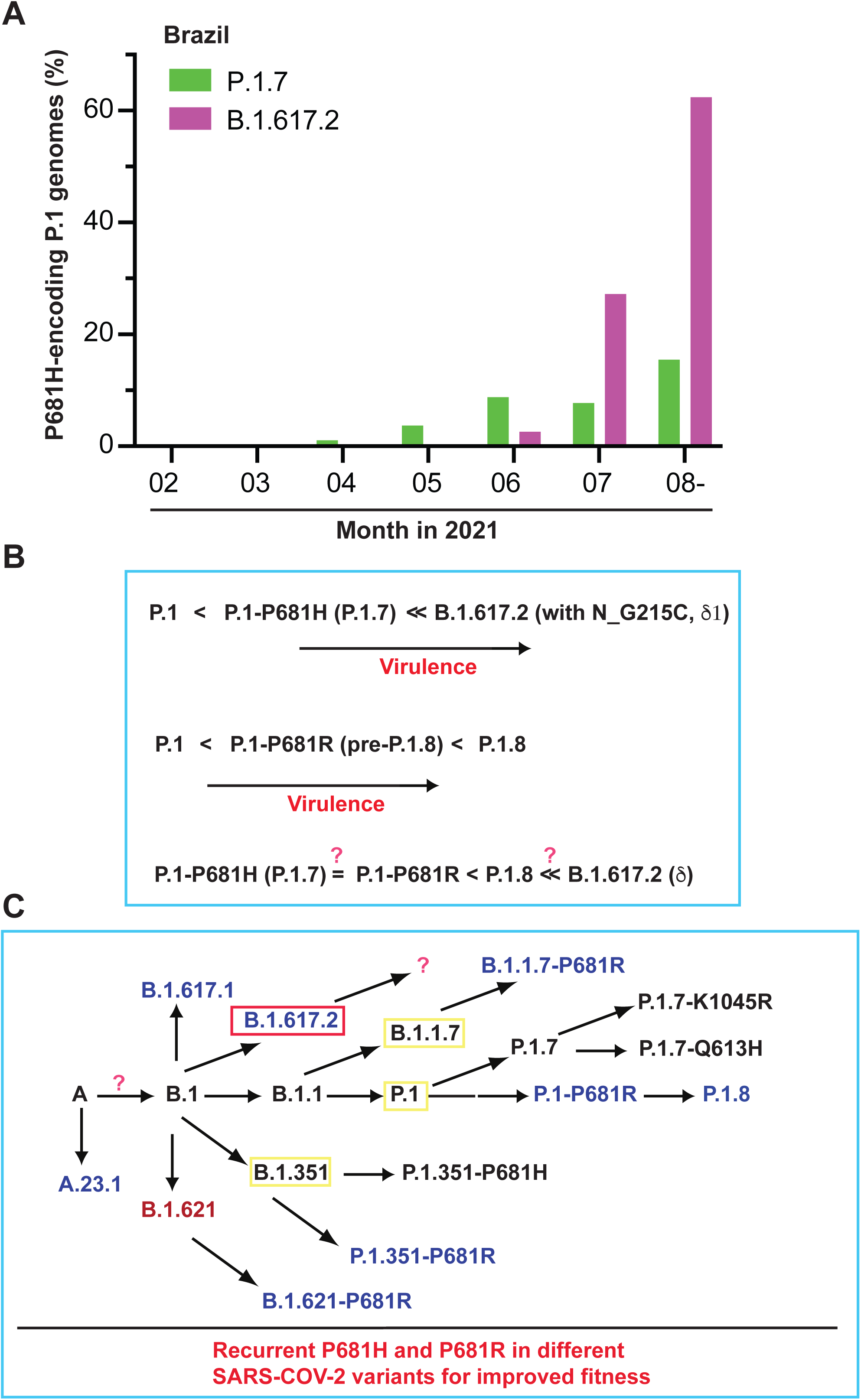
Relative virulence of P681H- or P681R-encoding P.1 subvariants compared to δ variant. **(A)** Differential growth of P.1.7 and δ genomes identified in Brazil. With the initial 6 genomes identified in April, δ variant has quickly become the dominant pandemic driver. Under such an aggressive invasion by δ variant, parental P.1 variant receded quickly whereas the P.1.7 subvariant displayed continuous growth, accounting for 29% genomes sequenced from in August 16 to September 01, 2021. For preparation of the panel, the GISAID database was accessed on September 08, 2021. (**B**) Schematic representation showing relative virulence of P681H- or P681R-encoding P.1 subvariants and δ variant. (**C**) Schematic representation of how SARS-COV-2 has evolved and acquired spike P681H or P681R. Both B.1.617.1 and B.1.617.3 variants encode P681R but are much less virulent than δ variant. Thus, while P681R is important in making δ variant the most virulent pandemic driver, this substitution itself is insufficient and requires other mutations to improve the overall viral fitness of this variant. Related to this, acquisition of P681H or P681R confers evolutionary advantage to P.1 subvariants, but these subvariants such as P.1.7 and perhaps also P.1.8 are still less virulent than δ variant.

As illustrated in Fig. 5C, there are many SARS-COV-2 variants possessing P681H or P681R. Unlike β and γ variants, α variant possesses P681H, suggesting the importance of this substitution in making this variant much more virulent than β and γ variants. This is consistent with the results that P.1.7 is more virulent than P.1 itself (Fig. 2B-C). In the GISAID database (accessed on September 08, 2021), there are 2,243 α variant genomes encoding P681R, with the first 13 cases identified December 2021. Because this subvariant has not become a dominant pandemic driver, P681R does not confer clear evolutionary advantage to α variant (Fig. 5B). One possibility is that P681H is sufficient for α variant in terms of furin cleavage efficiency. Of relevance, B.1.617.1 and B.1.617.3 variants encode P681R but are much less virulent than δ variant [14]. Moreover, A.23.1 variant also encodes P681R. After the initial identification of this variant in Africa in October 2020, it has spread to other continents [32]. According to the GISAID database, few new A.23.1 genomes have been identified June 2021.

Moreover, B.1.466.2 encodes P681R but it did not really become a major pandemic driver around the world [29]. Furthermore, among common cold coronaviruses, some do not show optimal furin cleavage sites (Fig. 1A). Thus, while P681R is important in making δ variant the most virulent pandemic driver, this substitution *per se* is insufficient and requires other mutations in enhancing overall viral fitness.

In summary, γ variant has evolved and yielded multiple subvariants. Two prominent subvariants are P.1.7 and P.1.8, encoding the extra spike substitutions P681H and P681R, respectively. Both are expected to improve furin cleavage of spike protein (Fig. 1A & 1E-F). Although acquisition of P681H confers evolutionary advantage, P.1.7 is still much less virulent than δ variant (Fig. 5A-B). P.1.8 also encodes T470N and C1235F, located at the receptor-binding domain and cytoplasmic tail of spike protein, respectively. There are P681R-encoding genomes carrying additional substitutions at the cytoplasmic tail (Fig. 3B). Thus, γ variant frequently alters the furin cleavage site and cytoplasmic tail of spike protein to improve its viral fitness. Thus, in addition to the receptor-binding domain of spike protein, the furin cleavage site and cytoplasmic tail are two important elements that demarcate the space of γ variant’s evolutionary cage [29].

## Supporting information

Acknowledgement table on the GISAID genomes used in this study

Acknowledgement table on the GISAID genomes used in this study

Acknowledgement table on the GISAID genomes used in this study

## ACKNOWLEDGEMENT

I gratefully acknowledge the GISAID database for maintenance SARS-COV-2 viral genomes and the investigators for the valuable genomes sequences used in this work (see the supplementary section for details). I am grateful to Professor Federico M. Giorgi at University of Bologna, Italy for generosity to allow the access to the Coronapp server. This work was supported by funds from Canadian Institutes of Health Research (CIHR), Natural Sciences and Engineering Research Council of Canada (NSERC) and Compute Canada (to X.J.Y.).

## MATERIALS AND METHODS

### SARS-COV-2 genome sequences, mutational profiling and phylogenetic analysis

The genomes were downloaded the GISAID database on the dates specified in the figure legends. CoVsurver (https://www.gisaid.org/epiflu-applications/covsurver-mutations-app/) was used to analyze mutations on representative SARS-COV-2 genomes. Fasta files containing specific groups of genomes were downloaded from the GISAID database. During downloading, each empty space in the Fasta file headers was replaced by an underscore because such a space makes the files incompatible for subsequent mutational profiling, sequence alignment and phylogenetic analysis, as described with details in two other studies [25,27]. The Fasta headers were shortened and modified further as described [25,27]. The cleaned Fasta file was used for mutational profiling via Coronapp (http://giorgilab.unibo.it/coronannotator/), a web-based mutation annotation application [16,17]. The cleaned Fasta file was also uploaded onto SnapGene (version 5.3.2) for multisequence alignment via the MAFFT tool. RAxML-NG version 0.9.0 [33] was used for phylogenetic as described [25].

### Defining different variant genomes using various markers

α, β, γ, δ and other variant genomes were downloaded from the GISAID database as defined by the server. δ subvariant genomes were defined as described [25]. Briefly, nucleocapsid substitutions G215C and R385K (Table 1) were used as markers for δ1 or δ3 genomes, respectively. Spike substitutions A222V and K77T were used as markers for δ2 or δ4 genomes, respectively. In Europe, there are many δ1V genomes that also encode spike A222V, so the NSP3 substitution P822L was used together with spike A222V to identify δ2 genomes. As discussed previously [25], there are several limitations with these markers. But they should not affect the overall conclusions.

### PyMol structural modeling

The PyMol molecular graphics system (version 2.4.2, https://pymol.org/2/) from Schrödinger, Inc. was used for downloading structure files from the PDB database for further analysis and image generation. Structural images were cropped via Adobe Photoshop for further presentation through Illustrator.

## SUPPLEMENTAL INFORMATION

This section includes 1) two supplementary figures with detailed information on two phylogenetic trees, and 2) three acknowledgement tables for the GISAID genomes used in this work.

## SUPPLEMENTAL FIGURE LEGENDS

**Figure S1.**
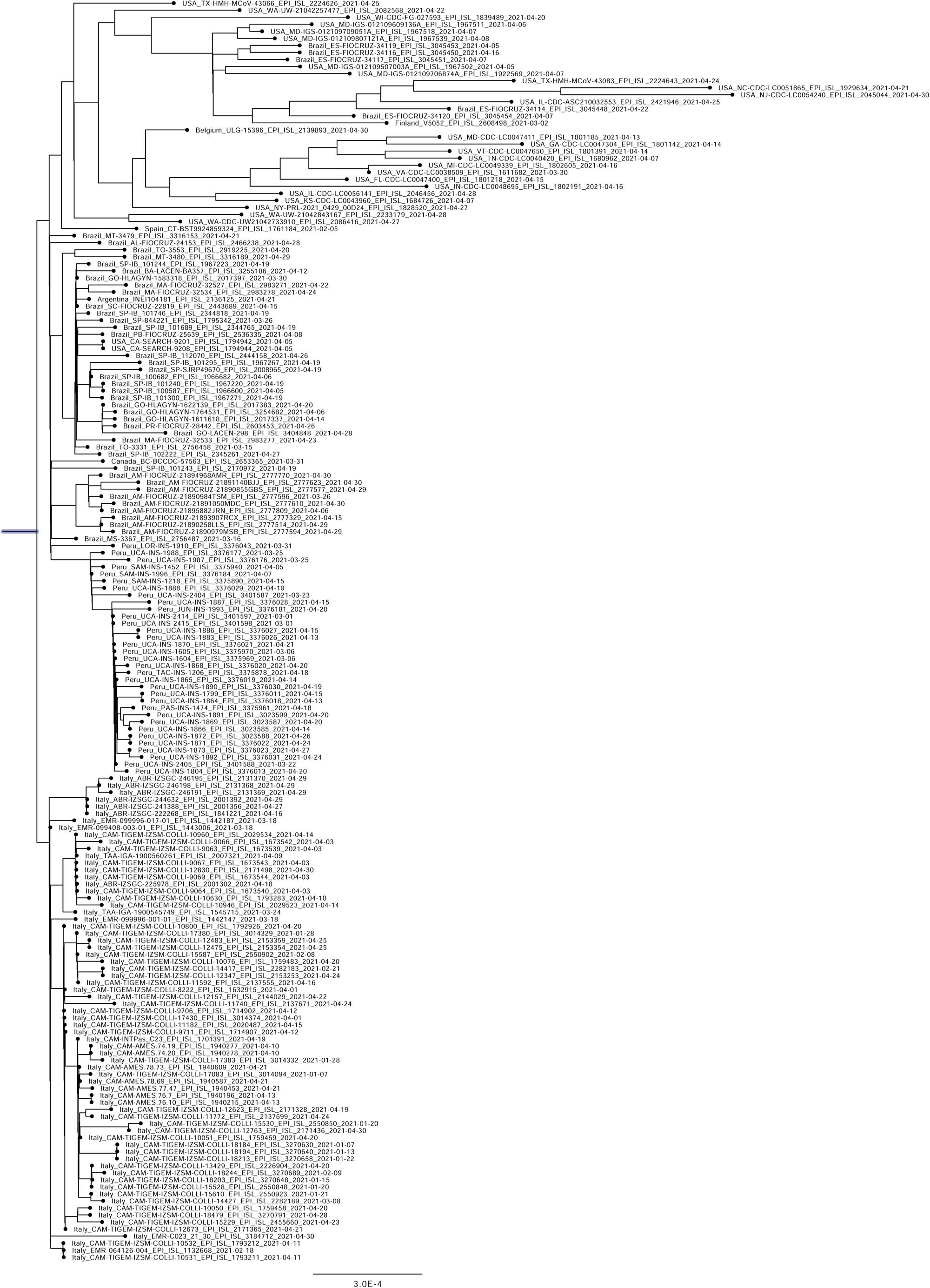
Phylogenetic analysis of 199 P.1 genomes identified by April 2021. A Fasta file containing the genomes was downloaded from the GISAID database. See Fig. 2D for comparison.

**Figure S2.**
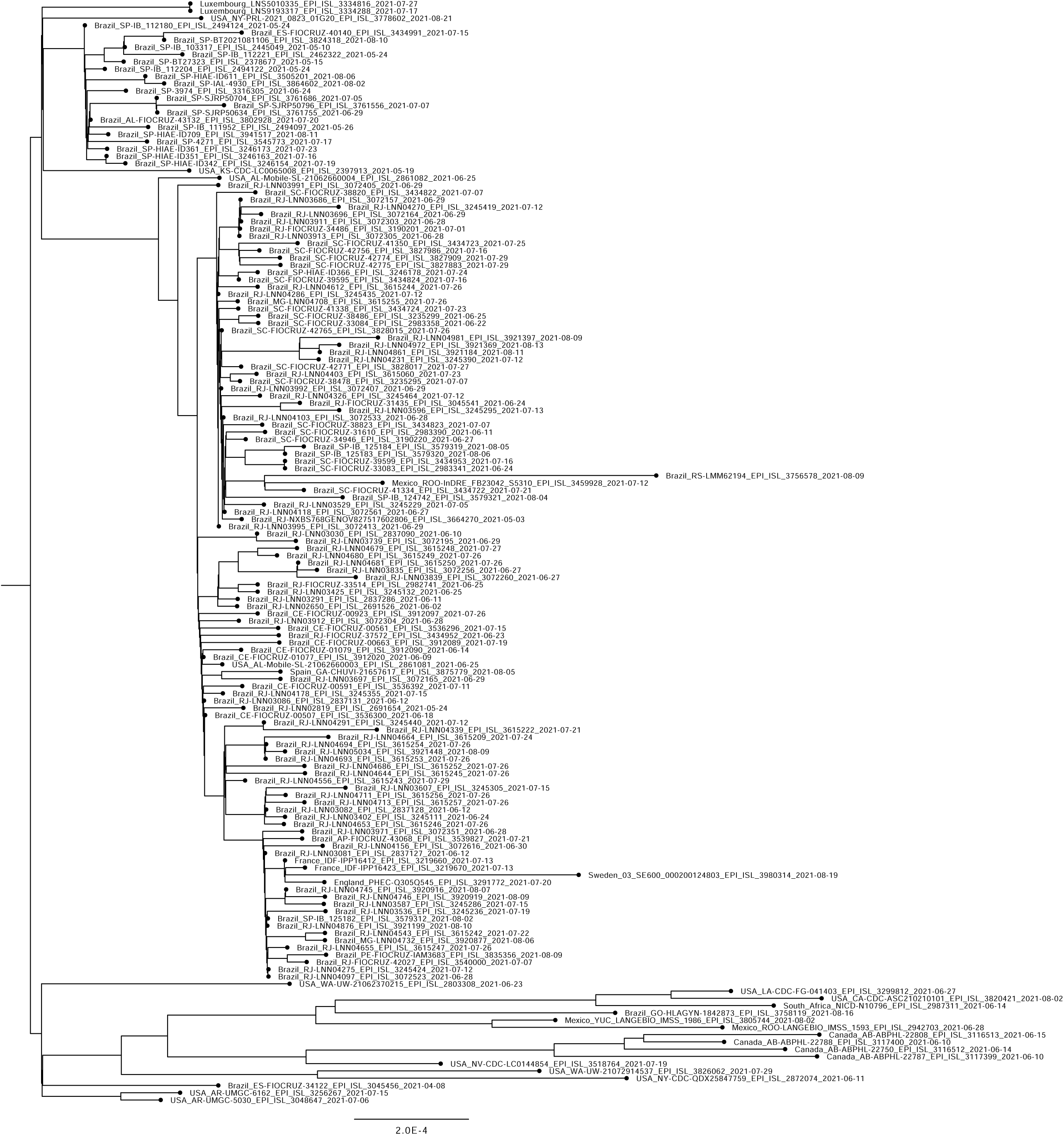
Phylogenetic analysis of 167 P.1 and P.1-like genomes identified by September 07, 2021. A Fasta file containing the genomes was downloaded from the GISAID database. See Fig. 4C for comparison.

