## Supplementary material for "SARS-COV-2 γ variant acquires spike P681H or P681R for improved viral fitness": Acknowledgement table on the GISAID genomes used in this study

We gratefully acknowledge the following Authors from the Originating laboratories responsible for obtaining the specimens, as well as the Submitting laboratories where the genome data were generated and shared via GISAID, on which this research is based.

All Submitters of data may be contacted directly via www.gisaid.org

Authors are sorted alphabetically.

| Accession ID | Originating Laboratory | Submitting Laboratory | Authors |
|---|---|---|---|
| EPI_ISL_3579321 | AFIP | Instituto Butantan | Antonio Jorge Martins; Claudia Renata dos Santos Barros; David Schlesinger; Debora Botequio Moretti; Dimas Tadeu Covas; Elaine Cristina Marqueze; Elaine Vieira Santos; Evandra Strazza Rodrigues; Heidge Fukumasu; Jayme Augusto de Souza-Neto; José Salvatore Leister Patané; Luiz Alcantara; Luiz Lehmann Coutinho; Maria Carolina Elias; Maurício Lacerda Nogueira; Rafael dos Santos Bezerra; Raul Machado Neto; Rejane Maria Tommasini Grotto; Ricardo Haddad; Sandra Coccuzzo Sampaio Vessoni; Simone Kashima; Svetoslav Nanev Slavov; Vincent Louis Viala |
| EPI_ISL_2494097 | AFIP SUL | Instituto Butantan | Antonio Jorge Martins; Claudia Renata dos Santos Barros; David Schlesinger; Debora Botequio Moretti; Dimas Tadeu Covas; Elaine Cristina Marqueze; Elaine Vieira Santos; Evandra Strazza Rodrigues; Heidge Fukumasu; Jayme Augusto de Souza-Neto; José Salvatore Leister Patané; Luiz Alcantara; Luiz Lehmann Coutinho; Maria Carolina Elias; Maurício Lacerda Nogueira; Rafael dos Santos Bezerra; Raul Machado Neto; Rejane Maria Tommasini Grotto; Ricardo Haddad; Sandra Coccuzzo Sampaio Vessoni; Simone Kashima; Svetoslav Nanev Slavov; Vincent Louis Viala |
| EPI_ISL_3820421 | Aegis Sciences Corporation | Centers for Disease Control and Prevention Division of Viral Diseases, Pathogen Discovery | Adrian Paskey; Alec Vest; Benjamin Rambo-Martin; Christopher Gulvick; Clinton Paden; Cyndi Clark; Dakota Howard; Darlene Wagner; Dhwani Batra; Dillon Nall; Duncan MacCannell; Ethan Sanders; Holly Houdeshell; Jason Caravas; Kara Moser; Matthew Hardison; Matthew Schmerer; Ola Kvalvaag; Patrick Campbell; Peter Cook; Rob Case; Scott Sammons; Shatavia Morrison; Shaun Westlund; Vikramsinha Ghorpade; Yvette Unoarumhi |
| EPI_ISL_3980314 | Akademiska Sjukhuset, Clinical Microbiology and Hospital Hygiene | Akademiska Sjukhuset, Clinical Microbiology and Hospital Hygiene | Jonathan Haars; Julia Bergholm; Patrik Ellström; Rene Kaden; Steinar Mannsverk |
| EPI_ISL_3048647, EPI_ISL_3256267 | Arkansas Public Health Laboratory, Arkansas Department of Health | University of Minnesota Genomics Center | Corbin Dirkx; Daryl M. Gohl; Jaquelyn Kuriger-Laber; John Garbe |
| EPI_ISL_3334288 | BioneXt Lab | Laboratoire national de sante, Microbiology, Microbial Genomics Platform | Anke Wienecke-Baldacchino; Catherine Ragimbeau; Elodie Solarino; Fatu Djabi; Jessica Tapp; Lise Pignon; Raoul Salmon; Tamir Abdelrahman; Thibault Ferrandon; Virginie Jover |
| EPI_ISL_3579320 | CENTRO DE SAUDE II | Instituto Butantan | Antonio Jorge Martins; Claudia Renata dos Santos Barros; David Schlesinger; Debora Botequio Moretti; Dimas Tadeu Covas; Elaine Cristina Marqueze; Elaine Vieira Santos; Evandra Strazza Rodrigues; Heidge Fukumasu; Jayme Augusto de Souza-Neto; José Salvatore Leister Patané; Luiz Alcantara; Luiz Lehmann Coutinho; Maria Carolina Elias; Maurício Lacerda Nogueira; Rafael dos Santos Bezerra; Raul Machado Neto; Rejane Maria Tommasini Grotto; Ricardo Haddad; Sandra Coccuzzo Sampaio Vessoni; Simone Kashima; Svetoslav Nanev Slavov; Vincent Louis Viala |
| EPI_ISL_3664270 | DASA | DASA | Adriano Bonaldi; Angelica Hristov; Annelise Lopes; Bianca Cota; Cristina Oliveira; Jose Levi; Lidia Yamamoto; Paulo Pierry; Rodrigo Guarischi; Rodrigo Salazar |
| EPI_ISL_2987311 | ERMELO LABORATORY | National Institute for Communicable Diseases of the National Health Laboratory Service | Amoako DG; Bhiman JN; Everatt J; Ismail A; Mahlangu B; Mnguni A; Mohale T; Ntuli N; Scheepers C |
| EPI_ISL_3299812 | Fulgent Genetics | Centers for Disease Control and Prevention Division of Viral Diseases, Pathogen Discovery | Adrian Paskey; Becky Tsai; Benafsh Sapra; Benjamin Rambo-Martin; Christopher Gulvick; Clinton R. Paden; Dakota Howard; Darlene Wagner; Dhwani Batra; Doreen Ng; Duncan MacCannell; Harry Gao; James Xie; Jason Caravas; John Gao; Joseph Fierro; Kara Moser; Matthew Schmerer; Mickey Li; Peter W. Cook; Scott Sammons; Shatavia Morrison; Yan Meng; Yvette Unoarumhi |
| EPI_ISL_3912097 | HGCC HOSPITAL GERAL DR CESAR CALS | ACME Lab, Oswaldo Cruz Foundation, FIOCRUZ/CE | Cleber Furtado Aksenen; Fabio Miyajima; Fernando Braga Stehling; Francisco Eder de Moura Lopes; Jamille Maria Mendes Bezerra; Joaquim Cesar do Nascimento Sousa Junior; Pedro Miguel Carneiro Jeronimo; Suzana Porto Almeida & Lucas Delerino on behalf of COVID-19 FIOCRUZ Genomic Network; Thais Ferreira de Oliveira; Thais de Oliveira Costa; Ticiane Cavalcante de Souza; Veridiana Pessoa Miyajima |
| EPI_ISL_3758119 | HLAGYN - Laboratorio de Imunologia de Transplantes de Goias | HLAGYN - Laboratorio de Imunologia de Transplantes de Goias | Alessandro Leonardo Alvares Magalhaes; Erika Lopes Rocha Batista; Fernando Antonio Vinhal dos Santos; Frederico Rodrigues Vinhal; Kamila Oliveira Reis De Freitas.; Lucas Carlos Gomes Pereira; Sabrina Sara Moreira Duarte |
| EPI_ISL_3912089 | HM HOSPITAL DE MESSEJANA DR CARLOS ALBERTO STUDART GOMES | ACME Lab, Oswaldo Cruz Foundation, FIOCRUZ/CE | Cleber Furtado Aksenen; Fabio Miyajima; Fernando Braga Stehling; Francisco Eder de Moura Lopes; Jamille Maria Mendes Bezerra; Joaquim Cesar do Nascimento Sousa Junior; Pedro Miguel Carneiro Jeronimo; Suzana Porto Almeida & Lucas Delerino on behalf of COVID-19 FIOCRUZ Genomic Network; Thais Ferreira de Oliveira; Thais de Oliveira Costa; Ticiane Cavalcante de Souza; Veridiana Pessoa Miyajima |
| EPI_ISL_3912020, EPI_ISL_3912090 | HOSPITAL CURA DARS | ACME Lab, Oswaldo Cruz Foundation, FIOCRUZ/CE | Cleber Furtado Aksenen; Fabio Miyajima; Fernando Braga Stehling; Francisco Eder de Moura Lopes; Jamille Maria Mendes Bezerra; Joaquim Cesar do Nascimento Sousa Junior; Pedro Miguel Carneiro Jeronimo; Suzana Porto Almeida & Lucas Delerino on behalf of COVID-19 FIOCRUZ Genomic Network; Thais Ferreira de Oliveira; Thais de Oliveira Costa; Ticiane Cavalcante de Souza; Veridiana Pessoa Miyajima |
| EPI_ISL_3536295, EPI_ISL_3536392 | HOSPITAL ESTADUAL LEONARDO DA VINCI | Oswaldo Cruz Institute, FIOCRUZ/CE | Cleber Furtado Aksenen; Fabio Miyajima; Fernando Braga Stehling; Francisco Eder de Moura Lopes; Jamille Maria Mendes Bezerra; Joaquim César do Nascimento Sousa Junior; Pedro Miguel Carneiro Jeronimo; Suzana Porto Almeida e Lucas Delerino; Thais Ferreira de Oliveira; Thais de Oliveira Costa; Ticiane Cavalcante de Souza; Veridiana Pessoa Miyajima |
| EPI_ISL_2445049 | INSIDE DIAGNÓSTICOS SUL PARELHEIROS | Instituto Butantan | Antonio Jorge Martins; Claudia Renata dos Santos Barros; David Schlesinger; Debora Botequio Moretti; Dimas Tadeu Covas; Elaine Cristina Marqueze; Elaine Vieira Santos; Evandra Strazza Rodrigues; Heidge Fukumasu; Jayme Augusto de Souza-Neto; José Salvatore Leister Patané; Luiz Alcantara; Luiz Lehmann Coutinho; Maria Carolina Elias; Maurício Lacerda Nogueira; Rafael dos Santos Bezerra; Raul Machado Neto; Rejane Maria Tommasini Grotto; Ricardo Haddad; Sandra Coccuzzo Sampaio Vessoni; Simone Kashima; Svetoslav Nanev Slavov; Vincent Louis Viala |
| EPI_ISL_3864602 | Instituto Adolfo Lutz - Regional de Marilia | Instituto Adolfo Lutz, Interdiciplinary Procedures Center, Strategic Laboratory | Claudio Tavares Sacchi; Karoline Rodrigues Campos |
| EPI_ISL_3316305 | Instituto Adolfo Lutz - Regional de Ribeirao Preto | Instituto Adolfo Lutz, Interdiciplinary Procedures Center, Strategic Laboratory | Caio Vinicius Dias Lopes; Claudia Regina Gonçalves; Claudio Tavares Sacchi; Karoline Rodrigues Campos; Leonardo Tadeu de Araujo; Marlon Benedito Nascomento Santos |
| EPI_ISL_3545773 | Instituto Adolfo Lutz - Regional de Sao Jose do Rio Preto | Instituto Adolfo Lutz, Interdiciplinary Procedures Center, Strategic Laboratory | Caio Vinicius Dias Lopes; Claudia Regina Gonçalves; Claudio Tavares Sacchi; Karoline Rodrigues Campos; Leonardo Tadeu de Araujo; Marlon Benedito Nascomento Santos |
| EPI_ISL_2378677, EPI_ISL_3824318 | Instituto de Biotecnologia - UNESP-Botucatu-SP | Instituto de Biotecnologia - UNESP-Botucatu-SP | Cecília Artico Banho; Cíntia Bittar; Fábio Sossai Possebon; Guilherme Campos; Helena Lage Ferreira; Jorge A. Petroli Marchesi; João Pessoa Araújo Jr.; Leila Sabrina Ullmann; Lívia Sacchetto; Maisa C. Pereira Parra; Marília Moraes; Maurício L. Nogueira; Paula Rahal; Paulo Inacio da Costa |
| EPI_ISL_3835356 | LACEN/PE | WallauLab on behalf of Fiocruz COVID-19 Genomic Surveillance Network | Alexandre Freitas da Silva; Cassia Docena; Constância Flávia Junqueira Ayres; Filipe Zimmer Dezordi; Gabriel Luz Wallau; Gustavo Barbosa de Lima; Lais Ceschini Machado; Lilian Caroliny Amorim Silva; Marcelo Henrique dos Santos Paiva; Matheus Filgueira Bezerra; Sinval Pinto Brandão Filho |
| EPI_ISL_3246154, EPI_ISL_3246163, EPI_ISL_3246173, EPI_ISL_3246178, EPI_ISL_3505201, EPI_ISL_3505203, EPI_ISL_3941517 | | | |
| see above | LATE - Laboratório de Técnicas Especiais - | LATE - Laboratório de Técnicas Especiais - Hospital Israelita Albert Einstein | Alexandre Hideaki Takara; Ana Paula Moreira Salles; Anelisie da Silva Santos; Deyvid Amgarten; Erick Gustavo Dorlass; Fernanda de Mello Malta; João Renato Rebello Pinho; Marcio Anunciacao Menezes; Pedro Henrique Sebe Rodrigues; Raquel Riyuzo |

| | | | |
|---|---|---|---|
| | Hospital Israelita Albert Einstein | | |
| EPI_ISL_3459928 | LESP Quintana Roo | Instituto de Diagnostico y Referencia Epidemiologicos (INDRE) | Abril Rodriguez-Maldonado; Ariadna Medina-Benitez; Claudia Wong-Arambula; Ernesto Ramirez-Gonzalez.; Gisela Barrera-Badillo; Irma Lopez-Martinez; Joaquin Quiroz-Mercado; Lucia Hernandez-Rivas; Maribel Gonzalez-Villa; Natividad Cruz-Ortiz; Sergio Rangel-Guerrero; Tatiana Nunez-Garcia; Vanessa Rivero-Arredondo |
| EPI_ISL_3219660, EPI_ISL_3219670, EPI_ISL_3306182 | Labo Analyses Med | National Reference Center for Viruses of Respiratory Infections, Institut Pasteur, Paris | Angela Brisebarre; Camille Capel; Christophe Malabat; Corinne Maufrais; Etienne Simon-Lorière; Frédéric Lemoine; Hub de Bioinformatique et Biostatistique; Louise Lefrançois; Marion Barbet; Maud Vanpeene; Méline Bizard; OphéLie Said-Delattre; Sylvie Behillil; Sylvie Van der Werf; Vincent Enouf |
| EPI_ISL_3334816 | Laboratoires Reunis, 38 Rue Hiel, 6131 Junglinster, Luxembourg | Laboratoire national de sante, Microbiology, Microbial Genomics Platform | Anke Wienecke-Baldacchino; Bernard Weber; Catherine Ragimbeau; Elodie Solarino; Fatu Djabi; Jessica Tapp; Lise Pignon; Raoul Salmon; Tamir Abdelrahman; Virginie Jover |
| EPI_ISL_2691526, EPI_ISL_2837090, EPI_ISL_2837127, EPI_ISL_2837128, EPI_ISL_2837131, EPI_ISL_3245111, EPI_ISL_3245132, EPI_ISL_3245229, EPI_ISL_3245236, EPI_ISL_3245286, EPI_ISL_3245295, EPI_ISL_3245305, EPI_ISL_3615060, EPI_ISL_3615222, EPI_ISL_3615242, EPI_ISL_3615243, EPI_ISL_3920877, EPI_ISL_3920916, EPI_ISL_3920919, EPI_ISL_3921184, EPI_ISL_3921198, EPI_ISL_3921199, EPI_ISL_3921369 | | | |
| see above | Laboratorio Central Noel Nutels | Bioinformatics Laboratory / LNCC | Alessandra P Lamarca; Alexandra L Gerber; Amilcar Tanuri; Ana Paula de C Guimaraes; Ana Tereza R Vasconcelos; Andrea Cony Cavalcanti; Caio Luiz Pereira Ribeiro; Cassia Alves; Cintia Policarpo; Claudia Maria Braga de Mello; Cristiane Gomes da Silva; Diana Mariani; Douglas Terra Machado; Erica Ramos dos Santos Nascimento; Fernanda Leitao dos Santos; Flavio Dias da Silva; Gleidson da Silva de Oliveira; Leandro Magalhaes de Souza; Liliane Cavalcante; Luiz G P de Almeida; Marcio Henrique de Oliveira Garcia; Mario Sergio Ribeiro; Ricardo Jose Barbosa Salviano; Ronaldo da Silva F Jr; Silvia Carvalho |
| EPI_ISL_3802928 | Laboratorio Central de Saude Publica do Estado de Alagoas (LACEN/AL) | Laboratory of Respiratory Viruses and Measles, Oswaldo Cruz Institute, FIOCRUZ | Agatha Soares; Alice Sampaio Rocha; Ana Carolina Mendonca; Anderson Brandao Leite; Anna Carolina Paixao; Elisa Cavalcante Pereira; Fernando Motta; Ighor Arantes; Luciana Appolinario; Marilda Siqueira on behalf of the Fiocruz COVID-19 Genomic Surveillance Network; Paola Resende; Renata Serrano Lopes; Taina Venas |
| EPI_ISL_2983341, EPI_ISL_2983358 | Laboratorio Central de Saude Publica do Estado de Santa Catarina (LACEN-SC) | Laboratory of Respiratory Viruses and Measles, Oswaldo Cruz Institute, FIOCRUZ | Alice Sampaio Rocha; Ana Carolina Mendonca; Anna Carolina Paixao; Darcita Buerger Rovaris; Elisa Cavalcante Pereira; Fernando Motta; Luciana Appolinario; Marilda Siqueira on behalf of the Fiocruz COVID-19 Genomic Surveillance Network; Paola Resende; Renata Serrano Lopes; Sandra Bianchini Fernandes; Taina Venas |
| EPI_ISL_2983390, EPI_ISL_3190220, EPI_ISL_3235295, EPI_ISL_3235299, EPI_ISL_3434722, EPI_ISL_3434723, EPI_ISL_3434724, EPI_ISL_3434725, EPI_ISL_3434822, EPI_ISL_3434823, EPI_ISL_3434824, EPI_ISL_3434953, EPI_ISL_3434954, EPI_ISL_3434955, EPI_ISL_3827883, EPI_ISL_3827909, EPI_ISL_3827986, EPI_ISL_3828015, EPI_ISL_3828017 | | | |
| see above | Laboratorio Central de Saude Publica do Estado de Santa Catarina (LACEN/SC) | Laboratory of Respiratory Viruses and Measles, Oswaldo Cruz Institute, FIOCRUZ | Alice Sampaio Rocha; Ana Carolina Mendonca; Anna Carolina Paixao; Darcita Buerger Rovaris; Elisa Cavalcante Pereira; Fernando Motta; Luciana Appolinario; Marilda Siqueira on behalf of the Fiocruz COVID-19 Genomic Surveillance Network; Paola Resende; Renata Serrano Lopes; Sandra Bianchini Fernandes; Taina Venas |
| EPI_ISL_3539827 | Laboratorio Central de Saude Publica do Estado do Amapa (LACEN/AP) | Laboratory of Respiratory Viruses and Measles, Oswaldo Cruz Institute, FIOCRUZ | Agatha Cristinne Prudencio; Alice Sampaio Rocha; Ana Carolina Mendonca; Andreia Santos Costa; Anna Carolina Paixao; Anne Caroline da Silva Soledade; Elisa Cavalcante Pereira; Fernando Motta; Ighor Leonardo Arantes Gomes; Lindomar dos Anjos Silva; Luciana Appolinario; Marcia Socorro Pereira Cavalcante; Marilda Siqueira on behalf of the Fiocruz COVID-19 Genomic Surveillance Network; Paola Resende; Renata Serrano Lopes; Taina Venas |
| EPI_ISL_3045456, EPI_ISL_3434991 | Laboratorio Central de Saude Publica do Estado do Espirito Santo (LACEN/ES) | Laboratory of Respiratory Viruses and Measles, Oswaldo Cruz Institute, FIOCRUZ | Alice Sampaio Rocha; Ana Carolina Mendonca; Anna Carolina Paixao; Elisa Cavalcante Pereira; Fernando Motta; Luciana Appolinario; Marilda Siqueira on behalf of the Fiocruz COVID-19 Genomic Surveillance Network; Paola Resende; Renata Serrano Lopes; Rodrigo Ribeiro Rodrigues; Taina Venas |
| EPI_ISL_3761556, EPI_ISL_3761686, EPI_ISL_3761755 | Laboratorio de Pesquisa em Virologia, FAMERP, SJRP | Laboratorio de Pesquisa em Virologia, FAMERP, SJRP | Beatriz de Carvalho Marques; Cecília Artico Banho; Cíntia Bittar; Fábio Sossai Possebon; Guilherme Campos; Helena Lage Ferreira; Jorge A. Petroli Marchesi; João Pessoa Araújo Jr.; Leila Sabrina Ullmann; Lívia Sacchetto; Maisa C. Pereira Parra; Marília Moraes; Maurício L. Nogueira.; Paula Rahal; Paulo Inacio da Costa |
| EPI_ISL_2397913 | Laboratory Corporation of America | Centers for Disease Control and Prevention Division of Viral Diseases, Pathogen Discovery | Adrian Paskey; Amanda Douglas; Amanda Suchanek; Andrea Throop; Ayla Burns; Benjamin Rambo-Martin; Bobbi Croy; Brian Krueger; Brian Norvell; Christopher Gulvick; Christos Petropoulos; Clinton R. Paden; Craig Lukasik; Dakota Howard; Darlene Wagner; Debbie Boles; Dhwani Batra; Duncan MacCannell; Eyad Almasri; Goran Stevovic; Howard Engler; Hrushikesh Deshmukh; Jake Humphrey; Jana Schroth; Jason Caravas; Joe Voshell; John Pruitt; Jonathan Meltzer; Jonathan Williams; Kara Moser; Kimberly Wagner; Lax Iyer; Lisa Pfefferle; Lyndon Tilson; Manoj Jain; Marcia Eisenberg; Mary Ann Cristobal; Mary Williamson; Matthew Robinson; Matthew Schmerer; Michael Levandoski; Mike Sapeta; Mindy Nye; Minoo Agarwal; Mohan Kolli; Nuthawin Charoensri; Oren Cohen; Peter W. Cook; Prashant Gupta; Qian Zeng; Rama Ghatti; Scott Parker; Scott Ryan; Scott Sammons; Shatavia Morrison; Stanley Letovsky; Steven Ragan; Suresh Babu Selvaraju; Susan Countryman; Susan Hicks; Suzanne Dale; Thomas Urban; Tim Kuphal; Tricia Zwiefelhofer; Vincent Drouillon; Yvette Unoarumhi |
| EPI_ISL_2982741, EPI_ISL_3045541, EPI_ISL_3045542, EPI_ISL_3434952, EPI_ISL_3540000, EPI_ISL_3832445 | Laboratory of Respiratory Viruses and Measles, Oswaldo Cruz Institute, FIOCRUZ | Laboratory of Respiratory Viruses and Measles, Oswaldo Cruz Institute, FIOCRUZ | Agatha Cristinne Prudencio; Agatha Soares; Alice Sampaio Rocha; Ana Carolina Mendonca; Anna Carolina Paixao; Elisa Cavalcante Pereira; Fernando Motta; Ighor Arantes; Ighor Leonardo Arantes Gomes; Luciana Appolinario; Marilda Siqueira on behalf of the Fiocruz COVID-19 Genomic Surveillance Network; Paola Resende; Renata Serrano Lopes; Taina Venas |
| EPI_ISL_3050350 | Laboratório Central de Saúde Pública do Amazonas - LACEN-AM | Laboratorio de Ecologia de Doencas Transmissiveis na Amazonia, Instituto Leonidas e Maria Deane - Fiocruz Amazonia | André Corado; Felipe Naveca; Fernanda Nascimento; George Silva; Karina Pessoa; Luciana Gonçalves; Maria Júlia Brandão; Matilde Mejía; Valdinete Nascimento; Victor Souza; Ágatha Costa |
| EPI_ISL_3492502, EPI_ISL_3756578 | Laboratório de Microbiologia Molecular - Universidade FEEVALE | Molecular Microbiology Laboratory | Alana Witt Hansen; Fernando Rosado Spilki; Fágner Henrique Heldt; Juliana Schons Gulart; Juliana Schons Gularte; Juliane Deise Fleck; Mariana Soares da Silva; Matheus Nunes Weber; Meriane Demoliner; Michele Filippi.; Micheli Filippi; Paula Rodrigues de Almeida; Vyctoria Malayhka de Abreu Góes Pereira. |
| EPI_ISL_3190201 | Labortorio Central de Saude Publica do Estado do Rio de Janeiro (LACEN/RJ) | Laboratory of Respiratory Viruses and Measles, Oswaldo Cruz Institute, FIOCRUZ | Agatha Cristinne Prudencio Soares; Alice Sampaio Rocha; Ana Carolina Mendonca; Andrea Cony Cavalcanti; Anna Carolina Paixao; Elisa Cavalcante Pereira; Fernando Motta; Ighor Leonardo Arantes Gomes; Luciana Appolinario; Marilda Siqueira on behalf of the Fiocruz COVID-19 Genomic Surveillance Network; Paola Resende; Renata Serrano Lopes; Taina Venas |
| EPI_ISL_3536300 | MATERNIDADE ESCOLA ASSIS CHATEAUBRIAND | Oswaldo Cruz Institute, FIOCRUZ/CE | Cleber Furtado Aksenen; Fabio Miyajima; Fernando Braga Stehling; Francisco Eder de Moura Lopes; Jamille Maria Mendes Bezerra; Joaquim César do Nascimento Sousa Junior; Pedro Miguel Carneiro Jeronimo; Suzana Porto Almeida e Lucas Delerino; Thais Ferreira de Oliveira; Thais de Oliveira Costa; Ticiane Cavalcante de Souza; Veridiana Pessoa Miyajima |
| EPI_ISL_3875779 | Microbiology Department. Complexo Hospitalario Universitario de Vigo | Microbiology Department. Complexo Hospitalario Universitario de Vigo | Alfaya N; Alonso I; Alvarez M; Cabrera JJ; Carballo R; Cores O; Cortizo S; Davina C; Martinez L; Mediero G; Perez S; Potel C; Regueiro B; Rey S; Vasallo FJ; del-Campo V |
| EPI_ISL_2861081, EPI_ISL_2861082 | Mobile County Health Department | Synergy Laboratories | Megan Cornwell |
| EPI_ISL_3778602 | Pandemic Response Lab - NYC | Pandemic Response Lab, R&D | Alex Carpio; Cybill del Castillo; Dylan Law; Haiping Hao; Henry Lee; Isabel Fernandez Escapa; Jon Laurent; Melissa Hopkins; Michael Hammerling; Pradeep Bugga; Shinyoung Clair Kang; Sol Rey; William Ward |
| EPI_ISL_3291772, EPI_ISL_3959735 | Respiratory Virus Unit, | COVID-19 Genomics UK (COG-UK) Consortium | PHE Covid Sequencing Team |

| | | | |
|---|---|---|---|
| | Microbiology Services Colindale, Public Health England | | |
| EPI_ISL_2494122 | UBS DR ALFREDO DANTAS DE SOUZA UMUARAMA | Instituto Butantan | Antonio Jorge Martins; Claudia Renata dos Santos Barros; David Schlesinger; Debora Botequio Moretti; Dimas Tadeu Covas; Elaine Cristina Marqueze; Elaine Vieira Santos; Evandra Strazza Rodrigues; Heidge Fukumasu; Jayme Augusto de Souza-Neto; José Salvatore Leister Patané; Luiz Alcantara; Luiz Lehmann Coutinho; Maria Carolina Elias; Maurício Lacerda Nogueira; Rafael dos Santos Bezerra; Raul Machado Neto; Rejane Maria Tommasini Grotto; Ricardo Haddad; Sandra Coccuzzo Sampaio Vessoni; Simone Kashima; Svetoslav Nanev Slavov; Vincent Louis Viala |
| EPI_ISL_3579319 | UBS II DE ALVARES MACHADO | Instituto Butantan | Antonio Jorge Martins; Claudia Renata dos Santos Barros; David Schlesinger; Debora Botequio Moretti; Dimas Tadeu Covas; Elaine Cristina Marqueze; Elaine Vieira Santos; Evandra Strazza Rodrigues; Heidge Fukumasu; Jayme Augusto de Souza-Neto; José Salvatore Leister Patané; Luiz Alcantara; Luiz Lehmann Coutinho; Maria Carolina Elias; Maurício Lacerda Nogueira; Rafael dos Santos Bezerra; Raul Machado Neto; Rejane Maria Tommasini Grotto; Ricardo Haddad; Sandra Coccuzzo Sampaio Vessoni; Simone Kashima; Svetoslav Nanev Slavov; Vincent Louis Viala |
| EPI_ISL_3536296 | UNIDADE PRONTO ATENDIMENTO CONJUNTO CEARA | Oswaldo Cruz Institute, FIOCRUZ/CE | Cleber Furtado Aksenen; Fabio Miyajima; Fernando Braga Stehling; Francisco Eder de Moura Lopes; Jamille Maria Mendes Bezerra; Joaquim César do Nascimento Sousa Junior; Pedro Miguel Carneiro Jeronimo; Suzana Porto Almeida e Lucas Delerino; Thais Ferreira de Oliveira; Thais de Oliveira Costa; Ticiane Cavalcante de Souza; Veridiana Pessoa Miyajima |
| EPI_ISL_2462322, EPI_ISL_2494124 | UPA DE BEBEDOURO | Instituto Butantan | Antonio Jorge Martins; Claudia Renata dos Santos Barros; David Schlesinger; Debora Botequio Moretti; Dimas Tadeu Covas; Elaine Cristina Marqueze; Elaine Vieira Santos; Evandra Strazza Rodrigues; Heidge Fukumasu; Jayme Augusto de Souza-Neto; José Salvatore Leister Patané; Luiz Alcantara; Luiz Lehmann Coutinho; Maria Carolina Elias; Maurício Lacerda Nogueira; Rafael dos Santos Bezerra; Raul Machado Neto; Rejane Maria Tommasini Grotto; Ricardo Haddad; Sandra Coccuzzo Sampaio Vessoni; Simone Kashima; Svetoslav Nanev Slavov; Vincent Louis Viala |
| EPI_ISL_3579312 | USF PAULISTA FERNANDOPOLIS ANTONIO PIVATO | Instituto Butantan | Antonio Jorge Martins; Claudia Renata dos Santos Barros; David Schlesinger; Debora Botequio Moretti; Dimas Tadeu Covas; Elaine Cristina Marqueze; Elaine Vieira Santos; Evandra Strazza Rodrigues; Heidge Fukumasu; Jayme Augusto de Souza-Neto; José Salvatore Leister Patané; Luiz Alcantara; Luiz Lehmann Coutinho; Maria Carolina Elias; Maurício Lacerda Nogueira; Rafael dos Santos Bezerra; Raul Machado Neto; Rejane Maria Tommasini Grotto; Ricardo Haddad; Sandra Coccuzzo Sampaio Vessoni; Simone Kashima; Svetoslav Nanev Slavov; Vincent Louis Viala |
| EPI_ISL_2803308, EPI_ISL_3826062 | UW Virology Lab | UW Virology Lab | Alexander Greninger; Hong Xie; Keith R Jerome; Lasata Shrestha; Maria Lukes; Meei-Li Huang; Noah R. Baker; Pavitra Roychoudhury; Ricardo Perez; Savanna S. Carmack; Sean Ellis; Shah Mohamed Bakhash; Tien V. Nguyen |
| EPI_ISL_2942703, EPI_ISL_3805744 | Unidad de Investigacion Medica de Yucatan (UIMY) | Unidad de Genomica Avanzada | ; Alejandra Garcia-Gasca; Alejandra Hernandez-Teran; Alejandro Sanchez-Flores; Alfredo Herrera-Estrella; Alicia Ocaña-Mondragon; Andreu Comas-Garcia; Angel Gustavo Salas-Lais; Antonio Loza Roman; Bernardo Martinez-Miguel; Blanca Taboada; Brenda Irasema Maldonado-Meza; Bruno Gomez-Gil; Carla Ivon Herrera-Najera; Carlos F. Arias; Celia Boukadida; Celida Duque Molina; Celida Martinez- Rodriguez; Clara Esperanza Santacruz-Tinoco; Concepcion Grajales-Muñiz; Consorcio Mexicano de Vigilancia Genomica (CoViGen-Mex). Authors (in alphabetical order): Julio Elias Alvarado-Yaah; Cristobal Chaidez-Quiroz; Daniel Fregoso-Rueda; Daniel Lira Morales; Eduardo Becerril-Vargas; Fernando Fontove-Herrera; Fidencio Mejia-Nepomuceno; Francisco Pulido; Gloria Elena Espinosa-Ayala; Gloria Maria Molina-Salinas; Gloria Vazquez; Hector Esteban Paz-Juarez; Hector Montoya-Fuentes; Helen Haydee Fernanda Ramirez-Plascencia; Irvin Gonzalez-Lopez; Jean Pierre Gonzalez; Jesus Hernandez; Joel Armando Vazquez-Perez.; Jorge Salas-Hernandez; Jose Antonio Enciso-Moreno; Jose Arturo Martinez-Orozco; Jose Esteban Muñoz-Medina; Jose de Jesus Nuñez-Contreras; Juan Bautista Chale-Dzul; Julissa Enciso-Ibarra; Luis Alberto Ochoa-Carrera; Margarita Matias-Florentino; Maria Guadalupe Santiago-Mauricio; Maria Guadalupe de Jesus Mireles-Rivera; Mario Mujica-Sanchez; Marissa Perez-Garcia; Nelly Selem-Mojica; Pavel Isa; Ricardo Ciria Merce; Ricardo Grande; Rosa Maria Gutierrez Rios; Santiago avila-Rios; Selene Zarate; Susana Lopez; Veronica Mata-Haro; Victor Eduardo Garcia-Arias; Victor Hugo Borja-Aburto |
| EPI_ISL_2691654, EPI_ISL_2837286, EPI_ISL_3072157, EPI_ISL_3072164, EPI_ISL_3072165, EPI_ISL_3072195, EPI_ISL_3072256, EPI_ISL_3072260, EPI_ISL_3072303, EPI_ISL_3072304, EPI_ISL_3072305, EPI_ISL_3072351, EPI_ISL_3072405, EPI_ISL_3072407, EPI_ISL_3072413, EPI_ISL_3072523, EPI_ISL_3072533, EPI_ISL_3072561, EPI_ISL_3072616, EPI_ISL_3245355, EPI_ISL_3245374, EPI_ISL_3245387, EPI_ISL_3245390, EPI_ISL_3245419, EPI_ISL_3245424, EPI_ISL_3245435, EPI_ISL_3245440, EPI_ISL_3245464, EPI_ISL_3615209, EPI_ISL_3615244, EPI_ISL_3615245, EPI_ISL_3615246, EPI_ISL_3615247, EPI_ISL_3615248, EPI_ISL_3615249, EPI_ISL_3615250, EPI_ISL_3615251, EPI_ISL_3615252, EPI_ISL_3615253, EPI_ISL_3615254, EPI_ISL_3615255, EPI_ISL_3615256, EPI_ISL_3615257, EPI_ISL_3921397, EPI_ISL_3921448 | | | |
| see above | Unidade de apoio ao diagnostico da COVID - UNADIG | Bioinformatics Laboratory / LNCC | Alessandra P Lamarca; Alexandra L Gerber; Amilcar Tanuri; Ana Paula de C Guimaraes; Ana Tereza R Vasconcelos; Andrea Cony Cavalcanti; Caio Luiz Pereira Ribeiro; Cassia Alves; Cintia Policarpo; Claudia Maria Braga de Mello; Cristiane Gomes da Silva; Diana Mariani; Douglas Terra Machado; Erica Ramos dos Santos Nascimento; Fernanda Leitao dos Santos; Flavio Dias da Silva; Gleidson da Silva de Oliveira; Leandro Magalhaes de Souza; Liliane Cavalcante; Luiz G P de Almeida; Marcio Henrique de Oliveira Garcia; Mario Sergio Ribeiro; Ricardo Jose Barbosa Salviano; Ronaldo da Silva F Jr; Silvia Carvalho |
