## Supplementary material for "SARS-COV-2 γ variant acquires spike P681H or P681R for improved viral fitness": Acknowledgement table on the GISAID genomes used in this study

We gratefully acknowledge the following Authors from the Originating laboratories responsible for obtaining the specimens, as well as the Submitting laboratories where the genome data were generated and shared via GISAID, on which this research is based.

All Submitters of data may be contacted directly via www.gisaid.org

Authors are sorted alphabetically.

| Accession ID | Originating Laboratory | Submitting Laboratory | Authors |
|---|---|---|---|
| EPI_ISL_3116512, EPI_ISL_3116513, EPI_ISL_3117399, EPI_ISL_3117400 | Edmonton Provincial Lab | Public Health Agency of Canada (PHAC) National Microbiology Laboratory | Buss; Croxen M; Deo A; Dieu P; E; Ferrato C; Gill K; Khan F; Koleva P; Li V; Lloyd C; Lynch T; Ma R; Murphy S; Pabbaraju K; Shokoples S; Thayer J; Tipples G; Whitehouse M; Wong A; Yu C; Zelyas N |
| EPI_ISL_3518764 | Laboratory Corporation of America | Centers for Disease Control and Prevention Division of Viral Diseases, Pathogen Discovery | Adrian Paskey; Amanda Douglas; Amanda Suchanek; Andrea Throop; Ayla Burns; Benjamin Rambo-Martin; Bobbi Croy; Brian Krueger; Brian Norvell; Christopher Gulvick; Christos Petropoulos; Clinton Paden; Craig Lukasik; Dakota Howard; Darlene Wagner; Debbie Boles; Dhwani Batra; Duncan MacCannell; Eyad Almasri; Goran Stevovic; Howard Engler; Hrushikesh Deshmukh; Jake Humphrey; Jana Schroth; Jason Caravas; Joe Voshell; John Pruitt; Jonathan Meltzer; Jonathan Williams; Kara Moser; Kimberly Wagner; Lax Iyer; Lisa Pfefferle; Lyndon Tilson; Manoj Jain; Marcia Eisenberg; Mary Cristobal; Mary Williamson; Matthew Robinson; Matthew Schmerer; Michael Levandoski; Mike Sapeta; Mindy Nye; Minoo Agarwal; Mohan Kolli; Nuthawin Charoensri; Oren Cohen; Peter Cook; Prashant Gupta; Qian Zeng; Rama Ghatti; Scott Parker; Scott Ryan; Scott Sammons; Shatavia Morrison; Stanley Letovsky; Steven Ragan; Suresh Selvaraju; Susan Countryman; Susan Hicks; Suzanne Dale; Thomas Urban; Tim Kuphal; Tricia Zwiefelhofer; Vincent Drouillon; Yvette Unoarumhi |
| EPI_ISL_2872074 | Quest Diagnostics Incorporated | Centers for Disease Control and Prevention Division of Viral Diseases, Pathogen Discovery | A. Gerasimova; A. Perez; Adrian Paskey; B. Anderson; Benjamin Rambo-Martin; Christopher Gulvick; Clinton R. Paden; Dakota Howard; Darlene Wagner; Dhwani Batra; Duncan MacCannell; F. Lacbawan; I. A. Shlyakhter; Jason Caravas; K.E. Livingston; Kara Moser; L.E. Bernstein; M. Hua; Matthew Schmerer; P. Tanpaiboon; Peter W. Cook; R. M. Kagan; R. Owen; R. V. Rolando; S. H. Rosenthal; Scott Sammons; Shatavia Morrison; Y. Liu; Yvette Unoarumhi |
