## Supplementary material for "SARS-COV-2 γ variant acquires spike P681H or P681R for improved viral fitness": Acknowledgement table on the GISAID genomes used in this study

We gratefully acknowledge the following Authors from the Originating laboratories responsible for obtaining the specimens, as well as the Submitting laboratories where the genome data were generated and shared via GISAID, on which this research is based.

All Submitters of data may be contacted directly via www.gisaid.org

Authors are sorted alphabetically.

| Accession ID | Originating Laboratory | Submitting Laboratory | Authors |
|---|---|---|---|
| EPI_ISL_3922281, EPI_ISL_3922282 | AMBULATORIO DE ESPECIALIDADE V E MOGI MIRIM | Instituto Butantan | Antonio Jorge Martins; Claudia Renata dos Santos Barros; David Schlesinger; Debora Botequio Moretti; Dimas Tadeu Covas; Elaine Cristina Marqueze; Elaine Vieira Santos; Evandra Strazza Rodrigues; Heidge Fukumasu; Jayme Augusto de Souza-Neto; José Salvatore Leister Patané; Luiz Alcantara; Luiz Lehmann Coutinho; Maria Carolina Elias; Maurício Lacerda Nogueira; Rafael dos Santos Bezerra; Raul Machado Neto; Rejane Maria Tommasini Grotto; Ricardo Haddad; Sandra Coccuzzo Sampaio Vessoni; Simone Kashima; Svetoslav Nanev Slavov; Vincent Louis Viala |
| EPI_ISL_2716544, EPI_ISL_2716548, EPI_ISL_2716558, EPI_ISL_2716561, EPI_ISL_2716563, EPI_ISL_2812655, EPI_ISL_2812658, EPI_ISL_2812667, EPI_ISL_2812672, EPI_ISL_2812681, EPI_ISL_2812683, EPI_ISL_2861866, EPI_ISL_2861888, EPI_ISL_2861892, EPI_ISL_2861896, EPI_ISL_3062957, EPI_ISL_3063004, EPI_ISL_3063032, EPI_ISL_3063033, EPI_ISL_3063035, EPI_ISL_3235077 | | | |
| see above | ASL Napoli 1 Centro | AMES Centro Polidiagnostico Strumentale S.r.l. | "Giovanni Savarese; Antonella Di Carlo; Antonio Fico; Antonio Fico"; Eloisa Evangelista; Giovanni Savarese; Luigi D'Amore; Luisa Circelli; Maurizio D'Amora; Maurizio D'Amora Antonio Fico; Maurizio D'Amora Antonio Fico"; Monica Ianniello; Nadia Petrillo; Raffaella Ruggiero; Roberto Sirica |
| EPI_ISL_2963435, EPI_ISL_2963439 | AULSS 3 Venezia | Istituto Zooprofilattico Sperimentale delle Venezie | Adelaide Milani; Alessia Schivo; Alice Fusaro; Ambra Pastori; Annalisa Salviato; Antonia Ricci; Calogero Terregino; Edoardo Giussani; Elisa Palumbo; Erika Giorgia Quaranta; Isabella Monne; Luca Tassoni |
| EPI_ISL_3007277 | AULSS 3 Venezia | UOSD Genetica e Citogenetica - Azienda ULSS 3 Serenissima; Istituto Zooprofilattico Sperimentale delle Venezie | Adelaide Milani; Alessia Schivo; Alice Fusaro; Ambra Pastori; Annalisa Salviato; Antonia Ricci; Calogero Terregino; Claudia Perini; Edoardo Giussani; Elisa Palumbo; Elisa Squarcina; Erika Giorgia Quaranta; Isabella Monne; Laura Bevilacqua; Laura Squarzon; Luca Tassoni; Mosé Favarato |
| EPI_ISL_2687308, EPI_ISL_3218504, EPI_ISL_3220296, EPI_ISL_3221016, EPI_ISL_3321445, EPI_ISL_3322141, EPI_ISL_3430438, EPI_ISL_3817112, EPI_ISL_3817288, EPI_ISL_3817333, EPI_ISL_3819479, EPI_ISL_3820453, EPI_ISL_3822027, EPI_ISL_3846175 | | | |
| see above | Aegis Sciences Corporation | Centers for Disease Control and Prevention Division of Viral Diseases, Pathogen Discovery | Adrian Paskey; Alec Vest; Benjamin Rambo-Martin; Christopher Gulvick; Clinton Paden; Clinton R. Paden; Cyndi Clark; Dakota Howard; Darlene Wagner; Dhwani Batra; Dillon Nall; Duncan MacCannell; Ethan Sanders; Holly Houdeshell; Jason Caravas; Kara Moser; Matthew Hardison; Matthew Schmerer; Ola Kvalvaag; Patrick Campbell; Peter Cook; Peter W. Cook; Rob Case; Scott Sammons; Shatavia Morrison; Shaun Westlund; Vikramsinha Ghorpade; Yvette Unoarumhi |
| EPI_ISL_3261131, EPI_ISL_3261139, EPI_ISL_3261140, EPI_ISL_3261181, EPI_ISL_3931962 | Arcispedale Santa Maria Nuova, Autoimmunità, Allergologia e Biotecnologie Innovative | Istituto Zooprofilattico Sperimentale della Lombardia e dell'Emilia Romagna (IZSLER), Risk Analysis and Genomic Epidemiology Unit | Alessandro Zerbini; Erika Scaltriti; Ilaria Menozzi; Lucia Belloni; Marina Morganti; Stefania Croci; Stefano Pongolini |
| EPI_ISL_3089183 | Arizona State Public Health Laboratory | Arizona State Public Health Laboratory | Jessica Escobar; Katherine Fullerton; Linda Getsinger; Matthew Contursi; Nobuko Fukushima; Stacy White; Trung Huynh; Victor Waddell |
| EPI_ISL_2839122 | Arizona State University | Arizona State University | Efrem S. Lim; Joshua LaBaer; Joy M. Blain; LaRinda A. Holland; Matthew F. Smith; Nicholas J. Mellor; Peter T. Skidmore; Rabia Maqsood; Valerie Harris; Vel Murugan |
| EPI_ISL_3425330 | Arlon | Plateforme de testing Namuroise | Degosserie Jonathan; Demars Aurore; Denis Olivier; Lesly Nyinkeu Kemamen; Maschietto Céline; Mullier François; Nicolas Gilliard; Nobis Chloé; Otto Gaetan |
| EPI_ISL_2861280, EPI_ISL_3030468 | Azienda Ospedaliera Terni | Istituto Zooprofilattico Sperimentale dell'Abruzzo e Molise "G. Caporale" | Ancora M; Calistri P; Cammà C; Curini V; Delli Compagni E; Di Domenico M; Di Pasquale A; Lorusso A; Mangone I; Marcacci M; Palumbo M; Puglia I; Rinaldi A; Savini G; Scaccetti A; Scialabba S |
| EPI_ISL_2820998, EPI_ISL_2820999, EPI_ISL_2821000, EPI_ISL_2821002, EPI_ISL_2821004, EPI_ISL_2821005, EPI_ISL_2821006, EPI_ISL_3932425, EPI_ISL_3932426, EPI_ISL_3932427, EPI_ISL_3932428, EPI_ISL_3932429, EPI_ISL_3932430, EPI_ISL_3932431, EPI_ISL_3932432, EPI_ISL_3932433, EPI_ISL_3932434, EPI_ISL_3932435, EPI_ISL_3932436, EPI_ISL_3932437, EPI_ISL_3932439, EPI_ISL_3932440, EPI_ISL_3932441, EPI_ISL_3932442 | | | |
| see above | Azienda Ospedaliero - Universitaria di Modena Policlinico - Virologia e Microbiologia Molecolare | Istituto Zooprofilattico Sperimentale della Lombardia e dell'Emilia Romagna (IZSLER), Risk Analysis and Genomic Epidemiology Unit | Erika Scaltriti; Giulia Fregni Serpini; Ilaria Menozzi; Marina Morganti; Monica Pecorari; Stefano Pongolini; William Gennari |
| EPI_ISL_2680975 | Azienda Sanitaria dell'Alto Adige - Laboratorio Aziendale di Microbiologia e Virologia | Azienda Sanitaria dell'Alto Adige | Irene Bianconi |
| EPI_ISL_2550630, EPI_ISL_2550634, EPI_ISL_2550635, EPI_ISL_2550642, EPI_ISL_2550661, EPI_ISL_2550672, EPI_ISL_2550673, EPI_ISL_2550675 | | | |
| see above | Azienda Sanitaria dell'Alto Adige Laboratorio Aziendale di Microbiologia e Virologia | Istituto di Genomica Applicata | Davide Scaglione; Eleonora Paparelli; Elisa Masi; Elisabetta Giacobazzi; Elisabetta Pagani; Gabriele Magris; Irena Jurman; Irene Bianconi; Michele Morgante; Stefanie Wieser; Vera Vendramin |
| EPI_ISL_3127241, EPI_ISL_3134448, EPI_ISL_3431226, EPI_ISL_3492394, EPI_ISL_3945739 | Broad Institute Clinical Research Sequencing Platform | Infectious Disease Program, Broad Institute of Harvard and MIT | Adams, G.; B.L.; B.W.; Bauer, M.; Birren; Blumenstiel, B.; Brown, C.; Carter, A.; Chaluvadi, S.; D.J.; DeFelice, M.; DeRuff, K.; Dodge, S.; Gabriel, S.; Gallagher, G.; Gladden-Young, A.; Granger, B.; J.E.; K.J.; Lagerborg, K.; Larkin, K.; Lee, M.; Lemieux; Lennon, N.; Loreth, C.; Madoff, L.; McGovern, S.; Meldrim, J.; Normandin, E.; P.C.; Park; Pearlman, L.; Reilly, S.; Rudy, M.; Sabeti; Siddle; Smole, S.; Tomkins-Tinch, C.; Vicente, G.; and MacInnis |
| EPI_ISL_3099595 | CAP Manlleu | Banc de Sang i Teixits | Carlos Hobeich; Francisco Vidal; Irene Corrales; Lorena Ramírez; Maria Glòria Soria; Natália Comes; Nina Borràs; Noemí Gonzalez; Silvia Sauleda |
| EPI_ISL_3669587, EPI_ISL_3669590, EPI_ISL_3672204 | CDPH VBL | California Department of Public Health | Emily Smith on behalf of CDPH-COVIDNet and UCSD EXCITE lab |
| EPI_ISL_3536351 | CENTRO DE ATENDIMENTO PARA ENFRENTAMENTO AO COVID 19 | Oswaldo Cruz Institute, FIOCRUZ/CE | Cleber Furtado Aksenen; Fabio Miyajima; Fernando Braga Stehling; Francisco Eder de Moura Lopes; Jamille Maria Mendes Bezerra; Joaquim César do Nascimento Sousa Junior; Pedro Miguel Carneiro Jeronimo; Suzana Porto Almeida e Lucas Delerino; Thais Ferreira de Oliveira; Thais de Oliveira Costa; Ticiane Cavalcante de Souza; Veridiana Pessoa Miyajima |
| EPI_ISL_3758165, EPI_ISL_3758170, EPI_ISL_3758172, EPI_ISL_3758173, EPI_ISL_3758175, EPI_ISL_3758176, EPI_ISL_3758177, EPI_ISL_3758180, EPI_ISL_3758182, EPI_ISL_3758183, EPI_ISL_3758187, EPI_ISL_3758190, EPI_ISL_3825450, EPI_ISL_3825452, EPI_ISL_3825453, EPI_ISL_3825454, EPI_ISL_3825457, EPI_ISL_3825459, EPI_ISL_3825461, EPI_ISL_3825462, EPI_ISL_3825463, EPI_ISL_3825464, EPI_ISL_3825465, EPI_ISL_3825466, EPI_ISL_3825467, EPI_ISL_3825468, EPI_ISL_3825470, EPI_ISL_3825471 | | | |
| see above | CHECK-UP - Medicina e Diagnóstico - Matriz | HLAGYN - Laboratorio de Imunologia de Transplantes de Goias | Elaize Maria Gomes de Paula; Fernando Antonio Vinhal dos Santos; Frederico Rodrigues Vinhal; Gladstone Rodrigues da Cunha Filho; Kamila Oliveira Reis De Freitas; Lucas Carlos Gomes Pereira; Nubia Silva Araújo; Sabrina Sara Moreira Duarte |
| EPI_ISL_2894568 | CIAD Hermosillo | CIAD Hermosillo | ; Alejandra García-Gasca; Alejandra Hernández-Terán; Alejandro Sánchez-Flores; Alfredo Herrera-Estrella; Alicia Ocaña-Mondragón; Andreu Comas-García; Angel Gustavo Salas-Lais; Antonio Loza Román; Bernardo Martínez-Miguel; Blanca Taboada; Brenda Irasema Maldonado-Meza; Bruno Gómez-Gil; Carla Ivón Herrera-Najera; Carlos F. Arias; Celia Boukadida; Clara Esperanza Santacruz-Tinoco; Concepción Grajales-Muñiz; Consorcio Mexicano de Vigilancia Genómica (CoViGen-Mex). Authors (in alphabetical order): Julio Elias Alvarado-Yaah; Cristóbal Cháidez-Quiróz; Célida Duque Molina; Célida Martínez- Rodríguez; Daniel Fregoso-Rueda; Daniel Lira Morales; Eduardo Becerril-Vargas; Fernando Fontove-Herrera; Fidencio Mejía-Nepomuceno; Francisco Pulido; Gloria Elena Espinosa-Ayala; Gloria María Molina-Salinas; Gloria Vazquez; Hector Esteban Paz-Juárez; Hector Montoya-Fuentes; Helen Haydee Fernanda Ramírez-Plascencia; Irvin González-López; Jean Pierre González; Jesús Hernández; Joel Armando Vázquez-Pérez.; Jorge Salas-Hernández; José Antonio Enciso-Moreno; José Arturo Martínez-Orozco; José Esteban Muñoz-Medina; José de Jesús Nuñez-Contreras; Juan Bautista Chale-Dzul; Julissa Enciso-Ibarra; Luis Alberto Ochoa-Carrera; Margarita Matías-Florentino; Mario Mújica-Sánchez; Marissa Perez-Garcia; María Guadalupe Santiago-Mauricio; María Guadalupe de Jesús Mireles-Rivera; Nelly Sélem-Mojica; Pavel Isa; Ricardo Ciria Merce; Ricardo Grande; Rosa María Gutiérrez Rios; Santiago Ávila-Ríos; Selene Zárate; Susana Lopez; Verónica Mata-Haro; Victor Eduardo García-Arias; Víctor Hugo Borja-Aburto |
| EPI_ISL_3944578, EPI_ISL_3944593, EPI_ISL_3944628 | Central Public Health Laboratory - LACEN - Bahia, Salvador, Brazil | Central Public Health Laboratory - LACEN -Bahia, Salvador, Brazil | Arabela Leal; Felicidade Pereira; Gabriela Menezes; Jaqueline Gomes; Lenisa Dandara; Luciana Oliveira; Luiz Alcantara; Marcela Gómez; Marta Giovanetti; Stephane Tosta; Vagner Fonseca; Vanessa Nardy |
| EPI_ISL_3347531 | Centro de Investigación Biomédica del Noreste (CIBIN) | Instituto de Biotecnología de la UNAM | ; Alejandra García-Gasca; Alejandra Hernández-Terán; Alejandro Sánchez-Flores; Alfredo Herrera-Estrella; Alicia Ocaña-Mondragón; Andreu Comas-García; Angel Gustavo Salas-Lais; Antonio Loza Román; Bernardo Martínez-Miguel; Blanca Taboada; Brenda Irasema Maldonado-Meza; Bruno Gómez-Gil; Carla Ivón Herrera-Najera; Carlos F. Arias; Celia Boukadida; Clara Esperanza Santacruz-Tinoco; Concepción Grajales-Muñiz; Consorcio Mexicano de Vigilancia Genómica (CoViGen-Mex). Authors (in alphabetical order): Julio Elias Alvarado-Yaah; Cristóbal Cháidez-Quiróz; Célida Duque Molina; Célida Martínez- Rodríguez; Daniel Fregoso-Rueda; Daniel Lira Morales; Eduardo Becerril-Vargas; Fernando Fontove-Herrera; Fidencio Mejía-Nepomuceno; Francisco Pulido; Gloria Elena Espinosa-Ayala; Gloria María Molina-Salinas; Gloria Vazquez; Hector Esteban Paz-Juárez; Hector Montoya-Fuentes; Helen Haydee Fernanda Ramírez-Plascencia; Irvin González-López; Jean Pierre González; Jesús Hernández; Joel Armando Vázquez-Pérez).; Jorge Salas-Hernández; José Antonio Enciso-Moreno; José Arturo Martínez-Orozco; José Esteban Muñoz-Medina; José de Jesús Nuñez-Contreras; Juan Bautista Chale-Dzul; Julissa Enciso-Ibarra; Kathia Elizabeth Tapia-Diaz; Luis Alberto Ochoa-Carrera; Margarita Matías-Florentino; Mario Mújica-Sánchez; Marissa Perez-Garcia; María Guadalupe Santiago-Mauricio; María Guadalupe de Jesús Mireles-Rivera; Nelly Sélem-Mojica; Pavel Isa; Ricardo Ciria Merce; Ricardo Grande; Rosa María Gutiérrez Rios; Santiago Ávila-Ríos; Selene Zárate; Susana Lopez; Verónica Mata-Haro; Victor Eduardo García-Arias; Víctor Hugo Borja-Aburto |
| EPI_ISL_3242534 | Clinica INDISA | "Facultad de Ciencias de la Vida, UNAB" | "Claudio Meneses; Ariel Orellana"; Claudio Olmos; Daniel Leon; Dayan Sanhueza; Eduardo Castro; Gonzalo Campaña; Macarena Bastias; Paola Pidal; Ricardo Yusta; Sebastian Wolter; Susana Saez; Víctor Monreal; Waldo Diaz |
| EPI_ISL_3988532 | Colorado Department of Public Health and Environment | Colorado Department of Public Health and Environment | Alexandria Rossheim; Diana Ir; Emily A. Travanty; Laura Bankers; Mandy Waters; Michael Martin; Molly C. Hetherington-Rauth; Sarah Elizabeth Totten; Shannon R. Matzinger |
| EPI_ISL_3663275 | Communicable Disease Laboratory, Public Health Directorate | Communicable Disease Laboratory, Public Health Directorate | AlAbbas, Z.; AlHujairi, Z.; Altaif, Z.; Alwasti, H.; Marhoon, A.; Touq, M. |
| EPI_ISL_2930322, EPI_ISL_2930325, EPI_ISL_2930327, EPI_ISL_2930330, EPI_ISL_2930347, EPI_ISL_2930362, EPI_ISL_2930407, EPI_ISL_3014120, EPI_ISL_3014123, EPI_ISL_3014156, EPI_ISL_3014157, EPI_ISL_3014158, EPI_ISL_3014159, EPI_ISL_3014160, EPI_ISL_3014161, EPI_ISL_3014162, EPI_ISL_3014163, EPI_ISL_3014164, EPI_ISL_3014165, EPI_ISL_3014166, EPI_ISL_3014167, EPI_ISL_3014168, EPI_ISL_3014169, EPI_ISL_3014170, EPI_ISL_3014171, EPI_ISL_3014173, EPI_ISL_3014174, EPI_ISL_3014175, EPI_ISL_3092096, EPI_ISL_3092133, EPI_ISL_3092134, EPI_ISL_3092163, EPI_ISL_3092226, EPI_ISL_3092270, EPI_ISL_3092309, EPI_ISL_3092315, EPI_ISL_3092336, EPI_ISL_3092406, EPI_ISL_3092419, EPI_ISL_3092420, EPI_ISL_3092421, EPI_ISL_3092422, EPI_ISL_3092423, EPI_ISL_3092424, EPI_ISL_3092425, EPI_ISL_3092426, EPI_ISL_3092427, EPI_ISL_3092428, EPI_ISL_3092429, EPI_ISL_3092430 | | | |
| see above | Cotugno | TIGEM | Antonio Grimaldi Patrizia Annunziata Francesco Panariello Biancamaria Pierri Claudia Tiberio Teresa Giuliano Valentina Bouche Chiara Colantuono Maria Concetta Cuomo Denise Di Concilio Lucio Di Filippo Anna Manfredi Marcello Salvi Antonio Limone Luigi Atripaldi Pellegrino Cerino Andrea Ballabio Davide Cacchiarelli |
| EPI_ISL_3064663 | Curative Labs | Curative Labs | Elias L. Salfati; Eugenia Khorosheva; George Way; J.Cesar Ignacio-Espinoza; Janet Chen; Mikhail Hanewich-Hollatz; Nabjot Sandhu; Sophia Quasem; Vladimir Slepnev; Zhiyi Xie |

| | | | |
|---|---|---|---|
| EPI_ISL_3841240, EPI_ISL_3841242, EPI_ISL_3841260, EPI_ISL_3841261, EPI_ISL_3841262, EPI_ISL_3841263, EPI_ISL_3841264, EPI_ISL_3841270, EPI_ISL_3841271, EPI_ISL_3841272, EPI_ISL_3841276, EPI_ISL_3841277, EPI_ISL_3841278, EPI_ISL_3841288, EPI_ISL_3841289, EPI_ISL_3841293, EPI_ISL_3841295, EPI_ISL_3841296, EPI_ISL_3841297, EPI_ISL_3841298, EPI_ISL_3841299, EPI_ISL_3841306, EPI_ISL_3841307, EPI_ISL_3841308 | | | |
| see above | DNAGYN | LGBio (Laboratorio de Genetica & Biodiversidade) | Alex Honda Bernardes; Amanda Alves de Melo; Aparecido Divino da Cruz; Cintia Pelegrineti Targueta de Azevedo Brito; Daniela de Melo e Silva; Elisangela de Paula Silveira Lacerda; Juliana Santana de Curcio; Luiz Augusto Pereira; Marc Alexandre Duarte Gigonzac; Mariana Pires de Campos Telles; Ramilla dos Santos Braga; Renata de Oliveira Dias; Rhewter Nunes; Thais Cidália Vieira Gigonzac; Thais Guimarães Castro; Thays Millena Alves Pedroso |
| EPI_ISL_3692660, EPI_ISL_3692711 | DPHL | Delaware Public Health Lab | Rebecca Savage |
| EPI_ISL_3383052 | DYOMEDEA-LABORATOIRE DE LA SAUVEGARDE | CNR Virus des Infections Respiratoires - France SUD | Antonin Bal; Bruno Lina; Gregory Destras; Gwendolyne Burfin; Hadrien Regue; Laurence Josset; Martine Valette; Quentin Semanas |
| EPI_ISL_2896927 | Department of Bacteria, Parasites and Fungi, Statens Serum Institut, Copenhagen, Denmark | Statens Serum Institut Bioinformatics and Microbial Genomics | Danish Covid-19 Genome Consortium |
| EPI_ISL_3050671, EPI_ISL_3050700, EPI_ISL_3050704, EPI_ISL_3050720, EPI_ISL_3050723, EPI_ISL_3372573, EPI_ISL_3372577, EPI_ISL_3372580, EPI_ISL_3372584, EPI_ISL_3372586, EPI_ISL_3372593, EPI_ISL_3372600 | | | |
| see above | Dirección regional de salud del Callao (DIRESA-CALLAO) | Centro de Investigaciones Tecnológicas, Biomédicas y Medioambientales (CITBM) | B; Huaman; J. Alarcon; M. Cuellar; M. Ramirez; M. Sovero |
| EPI_ISL_2672729, EPI_ISL_2981895 | Dutch COVID-19 response team | National Institute for Public Health and the Environment (RIVM) | Adam Meijer; AnneMarie van den Brandt; Annelies Kroneman; Bas van der Veer; Chantal Reusken; Dennis Schmitz; Dirk Eggink; Euníce Then; Florian Zwagemaker; Harry Vennema; Jeroen Cremer; Karim Hajji; Kim Freriks; Linda van de Nes; Lisa Wijsman; Lynn Aarts; Melissa van Tuil; Ryanne Jaarsma; Sanne Bos; Sharon van den Brink; on behalf of the national COVID-19 response team |
| EPI_ISL_3215815 | EXCITE Lab | Andersen lab at Scripps Research | Abigail Schnapper; Angela Scioscia; Chip Schooley; David Pride; Helena Tubb; Natasha Martin Cheryl Anderson; Sawyer Farmer; Sharon Reed; Tommy Valles + SEARCH |
| EPI_ISL_2728615 | Fondazione IRCCS Ca' Granda Ospedale Maggiore Policlinico | Fondazione IRCCS Ca' Granda Ospedale Maggiore Policlinico | Ferruccio Ceriotti; Sara Uceda Renteria |
| EPI_ISL_3299857, EPI_ISL_3299968, EPI_ISL_3344295, EPI_ISL_3349409, EPI_ISL_3350480, EPI_ISL_3351015, EPI_ISL_3352210, EPI_ISL_3429925, EPI_ISL_3429926, EPI_ISL_3429928, EPI_ISL_3430049, EPI_ISL_3456680, EPI_ISL_3456741, EPI_ISL_3456742, EPI_ISL_3520980, EPI_ISL_3521628, EPI_ISL_3608399, EPI_ISL_3659729, EPI_ISL_3661964, EPI_ISL_3663150 | | | |
| see above | Fulgent Genetics | Centers for Disease Control and Prevention Division of Viral Diseases, Pathogen Discovery | Adrian Paskey; Becky Tsai; Benafsh Sapra; Benjamin Rambo-Martin; Christopher Gulvick; Clinton Paden; Clinton R. Paden; Dakota Howard; Darlene Wagner; Dhwani Batra; Doreen Ng; Duncan MacCannell; Harry Gao; James Xie; Jason Caravas; John Gao; Joseph Fierro; Kara Moser; Matthew Schmerer; Mickey Li; Peter Cook; Peter W. Cook; Scott Sammons; Shatavia Morrison; Yan Meng; Yvette Unoarumhi |
| EPI_ISL_3099596, EPI_ISL_3320374, EPI_ISL_3386405, EPI_ISL_3386438 | Fundacio Althaia-Manresa | Banc de Sang i Teixits | Carlos Hobeich; Francisco Vidal; Irene Corrales; Lorena Ramírez; Maria Glòria Soria; Natália Comes; Nina Borràs; Noemí Gonzalez; Silvia Sauleda |
| EPI_ISL_3840802 | GA Department of Public Health | GA Department of Public Health | Aliyah Fields; Cynthia Dixey; Jonathan Edwards; Sharmila Talekar; Stacy Reeves; Taylor Smith; Tonia Parrott |
| EPI_ISL_2659183, EPI_ISL_2892829, EPI_ISL_3185595, EPI_ISL_3987698 | Genetica Molecular and Subdepartamento de Virologia ISP Chile | Instituto de Salud Publica de Chile | Andres Castillo; Barbara Parra; Constanza Campano; Gisselle Barra; Javier Tognarelli; Jorge Fernandez; Karen Orostica; Loredana Arata; Patricia Bustos; Rodrigo Fasce; Soledad Ulloa |
| EPI_ISL_3536215 | HGCC HOSPITAL GERAL DR CESAR CALS | Oswaldo Cruz Institute, FIOCRUZ/CE | Cleber Furtado Aksenen; Fabio Miyajima; Fernando Braga Stehling; Francisco Eder de Moura Lopes; Jamille Maria Mendes Bezerra; Joaquim César do Nascimento Sousa Junior; Pedro Miguel Carneiro Jeronimo; Suzana Porto Almeida e Lucas Delerino; Thais Ferreira de Oliveira; Thais de Oliveira Costa; Ticiane Cavalcante de Souza; Veridiana Pessoa Miyajima |
| EPI_ISL_2617597, EPI_ISL_2617598, EPI_ISL_2617599, EPI_ISL_2617600, EPI_ISL_2617601, EPI_ISL_2617602, EPI_ISL_2617603, EPI_ISL_2617604, EPI_ISL_2617605, EPI_ISL_2617606, EPI_ISL_2617613, EPI_ISL_2617614, EPI_ISL_2617615, EPI_ISL_2617616, EPI_ISL_2617617, EPI_ISL_2617618, EPI_ISL_2617619, EPI_ISL_2617620, EPI_ISL_2617621, EPI_ISL_2617622, EPI_ISL_2617623, EPI_ISL_2617624, EPI_ISL_2617625, EPI_ISL_2680908, EPI_ISL_2680911, EPI_ISL_2921536, EPI_ISL_2921537, EPI_ISL_2921546, EPI_ISL_2921548, EPI_ISL_2921549, EPI_ISL_2921551, EPI_ISL_2921553, EPI_ISL_2921558, EPI_ISL_2921561, EPI_ISL_2921562, EPI_ISL_2921563, EPI_ISL_2921575, EPI_ISL_2921579, EPI_ISL_2921581, EPI_ISL_2921582, EPI_ISL_2921585, EPI_ISL_2921593, EPI_ISL_2921596, EPI_ISL_2921600, EPI_ISL_2921601, EPI_ISL_2921607, EPI_ISL_3087897, EPI_ISL_3254317, EPI_ISL_3254318, EPI_ISL_3254329, EPI_ISL_3254371, EPI_ISL_3254376, EPI_ISL_3254377, EPI_ISL_3254385, EPI_ISL_3254389, EPI_ISL_3254394, EPI_ISL_3254399, EPI_ISL_3254415, EPI_ISL_3254417, EPI_ISL_3254420, EPI_ISL_3254430, EPI_ISL_3254432, EPI_ISL_3254436, EPI_ISL_3254437, EPI_ISL_3254444, EPI_ISL_3254456, EPI_ISL_3254457, EPI_ISL_3254474, EPI_ISL_3254478, EPI_ISL_3254483, EPI_ISL_3254484, EPI_ISL_3254490, EPI_ISL_3254497, EPI_ISL_3254524, EPI_ISL_3254525, EPI_ISL_3254529, EPI_ISL_3254534, EPI_ISL_3254537, EPI_ISL_3254540, EPI_ISL_3254550, EPI_ISL_3254552, EPI_ISL_3254555, EPI_ISL_3254557, EPI_ISL_3254559, EPI_ISL_3254566, EPI_ISL_3254567, EPI_ISL_3254570, EPI_ISL_3254588, EPI_ISL_3254591, EPI_ISL_3254592, EPI_ISL_3254593, EPI_ISL_3254599, EPI_ISL_3254600, EPI_ISL_3254607, EPI_ISL_3254609, EPI_ISL_3254616, EPI_ISL_3254631, EPI_ISL_3254638, EPI_ISL_3254639, EPI_ISL_3254644, EPI_ISL_3254646, EPI_ISL_3254648, EPI_ISL_3254649, EPI_ISL_3254650, EPI_ISL_3254655, EPI_ISL_3254658, EPI_ISL_3254661, EPI_ISL_3254667, EPI_ISL_3254669, EPI_ISL_3254670, EPI_ISL_3254671, EPI_ISL_3254674, EPI_ISL_3254678, EPI_ISL_3254680, EPI_ISL_3254681, EPI_ISL_3254683, EPI_ISL_3259352, EPI_ISL_3259355, EPI_ISL_3259357, EPI_ISL_3259360, EPI_ISL_3259361, EPI_ISL_3259367, EPI_ISL_3259369, EPI_ISL_3259370, EPI_ISL_3259373, EPI_ISL_3259375, EPI_ISL_3274744, EPI_ISL_3274751, EPI_ISL_3274752, EPI_ISL_3274754, EPI_ISL_3274756, EPI_ISL_3274758, EPI_ISL_3274759, EPI_ISL_3274760, EPI_ISL_3274765, EPI_ISL_3274766, EPI_ISL_3274771, EPI_ISL_3274772, EPI_ISL_3274775, EPI_ISL_3274776, EPI_ISL_3274778, EPI_ISL_3274779, EPI_ISL_3274780, EPI_ISL_3274782, EPI_ISL_3274785, EPI_ISL_3299425, EPI_ISL_3299427, EPI_ISL_3386128, EPI_ISL_3386129, EPI_ISL_3386130, EPI_ISL_3386131, EPI_ISL_3386133, EPI_ISL_3386134, EPI_ISL_3386135, EPI_ISL_3386136, EPI_ISL_3386139, EPI_ISL_3386140, EPI_ISL_3386144, EPI_ISL_3386145, EPI_ISL_3386146, EPI_ISL_3386147, EPI_ISL_3386148, EPI_ISL_3386150, EPI_ISL_3536437, EPI_ISL_3536440, EPI_ISL_3536444, EPI_ISL_3536447, EPI_ISL_3536448, EPI_ISL_3536449, EPI_ISL_3536450, EPI_ISL_3536451, EPI_ISL_3536454, EPI_ISL_3536455, EPI_ISL_3536456, EPI_ISL_3536457, EPI_ISL_3536460, EPI_ISL_3536461, EPI_ISL_3536463, EPI_ISL_3536464, EPI_ISL_3536465, EPI_ISL_3536467, EPI_ISL_3536468, EPI_ISL_3536469, EPI_ISL_3536471, EPI_ISL_3536473, EPI_ISL_3536475, EPI_ISL_3536476, EPI_ISL_3536477, EPI_ISL_3536481, EPI_ISL_3536486, EPI_ISL_3536487, EPI_ISL_3536491, EPI_ISL_3536495, EPI_ISL_3536497, EPI_ISL_3536499, EPI_ISL_3536500, EPI_ISL_3536505, EPI_ISL_3536506, EPI_ISL_3536509, EPI_ISL_3536510, EPI_ISL_3536513, EPI_ISL_3536514, EPI_ISL_3536515, EPI_ISL_3536517, EPI_ISL_3536523, EPI_ISL_3536524, EPI_ISL_3536525, EPI_ISL_3536526, EPI_ISL_3536528, EPI_ISL_3536529, EPI_ISL_3536530, EPI_ISL_3536532, EPI_ISL_3536533, EPI_ISL_3536536, EPI_ISL_3536538, EPI_ISL_3536539, EPI_ISL_3536540, EPI_ISL_3758059, EPI_ISL_3758061, EPI_ISL_3758062, EPI_ISL_3758063, EPI_ISL_3758064, EPI_ISL_3758067, EPI_ISL_3758068, EPI_ISL_3758070, EPI_ISL_3758071, EPI_ISL_3758072, EPI_ISL_3758073, EPI_ISL_3758075, EPI_ISL_3758078, EPI_ISL_3758079, EPI_ISL_3758081, EPI_ISL_3758082, EPI_ISL_3758083, EPI_ISL_3758084, EPI_ISL_3758088, EPI_ISL_3758089, EPI_ISL_3758090, EPI_ISL_3758091, EPI_ISL_3758094, EPI_ISL_3758095, EPI_ISL_3758096, EPI_ISL_3758098, EPI_ISL_3758099, EPI_ISL_3758101, EPI_ISL_3758102, EPI_ISL_3758103, EPI_ISL_3758104, EPI_ISL_3758105, EPI_ISL_3758106, EPI_ISL_3758107, EPI_ISL_3758108, EPI_ISL_3758112, EPI_ISL_3758113, EPI_ISL_3758114, EPI_ISL_3758117, EPI_ISL_3758118, EPI_ISL_3758120, EPI_ISL_3758121, EPI_ISL_3758122, EPI_ISL_3758125, EPI_ISL_3758126, EPI_ISL_3758129, EPI_ISL_3758131, EPI_ISL_3758132, EPI_ISL_3758137, EPI_ISL_3758139, EPI_ISL_3758140, EPI_ISL_3758143, EPI_ISL_3758147, EPI_ISL_3758148, EPI_ISL_3758150, EPI_ISL_3758151, EPI_ISL_3758152, EPI_ISL_3758153, EPI_ISL_3758155, EPI_ISL_3758157, EPI_ISL_3758158, EPI_ISL_3758160, EPI_ISL_3758161, EPI_ISL_3758162, EPI_ISL_3758163, EPI_ISL_3758164, EPI_ISL_3922201, EPI_ISL_3922203, EPI_ISL_3922204, EPI_ISL_3922207, EPI_ISL_3922208, EPI_ISL_3922209, EPI_ISL_3922211, EPI_ISL_3922216, EPI_ISL_3922217, EPI_ISL_3922219, EPI_ISL_3922220, EPI_ISL_3922222, EPI_ISL_3922224, EPI_ISL_3922226, EPI_ISL_3922227, EPI_ISL_3922228, EPI_ISL_3922233, EPI_ISL_3922235, EPI_ISL_3922239, EPI_ISL_3922240, EPI_ISL_3922241, EPI_ISL_3922243, EPI_ISL_3922244, EPI_ISL_3922245, EPI_ISL_3922246, EPI_ISL_3922247, EPI_ISL_3922248, EPI_ISL_3922249, EPI_ISL_3922251, EPI_ISL_3922253, EPI_ISL_3922255, EPI_ISL_3922259, EPI_ISL_3997109, EPI_ISL_3997112, EPI_ISL_3997114, EPI_ISL_3997116, EPI_ISL_3997125, EPI_ISL_3997127, EPI_ISL_3997131, EPI_ISL_3997133, EPI_ISL_3997138, EPI_ISL_3997140, EPI_ISL_3997142, EPI_ISL_3997145, EPI_ISL_3997148, EPI_ISL_3997150, EPI_ISL_3997152, EPI_ISL_3997156, EPI_ISL_3997158, EPI_ISL_3997159, EPI_ISL_3997161, EPI_ISL_3997162, EPI_ISL_3997165, EPI_ISL_3997166, EPI_ISL_3997167, EPI_ISL_3997168, EPI_ISL_3997169, EPI_ISL_3997170, EPI_ISL_3997171, EPI_ISL_3997173, EPI_ISL_3997174, EPI_ISL_3997175, EPI_ISL_3997176, EPI_ISL_3997177, EPI_ISL_3997178, EPI_ISL_3997179, EPI_ISL_3997181, EPI_ISL_3997182, EPI_ISL_3997184, EPI_ISL_3997186, EPI_ISL_3997187, EPI_ISL_3997188, EPI_ISL_3997189, EPI_ISL_3997191, EPI_ISL_3997192, EPI_ISL_3997193, EPI_ISL_3997194, EPI_ISL_3997196, EPI_ISL_3997197, EPI_ISL_3997198, EPI_ISL_3997201, EPI_ISL_3997202, EPI_ISL_3997203, EPI_ISL_3997204, EPI_ISL_3997205, EPI_ISL_3997207, EPI_ISL_3997215, EPI_ISL_3997217, EPI_ISL_3997220, EPI_ISL_3997223, EPI_ISL_3997224, EPI_ISL_3997226 | | | |
| see above | HLAGYN - Laboratorio de Imunologia de Transplantes de Goias | HLAGYN - Laboratorio de Imunologia de Transplantes de Goias | Alessandro Leonardo Alvares Magalhaes; Daniel Ferreira de Sousa; Danielle de Paiva Rezende; Erika Lopes Rocha Batista; Fernando Antonio Vinhal dos Santos; Frederico Rodrigues Vinhal; Kamila Oliveira Reis De Freitas.; Lucas Carlos Gomes Pereira; Sabrina Sara Moreira Duarte |
| EPI_ISL_3536226, EPI_ISL_3536234, EPI_ISL_3536246, EPI_ISL_3536253, EPI_ISL_3536255, EPI_ISL_3536256 | HM HOSPITAL DE MESSEJANA DR CARLOS ALBERTO STUDART GOMES | Oswaldo Cruz Institute, FIOCRUZ/CE | Cleber Furtado Aksenen; Fabio Miyajima; Fernando Braga Stehling; Francisco Eder de Moura Lopes; Jamille Maria Mendes Bezerra; Joaquim César do Nascimento Sousa Junior; Pedro Miguel Carneiro Jeronimo; Suzana Porto Almeida e Lucas Delerino; Thais Ferreira de Oliveira; Thais de Oliveira Costa; Ticiane Cavalcante de Souza; Veridiana Pessoa Miyajima |
| EPI_ISL_3221316 | HOSPITAL CAN MISSES | HOSPITAL UNIVERSITARIO SON ESPASES | Dr. Antonio Oliver; Dr. Carla López-Causapé; Dr. Gabriel Cabot; Hospital Universitario Son Espases; on behalf of Servicio de Microbiología |
| EPI_ISL_3912200, EPI_ISL_3912201, EPI_ISL_3912204 | HOSPITAL ESTADUAL LEONARDO DA VINCI | ACME Lab, Oswaldo Cruz Foundation, FIOCRUZ/CE | Cleber Furtado Aksenen; Fabio Miyajima; Fernando Braga Stehling; Francisco Eder de Moura Lopes; Jamille Maria Mendes Bezerra; Joaquim Cesar do Nascimento Sousa Junior; Pedro Miguel Carneiro Jeronimo; Suzana Porto Almeida & Lucas Delerino on behalf of COVID-19 FIOCRUZ Genomic Network; Thais Ferreira de Oliveira; Thais de Oliveira Costa; Ticiane Cavalcante de Souza; Veridiana Pessoa Miyajima |
| EPI_ISL_3536168, EPI_ISL_3536185, EPI_ISL_3536189, EPI_ISL_3536269, EPI_ISL_3536322 | HOSPITAL ESTADUAL LEONARDO DA VINCI | Oswaldo Cruz Institute, FIOCRUZ/CE | Cleber Furtado Aksenen; Fabio Miyajima; Fernando Braga Stehling; Francisco Eder de Moura Lopes; Jamille Maria Mendes Bezerra; Joaquim César do Nascimento Sousa Junior; Pedro Miguel Carneiro Jeronimo; Suzana Porto Almeida e Lucas Delerino; Thais Ferreira de Oliveira; Thais de Oliveira Costa; Ticiane Cavalcante de Souza; Veridiana Pessoa Miyajima |
| EPI_ISL_3912211 | HOSPITAL MATERNIDADE AGENOR ARAUJO | ACME Lab, Oswaldo Cruz Foundation, FIOCRUZ/CE | Cleber Furtado Aksenen; Fabio Miyajima; Fernando Braga Stehling; Francisco Eder de Moura Lopes; Jamille Maria Mendes Bezerra; Joaquim Cesar do Nascimento Sousa Junior; Pedro Miguel Carneiro Jeronimo; Suzana Porto Almeida & Lucas Delerino on behalf of COVID-19 FIOCRUZ Genomic Network; Thais Ferreira de Oliveira; Thais de Oliveira Costa; Ticiane Cavalcante de Souza; Veridiana Pessoa Miyajima |
| EPI_ISL_3912227 | HOSPITAL SAO JOSE DE DOENCAS INFECCIOSAS | ACME Lab, Oswaldo Cruz Foundation, FIOCRUZ/CE | Cleber Furtado Aksenen; Fabio Miyajima; Fernando Braga Stehling; Francisco Eder de Moura Lopes; Jamille Maria Mendes Bezerra; Joaquim Cesar do Nascimento Sousa Junior; Pedro Miguel Carneiro Jeronimo; Suzana Porto Almeida & Lucas Delerino on behalf of COVID-19 FIOCRUZ Genomic Network; Thais Ferreira de Oliveira; Thais de Oliveira Costa; Ticiane Cavalcante de Souza; Veridiana Pessoa Miyajima |
| EPI_ISL_3620498 | HOSPITAL UNIVERSITARI DE BELLVITGE | Microbiology Department | Aida Gonzalez-Diaz; Carmen Ardanuy; Jordi Camara; Jordi Niubó; Laura Calatayud; M Angeles Domínguez; Sara Marti; Yolanda Hernandez |
| EPI_ISL_3221262 | HOSPITAL UNIVERSITARIO SON ESPASES | HOSPITAL UNIVERSITARIO SON ESPASES | Dr. Antonio Oliver; Dr. Carla López-Causapé; Dr. Gabriel Cabot; Hospital Universitario Son Espases; on behalf of Servicio de Microbiología |
| EPI_ISL_3912229 | HPP LUIZ ROBERTO PESSOA AIRES | ACME Lab, Oswaldo Cruz Foundation, FIOCRUZ/CE | Cleber Furtado Aksenen; Fabio Miyajima; Fernando Braga Stehling; Francisco Eder de Moura Lopes; Jamille Maria Mendes Bezerra; Joaquim Cesar do Nascimento Sousa Junior; Pedro Miguel Carneiro Jeronimo; Suzana Porto Almeida & Lucas Delerino on behalf of COVID-19 FIOCRUZ Genomic Network; Thais Ferreira de Oliveira; Thais de Oliveira Costa; Ticiane Cavalcante de Souza; Veridiana Pessoa Miyajima |
| EPI_ISL_3536367 | HPP LUIZ ROBERTO PESSOA AIRES | Oswaldo Cruz Institute, FIOCRUZ/CE | Cleber Furtado Aksenen; Fabio Miyajima; Fernando Braga Stehling; Francisco Eder de Moura Lopes; Jamille Maria Mendes Bezerra; Joaquim César do Nascimento Sousa Junior; Pedro Miguel Carneiro Jeronimo; Suzana Porto Almeida e Lucas Delerino; Thais Ferreira de Oliveira; Thais de Oliveira Costa; Ticiane Cavalcante de Souza; Veridiana Pessoa Miyajima |
| EPI_ISL_3334590 | Hospital Center Emile Mayrisch | Laboratoire national de sante, Microbiology, Microbial Genomics Platform | Anke Wienecke-Baldacchino; Catherine Ragimbeau; Cynthia Oxacelay; Elodie Solarino; Fatu Djabi; Jessica Tapp; Lise Pignon; Raoul Salmon; Tamir Abdelrahman; Virginie Jover |
| EPI_ISL_3147603 | Hospital Center Luxembourg | Laboratoire national de sante, Microbiology, Microbial Genomics Platform | Anke Wienecke-Baldacchino; Catherine Ragimbeau; Elodie Solarino; Fatu Djabi; Jean-Hugues Francois; Jessica Tapp; Lise Pignon; Michel Kohnen; Raoul Salmon; Tamir Abdelrahman; Virginie Jover |
| EPI_ISL_2894554 | Hospital Margarita Maza de Juárez | Microbial Genomics Laboratory | ; Alejandra García-Gasca; Alejandra Hernández-Terán; Alejandro Sánchez-Flores; Alfredo Herrera-Estrella; Alicia Ocaña-Mondragón; Andreu Comas-García; Angel Gustavo Salas-Lais; Antonio Loza Román; Bernardo Martínez-Miguel; Blanca Taboada; Brenda Irasema Maldonado-Meza; Bruno Gómez-Gil; Carla Ivón Herrera-Najera; Carlos F. Arias; Celia Boukadida; Clara Esperanza Santacruz-Tinoco; Concepción Grajales-Muñiz; Consorcio Mexicano de Vigilancia Genómica (CoViGen-Mex). Authors (in alphabetical order): Julio Elias Alvarado-Yaah; Cristóbal Cháidez-Quiróz; Célida Duque Molina; Célida Martínez- Rodríguez; Daniel Fregoso-Rueda; Daniel Lira Morales; Eduardo Becerril-Vargas; Fernando Fontove-Herrera; Fidencio Mejía-Nepomuceno; Francisco Pulido; Gloria Elena Espinosa-Ayala; Gloria María Molina-Salinas; Gloria Vazquez; Hector Esteban Paz-Juárez; Hector Montoya-Fuentes; Helen Haydee Fernanda Ramírez-Plascencia; Irvin González-López; Jean Pierre González; Jesús Hernández; Joel Armando Vázquez-Pérez.; Jorge Salas-Hernández; José Antonio Enciso-Moreno; José Arturo Martínez-Orozco; José Esteban Muñoz-Medina; José de Jesús Nuñez-Contreras; Juan Bautista Chale-Dzul; Julissa Enciso-Ibarra; Luis Alberto Ochoa-Carrera; Margarita Matías-Florentino; Mario Mújica-Sánchez; Marissa Perez-Garcia; María Guadalupe Santiago-Mauricio; María Guadalupe de Jesús Mireles-Rivera; Nelly Sélem-Mojica; Pavel Isa; Ricardo Ciria Merce; Ricardo Grande; Rosa María Gutiérrez Rios; Santiago Ávila-Ríos; Selene Zárate; Susana Lopez; Verónica Mata-Haro; Victor Eduardo García-Arias; Víctor Hugo Borja-Aburto |
| EPI_ISL_3374776, EPI_ISL_3983948, EPI_ISL_3983950, EPI_ISL_3983951, EPI_ISL_3983952, EPI_ISL_3983953, EPI_ISL_3983954, EPI_ISL_3983955, EPI_ISL_3983956, EPI_ISL_3983957, EPI_ISL_3983989, EPI_ISL_3983993 | | | |

| | | | |
|---|---|---|---|
| see above | Hospital Universitari Arnau de Vilanova | Hospital Universitari Vall d'Hebron - Vall d'Hebron Institut de Recerca | Alejandra González-Sánchez; Andrés Antón; Ariadna Rando; Carla Castillo; Cristina Andrés; Damir Garcia-Cehic; Josep Quer; Juliana Esperalba; Karen García; Maria Carmen Martin; Maria Gema Codina; Maria Piñana; Rodrigo Vásquez; Tomàs Pumarola |
| EPI_ISL_3030597 | Hospital Universitari Bellvitge | Microbiology Department | Aida Gonzalez-Diaz; Carmen Ardanuy; Jordi Camara; Jordi Niubó; Laura Calatayud; M Angeles Domínguez; Sara Marti |
| EPI_ISL_2828109, EPI_ISL_2828114, EPI_ISL_2828131, EPI_ISL_2885277, EPI_ISL_2885342, EPI_ISL_2938032, EPI_ISL_2938048, EPI_ISL_2938086, EPI_ISL_2982418, EPI_ISL_3009909, EPI_ISL_3009910, EPI_ISL_3009942, EPI_ISL_3060338, EPI_ISL_3060366, EPI_ISL_3154413, EPI_ISL_3154414, EPI_ISL_3189784, EPI_ISL_3246748, EPI_ISL_3246761, EPI_ISL_3326263 | | | |
| see above | Hospital Universitari Vall d'Hebron - Vall d'Hebron Institut de Recerca | Hospital Universitari Vall d'Hebron - Vall d'Hebron Institut de Recerca | Alejandra González-Sánchez; Andrés Antón; Ariadna Rando; Carla Castillo; Cristina Andrés; Damir Garcia-Cehic; Josep Quer; Juliana Esperalba; Karen García; Maria Carmen Martin; Maria Gema Codina; Maria Piñana; Rodrigo Vásquez; Tomàs Pumarola |
| EPI_ISL_3118358 | Hospital Universitat de Bellvitge | Microbiology Department | Aida Gonzalez-Diaz; Carmen Ardanuy; Jordi Camara; Jordi Niubó; Laura Calatayud; M Angeles Domínguez; Sara Marti; Yolanda Hernandez |
| EPI_ISL_3259729 | Hospital de Base de Sao Jose do Rio Preto | Instituto Adolfo Lutz, Interdiciplinary Procedures Center, Strategic Laboratory | Caio Vinicius Dias Lopes; Claudia Regina Gonçalves; Claudio Tavares Sacchi; Erica Valessa Ramos Gomes; Karoline Rodrigues Campos; Leonardo Tadeu de Araujo; Marlon Benedito Nascomento Santos |
| EPI_ISL_3696703, EPI_ISL_3696736 | Hospital de Campanha de Guaratinguetá | Instituto Butantan | Antonio Jorge Martins; Claudia Renata dos Santos Barros; David Schlesinger; Debora Botequio Moretti; Dimas Tadeu Covas; Elaine Cristina Marqueze; Elaine Vieira Santos; Evandra Strazza Rodrigues; Heidge Fukumasu; Jayme Augusto de Souza-Neto; José Salvatore Leister Patané; Luiz Alcantara; Luiz Lehmann Coutinho; Maria Carolina Elias; Maurício Lacerda Nogueira; Rafael dos Santos Bezerra; Raul Machado Neto; Rejane Maria Tommasini Grotto; Ricardo Haddad; Sandra Coccuzzo Sampaio Vessoni; Simone Kashima; Svetoslav Nanev Slavov; Vincent Louis Viala |
| EPI_ISL_3115715, EPI_ISL_3936902 | IN State Department of Health Laboratory Services | IN State Department of Health Laboratory Services | Brian Pope; Cassandra Campion; Jamie Yeadon; Kyle Brownlee; Lixia Liu; Mark Glazier; Melissa Hindenlang |
| EPI_ISL_2774217, EPI_ISL_2774219, EPI_ISL_2774220 | IRCCS San Gallicano Dermatological Institute | IRCCS Regina Elena National Cancer Institute | Aldo Morrone; Eleonora Sperandio; Fabrizio Ensoli; Francesca De Nicola; Frauke Goeman; Fulvia Pimpinelli; Gennaro Ciliberto; Giovanni Blandino; Giulia Orlandi; Grazia Prignano; Matteo Pallocca; Maurizio Fanciulli; Sabrina Strano; Sara Donzelli; Sara Petrolo; Serena Salvo |
| EPI_ISL_3259693 | Instituto Adolfo Lutz - Regional de Campinas | Instituto Adolfo Lutz, Interdiciplinary Procedures Center, Strategic Laboratory | Caio Vinicius Dias Lopes; Claudia Regina Gonçalves; Claudio Tavares Sacchi; Erica Valessa Ramos Gomes; Karoline Rodrigues Campos; Leonardo Tadeu de Araujo; Marlon Benedito Nascomento Santos |
| EPI_ISL_3545786, EPI_ISL_3717852, EPI_ISL_3717856, EPI_ISL_3717862 | Instituto Adolfo Lutz - Regional de Marilia | Instituto Adolfo Lutz, Interdiciplinary Procedures Center, Strategic Laboratory | Caio Vinicius Dias Lopes; Claudia Regina Gonçalves; Claudio Tavares Sacchi; Karoline Rodrigues Campos; Leonardo Tadeu de Araujo; Marlon Benedito Nascimento Santos; Marlon Benedito Nascomento Santos |
| EPI_ISL_3316178, EPI_ISL_3316206, EPI_ISL_3316302, EPI_ISL_3316304, EPI_ISL_3316306 | Instituto Adolfo Lutz - Regional de Ribeirao Preto | Instituto Adolfo Lutz, Interdiciplinary Procedures Center, Strategic Laboratory | Caio Vinicius Dias Lopes; Claudia Regina Gonçalves; Claudio Tavares Sacchi; Karoline Rodrigues Campos; Leonardo Tadeu de Araujo; Marlon Benedito Nascomento Santos |
| EPI_ISL_3946610 | Instituto Adolfo Lutz - Regional de Santo Andre | Instituto Adolfo Lutz, Interdiciplinary Procedures Center, Strategic Laboratory | Claudio Tavares Sacchi; Karoline Rodrigues Campos |
| EPI_ISL_3671889 | Instituto Adolfo Lutz - Regional de Santos | Instituto Adolfo Lutz, Interdisciplinary Procedures Center, Strategic Laboratory | Caio Vinicius Dias Lopes; Claudia Regina Gonçalves; Claudio Tavares Sacchi; Karoline Rodrigues Campos; Leonardo Tadeu de Araujo; Marlon Benedito Nascomento Santos |
| EPI_ISL_3259719, EPI_ISL_3259727, EPI_ISL_3545769, EPI_ISL_3545777, EPI_ISL_3545778, EPI_ISL_3545779 | Instituto Adolfo Lutz - Regional de Sao Jose do Rio Preto | Instituto Adolfo Lutz, Interdiciplinary Procedures Center, Strategic Laboratory | Caio Vinicius Dias Lopes; Claudia Regina Gonçalves; Claudio Tavares Sacchi; Erica Valessa Ramos Gomes; Karoline Rodrigues Campos; Leonardo Tadeu de Araujo; Marlon Benedito Nascomento Santos |
| EPI_ISL_3259679, EPI_ISL_3259681 | Instituto Adolfo Lutz - Regional de Sorocaba | Instituto Adolfo Lutz, Interdiciplinary Procedures Center, Strategic Laboratory | Caio Vinicius Dias Lopes; Claudia Regina Gonçalves; Claudio Tavares Sacchi; Erica Valessa Ramos Gomes; Karoline Rodrigues Campos; Leonardo Tadeu de Araujo; Marlon Benedito Nascomento Santos |
| EPI_ISL_2919283, EPI_ISL_3316270, EPI_ISL_3864576 | Instituto Adolfo Lutz Central | Instituto Adolfo Lutz, Interdiciplinary Procedures Center, Strategic Laboratory | Caio Vinicius Dias Lopes; Claudia Regina Gonçalves; Claudio Tavares Sacchi; Erica Valessa Ramos Gomes; Karoline Rodrigues Campos; Leonardo Tadeu de Araujo; Marlon Benedito Nascomento Santos |
| EPI_ISL_3761514 | Instituto Adolfo Lutz Central | Instituto Adolfo Lutz, Rapid Response Center, Strategic Laboratory | Claudio Tavares Sacchi; Karoline Rodrigues Campos |
| EPI_ISL_3983517 | Instituto Nacional de Cardiología (INC) | Centro de Investigación en Enfermedades Infecciosas (CIENI), Instituto Nacional de Enfermedades Respiratorias (INER) | Alejandra García-Gasca; Alejandra Hernández-Terán; Alejandro Sánchez-Flores; Alfredo Herrera-Estrella; Alicia Ocaña-Mondragón; Andreu Comas-García; Angel Gustavo Salas-Lais; Antonio Loza Román; Bernardo Martínez-Miguel; Blanca Taboada; Brenda Irasema Maldonado-Meza; Bruno Gómez-Gil; Carla Rodríguez; Daniel Fregoso-Rueda; Daniel Lira Morales; Eduardo Becerril-Vargas; Eduardo Rivera-Martínez; Fernando Fontove-Herrera; Fidencio Mejía-Nepomuceno; Francisco Pulido; Gabriel Chavira-Trujillo; Gloria Elena Espinosa-Ayala; Gloria María Molina-Salinas; Gloria Vazquez; Hector Montoya-Fuentes; Helen Haydee Fernanda Ramírez-Plascencia; Irvin González-López; Jean Pierre González; Jesús Hernández; Joel Armando Vázquez-Pérez).; Jorge Salas-Hernández; José Antonio Enciso-Moreno; José Arturo Martínez-Orozco; José Esteban Muñoz-Medina; José de Jesús Nuñez-Contreras; Juan Bautista Chale-Dzul; Julissa Enciso-Ibarra; Kathia Elizabeth Tapia-Diaz; Luis Alberto Ochoa-Carrera; Margarita Matías-Florentino; Mario Mújica-Sánchez; Marissa Perez-Garcia; María Eugenia Jiménez-Corona; María Guadalupe Santiago-Mauricio; María Guadalupe de Jesús Mireles-Rivera; Nelly Sélem-Mojica; Pavel Isa; Ricardo Ciria Merce; Ricardo Grande; Rosa María Gutiérrez Rios; Rosario Vazquez-Larios; Santiago Ávila-Ríos; Selene Zárate; Susana Lopez; Verónica Mata-Haro; Victor Eduardo García-Arias; Víctor Hugo Borja-Aburto |
| EPI_ISL_3277577 | Instituto Nacional de Medicina Genomica | Instituto Nacional de Medicina Genomica | Cedro-Tanda A; Escobar-Arrazola MA; Herrera-Montalvo LA.; Hidalgo-Miranda A; Mendoza-Vargas A; Munguia-Garza P; Ramirez-Vega O; Rangel-DeLeon D; Reyes-Grajeda JP |
| EPI_ISL_2844013, EPI_ISL_2844023, EPI_ISL_2844031, EPI_ISL_2844033, EPI_ISL_2844034, EPI_ISL_2844039, EPI_ISL_2844046, EPI_ISL_2844074, EPI_ISL_2844088, EPI_ISL_2844095, EPI_ISL_2844101, EPI_ISL_2844123, EPI_ISL_2844124, EPI_ISL_2844130, EPI_ISL_2844146, EPI_ISL_2844153, EPI_ISL_2844164, EPI_ISL_2844166, EPI_ISL_2844167, EPI_ISL_2844169, EPI_ISL_2844173, EPI_ISL_2844174, EPI_ISL_2844180, EPI_ISL_2844186, EPI_ISL_3118926, EPI_ISL_3118929, EPI_ISL_3118931, EPI_ISL_3118934, EPI_ISL_3244548, EPI_ISL_3244556, EPI_ISL_3244581, EPI_ISL_3824274, EPI_ISL_3824285, EPI_ISL_3824289, EPI_ISL_3824306, EPI_ISL_3824314, EPI_ISL_3824315, EPI_ISL_3824316, EPI_ISL_3824324, EPI_ISL_3824325, EPI_ISL_3824327, EPI_ISL_3824328, EPI_ISL_3824331, EPI_ISL_3824334, EPI_ISL_3824335, EPI_ISL_3824342, EPI_ISL_3824343, EPI_ISL_3824344, EPI_ISL_3824345 | | | |
| see above | Instituto de Biotecnologia - UNESP-Botucatu-SP | Instituto de Biotecnologia - UNESP-Botucatu-SP | Cecília Artico Banho; Cíntia Bittar; Fábio Sossai Possebon; Guilherme Campos; Helena Lage Ferreira; Jorge A. Petroli Marchesi; João Pessoa Araújo Jr.; Leila Sabrina Ullmann; Lívia Sacchetto; Maisa C. Pereira Parra; Marília Moraes; Maurício L. Nogueira; Paula Rahal; Paulo Inacio da Costa |
| EPI_ISL_2550983, EPI_ISL_2550985, EPI_ISL_2551114, EPI_ISL_2551117, EPI_ISL_2844367, EPI_ISL_2844377, EPI_ISL_2844381, EPI_ISL_2844389, EPI_ISL_2844401, EPI_ISL_2844428, EPI_ISL_2844431, EPI_ISL_2844471, EPI_ISL_2844476, EPI_ISL_2844477, EPI_ISL_2844479, EPI_ISL_2844480, EPI_ISL_2844484 | | | |
| see above | Istituto Zooprofilattico Sperimentale del Mezzogiorno | TIGEM | Antonio Grimaldi Patrizia Annunziata Francesco Panariello Biancamaria Pierri Claudia Tiberio Teresa Giuliano Valentina Bouche Chiara Colantuono Maria Concetta Cuomo Denise Di Concilio Lucio Di Filippo Anna Manfredi Marcello Salvi Antonio Limone Luigi Atripaldi Pellegrino Cerino Andrea Ballabio Davide Cacchiarelli |
| EPI_ISL_2641746, EPI_ISL_2641747, EPI_ISL_2641750, EPI_ISL_2641751, EPI_ISL_2641753, EPI_ISL_2641754, EPI_ISL_2641775, EPI_ISL_2641782, EPI_ISL_2641789, EPI_ISL_2641793, EPI_ISL_2641794, EPI_ISL_2641800, EPI_ISL_2641816, EPI_ISL_2641817, EPI_ISL_2641818, EPI_ISL_2641820, EPI_ISL_2641824, EPI_ISL_2641829, EPI_ISL_2641862, EPI_ISL_2641865, EPI_ISL_2641884, EPI_ISL_2641907, EPI_ISL_2641938, EPI_ISL_2641973, EPI_ISL_2642007, EPI_ISL_2642011, EPI_ISL_2642018, EPI_ISL_2642020, EPI_ISL_2642029, EPI_ISL_2642040, EPI_ISL_2725363, EPI_ISL_2725393, EPI_ISL_2725394, EPI_ISL_2725407 | | | |
| see above | Istituto Zooprofilattico Sperimentale del Mezzogiorno | Telethon Institute of Genetics and Medicine (TIGEM) | Antonio Grimaldi Patrizia Annunziata Francesco Panariello Biancamaria Pierri Claudia Tiberio Teresa Giuliano Valentina Bouche Chiara Colantuono Maria Concetta Cuomo Denise Di Concilio Lucio Di Filippo Anna Manfredi Marcello Salvi Antonio Limone Luigi Atripaldi Pellegrino Cerino Andrea Ballabio Davide Cacchiarelli |
| EPI_ISL_2611997, EPI_ISL_3485854 | Istituto Zooprofilattico Sperimentale della Puglia e della Basilicata | Istituto Zooprofilattico Sperimentale della Puglia e della Basilicata | Bianco A.; Capozzi L.; Del Sambro L.; Difato L.; Galante D.; Pace L.; Parisi A.; Pennuzzi G.; Simone D. |
| EPI_ISL_3384937 | KAISER SANTA CLARA | Santa Clara County Public Health Laboratory | Santa Clara County Public Health Department |
| EPI_ISL_3404554, EPI_ISL_3404555, EPI_ISL_3404556, EPI_ISL_3404558, EPI_ISL_3404559, EPI_ISL_3404560, EPI_ISL_3404561, EPI_ISL_3404570, EPI_ISL_3404571, EPI_ISL_3404572, EPI_ISL_3404573, EPI_ISL_3404579, EPI_ISL_3404585, EPI_ISL_3404586, EPI_ISL_3404588, EPI_ISL_3404590, EPI_ISL_3404591, EPI_ISL_3404607, EPI_ISL_3404608, EPI_ISL_3404864, EPI_ISL_3404865, EPI_ISL_3404866, EPI_ISL_3841241, EPI_ISL_3841252, EPI_ISL_3841253, EPI_ISL_3841254, EPI_ISL_3841256, EPI_ISL_3841257, EPI_ISL_3841258, EPI_ISL_3841259, EPI_ISL_3841285, EPI_ISL_3841290, EPI_ISL_3841291, EPI_ISL_3841292, EPI_ISL_3841315, EPI_ISL_3841316, EPI_ISL_3841317 | | | |
| see above | LACEN | LGBio (Laboratorio de Genetica & Biodiversidade) | Alex Honda Bernardes; Amanda Alves de Melo; Aparecido Divino da Cruz; Cintia Pelegrineti Targueta de Azevedo Brito; Daniela de Melo e Silva; Elisangela de Paula Silveira Lacerda; Juliana Santana de Curcio; Luiz Augusto Pereira; Marc Alexandre Duarte Gigonzac; Mariana Pires de Campos Telles; Ramilla dos Santos Braga; Renata de Oliveira Dias; Rhewter Nunes; Thais Cidalia Vieira Gigonzac; Thais Cidália Vieira Gigonzac; Thais Guimarães Castro; Thays Millena Alves Pedroso |
| EPI_ISL_3046269, EPI_ISL_3046279, EPI_ISL_3046324, EPI_ISL_3134664, EPI_ISL_3134673, EPI_ISL_3134674, EPI_ISL_3134675, EPI_ISL_3134678, EPI_ISL_3134679, EPI_ISL_3134711, EPI_ISL_3447418, EPI_ISL_3447453, EPI_ISL_3447456, EPI_ISL_3447464, EPI_ISL_3447489, EPI_ISL_3447572, EPI_ISL_3447581, EPI_ISL_3447586, EPI_ISL_3447588, EPI_ISL_3447599, EPI_ISL_3703737, EPI_ISL_3835243, EPI_ISL_3835255, EPI_ISL_3835261, EPI_ISL_3835269, EPI_ISL_3835271, EPI_ISL_3835288, EPI_ISL_3835289, EPI_ISL_3835291, EPI_ISL_3835292, EPI_ISL_3835293, EPI_ISL_3835294 | | | |
| see above | LACEN/PE | WallauLab on behalf of Fiocruz COVID-19 Genomic Surveillance Network | Alexandre Freitas da Silva; Cassia Docena; Constância Flávia Junqueira Ayres; Filipe Zimmer Dezordi; Gabriel Luz Wallau; Gustavo Barbosa de Lima; Lais Ceschini Machado; Lilian Caroliny Amorim Silva; Marcelo Henrique dos Santos Paiva; Matheus Filgueira Bezerra; Sinval Pinto Brandão Filho |
| EPI_ISL_3912231 | LACEN_CENTRO DE TESTAGEM PARA VIAJANTE | ACME Lab, Oswaldo Cruz Foundation, FIOCRUZ/CE | Cleber Furtado Aksenen; Fabio Miyajima; Fernando Braga Stehling; Francisco Eder de Moura Lopes; Jamille Maria Mendes Bezerra; Joaquim Cesar do Nascimento Sousa Junior; Pedro Miguel Carneiro Jeronimo; Suzana Porto Almeida & Lucas Delerino on behalf of COVID-19 FIOCRUZ Genomic Network; Thais Ferreira de Oliveira; Thais de Oliveira Costa; Ticiane Cavalcante de Souza; Veridiana Pessoa Miyajima |
| EPI_ISL_3010763, EPI_ISL_3010767, EPI_ISL_3010913, EPI_ISL_3010928, EPI_ISL_3020990, EPI_ISL_3047724, EPI_ISL_3134937, EPI_ISL_3134958, EPI_ISL_3134963, EPI_ISL_3246164, EPI_ISL_3246197, EPI_ISL_3261089, EPI_ISL_3261108, EPI_ISL_3267913, EPI_ISL_3267927, EPI_ISL_3267932, EPI_ISL_3267939, EPI_ISL_3267964, EPI_ISL_3268000, EPI_ISL_3268008, EPI_ISL_3268015, EPI_ISL_3385148, EPI_ISL_3385154, EPI_ISL_3505188, EPI_ISL_3505189, EPI_ISL_3505190, EPI_ISL_3505192, EPI_ISL_3505194, EPI_ISL_3505198, EPI_ISL_3505199, EPI_ISL_3505200, EPI_ISL_3505202, EPI_ISL_3505208, EPI_ISL_3505209, EPI_ISL_3804726, EPI_ISL_3804732, EPI_ISL_3804734, EPI_ISL_3804737, EPI_ISL_3804739, EPI_ISL_3804740, EPI_ISL_3804742, EPI_ISL_3804743, EPI_ISL_3941507, EPI_ISL_3941514, EPI_ISL_3941516, EPI_ISL_3941519 | | | |
| see above | LATE - Laboratório de Técnicas Especiais - Hospital Israelita Albert Einstein | LATE - Laboratório de Técnicas Especiais - Hospital Israelita Albert Einstein | Alexandre Hideaki Takara; Ana Paula Moreira Salles; Anelisie da Silva Santos; Deyvid Amgarten; Erick Gustavo Dorlass; Fernanda de Mello Malta; João Renato Rebello Pinho; Marcio Anunciacao Menezes; Pedro Henrique Sebe Rodrigues; Raquel Riyuzo |
| EPI_ISL_3923749 | LESP Baja California | Instituto de Diagnostico y Referencia Epidemiologicos | Abril Rodriguez-Maldonado; Ariadna Medina-Benitez; Claudia Wong-Arambula; Ernesto Ramirez-Gonzalez.; Gisela Barrera-Badillo; Irma Lopez-Martinez; Joaquin Quiroz-Mercado; Lucia Hernandez-Rivas; Maribel Gonzalez-Villa; Natividad Cruz-Ortiz; Sergio Rangel-Guerrero; Tatiana Nunez-Garcia; Vanessa Rivero-Arredondo |

| | | | |
|---|---|---|---|
| | | (INDRE) | |
| EPI_ISL_3459959 | LESP Tamaulipas | Instituto de Diagnostico y Referencia Epidemiologicos (INDRE) | Abril Rodriguez-Maldonado; Ariadna Medina-Benitez; Claudia Wong-Arambula; Ernesto Ramirez-Gonzalez.; Gisela Barrera-Badillo; Irma Lopez-Martinez; Joaquin Quiroz-Mercado; Lucia Hernandez-Rivas; Maribel Gonzalez-Villa; Natividad Cruz-Ortiz; Sergio Rangel-Guerrero; Tatiana Nunez-Garcia; Vanessa Rivero-Arredondo |
| EPI_ISL_3265604 | LESP Veracruz | Instituto de Diagnostico y Referencia Epidemiologicos (INDRE) | Abril Rodriguez-Maldonado; Ariadna Medina-Benitez; Claudia Wong-Arambula; Ernesto Ramirez-Gonzalez.; Gisela Barrera-Badillo; Irma Lopez-Martinez; Joaquin Quiroz-Mercado; Lucia Hernandez-Rivas; Maribel Gonzalez-Villa; Natividad Cruz-Ortiz; Sergio Rangel-Guerrero; Tatiana Nunez-Garcia; Vanessa Rivero-Arredondo |
| EPI_ISL_3214387, EPI_ISL_3214389, EPI_ISL_3214390, EPI_ISL_3214391, EPI_ISL_3214392, EPI_ISL_3214397, EPI_ISL_3214398, EPI_ISL_3214405, EPI_ISL_3214406, EPI_ISL_3214426, EPI_ISL_3214430, EPI_ISL_3214488, EPI_ISL_3214524, EPI_ISL_3214611, EPI_ISL_3214612, EPI_ISL_3214616, EPI_ISL_3214630, EPI_ISL_3276430, EPI_ISL_3276431, EPI_ISL_3276432 | | | |
| see above | Lab. Microbiologia e Virologia Cotugno A.O. dei Colli | TIGEM | Antonio Grimaldi Patrizia Annunziata Francesco Panariello Biancamaria Pierri Claudia Tiberio Teresa Giuliano Valentina Bouche Chiara Colantuono Maria Concetta Cuomo Denise Di Concilio Lucio Di Filippo Anna Manfredi Marcello Salvi Antonio Limone Luigi Atripaldi Pellegrino Cerino Andrea Ballabio Davide Cacchiarelli |
| EPI_ISL_2844492, EPI_ISL_2844510, EPI_ISL_2844517 | Lab. Microbiologia e Virologia Cotugno A.O. dei Colli - Istituto Zooprofilattico Sperimentale del Mezzogiorno | TIGEM | Antonio Grimaldi Patrizia Annunziata Francesco Panariello Biancamaria Pierri Claudia Tiberio Teresa Giuliano Valentina Bouche Chiara Colantuono Maria Concetta Cuomo Denise Di Concilio Lucio Di Filippo Anna Manfredi Marcello Salvi Antonio Limone Luigi Atripaldi Pellegrino Cerino Andrea Ballabio Davide Cacchiarelli |
| EPI_ISL_2725152, EPI_ISL_2725175, EPI_ISL_2725183, EPI_ISL_2725184, EPI_ISL_2725186, EPI_ISL_2725190, EPI_ISL_2725193, EPI_ISL_2725195, EPI_ISL_2725198, EPI_ISL_2725206, EPI_ISL_2725209, EPI_ISL_2725210, EPI_ISL_2725212, EPI_ISL_2725213, EPI_ISL_2725216, EPI_ISL_2725229, EPI_ISL_2725238, EPI_ISL_2725239, EPI_ISL_2725245, EPI_ISL_2725248, EPI_ISL_2725249, EPI_ISL_2725250, EPI_ISL_2725252, EPI_ISL_2725256, EPI_ISL_2725257, EPI_ISL_2725263, EPI_ISL_2725265, EPI_ISL_2725266, EPI_ISL_2725268, EPI_ISL_2725271, EPI_ISL_2725275, EPI_ISL_2725276, EPI_ISL_2725281, EPI_ISL_2725282, EPI_ISL_2725294 | | | |
| see above | Lab. Microbiologia e Virologia Cotugno A.O. dei Colli - Istituto Zooprofilattico Sperimentale del Mezzogiorno | Telethon Institute of Genetics and Medicine (TIGEM) | Antonio Grimaldi Patrizia Annunziata Francesco Panariello Biancamaria Pierri Claudia Tiberio Teresa Giuliano Valentina Bouche Chiara Colantuono Maria Concetta Cuomo Denise Di Concilio Lucio Di Filippo Anna Manfredi Marcello Salvi Antonio Limone Luigi Atripaldi Pellegrino Cerino Andrea Ballabio Davide Cacchiarelli |
| EPI_ISL_3497506, EPI_ISL_3498165 | Labor Mönchengladbach MVZ Dr. Stein + Kollegen GbR | Robert Koch Institute | |
| EPI_ISL_3334766 | Laboratoires d'analyses medicales - Ketterthill | Laboratoire national de sante, Microbiology, Microbial Genomics Platform | Anke Wienecke-Baldacchino; Caroline Scheiber; Catherine Ragimbeau; Elodie Solarino; Fatu Djabi; Jessica Tapp; Lise Pignon; Raoul Salmon; Serge Vedy; Tamir Abdelrahman; Virginie Jover |
| EPI_ISL_2983184, EPI_ISL_2983230, EPI_ISL_3190353, EPI_ISL_3190356, EPI_ISL_3539229, EPI_ISL_3539230, EPI_ISL_3539232, EPI_ISL_3539233, EPI_ISL_3539234, EPI_ISL_3802964 | | | |
| see above | Laboratorio Central de Saude Publica do Estado de Goias (LACEN/GO) | Laboratory of Respiratory Viruses and Measles, Oswaldo Cruz Institute, FIOCRUZ | Agatha Cristinne Prudencio; Alice Sampaio Rocha; Ana Carolina Mendonca; Ana Flavia Mendonça; Anna Carolina Paixao; Carmen Helena Ramos; Cassiane Casanova; Elisa Cavalcante Pereira; Fernando Motta; Flavia Pereira Amorim da Silva; Ighor Leonardo Arantes Gomes; Luciana Appolinario; Luiz Augusto Pereira; Marilda Siqueira on behalf of the Fiocruz COVID-19 Genomic Surveillance Network; Paola Resende; Rafael Souza Guedes; Renata Serrano Lopes; Taina Venas; Vinicius Lemes da Silva |
| EPI_ISL_2863818, EPI_ISL_2863836, EPI_ISL_3539238, EPI_ISL_3802965, EPI_ISL_3802967, EPI_ISL_3802972, EPI_ISL_3802974 | | | |
| see above | Laboratorio Central de Saude Publica do Estado do Para (LACEN/PA) | Laboratory of Respiratory Viruses and Measles, Oswaldo Cruz Institute, FIOCRUZ | Agatha Cristinne Prudencio Soares; Agatha Soares; Alice Sampaio Rocha; Ana Carolina Mendonca; Anna Carolina Paixao; Elisa Cavalcante Pereira; Fernando Motta; Ighor Arantes; Ighor Leonardo Arantes Gomes; Luciana Appolinario; Marilda Siqueira on behalf of the Fiocruz COVID-19 Genomic Surveillance Network; Paola Resende; Renata Serrano Lopes; Taina Venas; Valnete Andrade |
| EPI_ISL_3245110, EPI_ISL_3245265, EPI_ISL_3245266, EPI_ISL_3245317, EPI_ISL_3615218, EPI_ISL_3615225, EPI_ISL_3615229, EPI_ISL_3615241, EPI_ISL_3920958, EPI_ISL_3921076, EPI_ISL_3921319 | | | |
| see above | Laboratorio Central Noel Nutels | Bioinformatics Laboratory / LNCC | Alessandra P Lamarca; Alexandra L Gerber; Amilcar Tanuri; Ana Paula de C Guimaraes; Ana Tereza R Vasconcelos; Andrea Cony Cavalcanti; Caio Luiz Pereira Ribeiro; Cintia Policarpo; Claudia Maria Braga de Mello; Cristiane Gomes da Silva; Douglas Terra Machado; Erica Ramos dos Santos Nascimento; Fernanda Leitao dos Santos; Flavio Dias da Silva; Gleidson da Silva de Oliveira; Leandro Magalhaes de Souza; Liliane Cavalcante; Luiz G P de Almeida; Marcio Henrique de Oliveira Garcia; Mario Sergio Ribeiro; Ricardo Jose Barbosa Salviano; Ronaldo da Silva F Jr; Silvia Carvalho |
| EPI_ISL_2942818, EPI_ISL_2942829, EPI_ISL_2942909, EPI_ISL_2942934 | Laboratorio Central de Epidemiologia (LCE) | Unidad de Genomica Avanzada | ; Alejandra Garcia-Gasca; Alejandra Hernandez-Teran; Alejandro Sanchez-Flores; Alfredo Herrera-Estrella; Alicia Ocaña-Mondragon; Andreu Comas-Garcia; Angel Gustavo Salas-Lais; Antonio Loza Roman; Bernardo Martinez-Miguel; Blanca Taboada; Brenda Irasema Maldonado-Meza; Bruno Gomez-Gil; Carla Ivon Herrera-Najera; Carlos F. Arias; Celia Boukadida; Celida Duque Molina; Celida Martinez- Rodriguez; Clara Esperanza Santacruz-Tinoco; Concepcion Grajales-Muñiz; Consorcio Mexicano de Vigilancia Genomica (CoViGen-Mex). Authors (in alphabetical order): Julio Elias Alvarado-Yaah; Cristobal Chaidez-Quiroz; Daniel Fregoso-Rueda; Daniel Lira Morales; Eduardo Becerril-Vargas; Fernando Fontove-Herrera; Fidencio Mejia-Nepomuceno; Francisco Pulido; Gloria Elena Espinosa-Ayala; Gloria Maria Molina-Salinas; Gloria Vazquez; Hector Esteban Paz-Juarez; Hector Montoya-Fuentes; Helen Haydee Fernanda Ramirez-Plascencia; Irvin Gonzalez-Lopez; Jean Pierre Gonzalez; Jesus Hernandez; Joel Armando Vazquez-Perez.; Jorge Salas-Hernandez; Jose Antonio Enciso-Moreno; Jose Arturo Martinez-Orozco; Jose Esteban Muñoz-Medina; Jose de Jesus Nuñez-Contreras; Juan Bautista Chale-Dzul; Julissa Enciso-Ibarra; Luis Alberto Ochoa-Carrera; Margarita Matias-Florentino; Maria Guadalupe Santiago-Mauricio; Maria Guadalupe de Jesus Mireles-Rivera; Mario Mujica-Sanchez; Marissa Perez-Garcia; Nelly Selem-Mojica; Pavel Isa; Ricardo Ciria Merce; Ricardo Grande; Rosa Maria Gutierrez Rios; Santiago avila-Rios; Selene Zarate; Susana Lopez; Veronica Mata-Haro; Victor Eduardo Garcia-Arias; Victor Hugo Borja-Aburto |
| EPI_ISL_3805732 | Laboratorio Central de Epidemiologia (LCE) | Unidad de Genomica Avanzada | ; Alejandra Garcia-Gasca; Alejandra Hernandez-Teran; Alejandro Sanchez-Flores; Alfredo Herrera-Estrella; Alicia Ocaña-Mondragon; Andreu Comas-Garcia; Angel Gustavo Salas-Lais; Antonio Loza Roman; Bernardo Martinez-Miguel; Blanca Taboada; Brenda Irasema Maldonado-Meza; Bruno Gomez-Gil; Carla Ivon Herrera-Najera; Carlos F. Arias; Celia Boukadida; Celida Duque Molina; Celida Martinez- Rodriguez; Clara Esperanza Santacruz-Tinoco; Concepcion Grajales-Muñiz; Consorcio Mexicano de Vigilancia Genomica (CoViGen-Mex). Authors (in alphabetical order): Julio Elias Alvarado-Yaah; Cristobal Chaidez-Quiroz; Daniel Fregoso-Rueda; Daniel Lira Morales; Eduardo Becerril-Vargas; Fernando Fontove-Herrera; Fidencio Mejia-Nepomuceno; Francisco Pulido; Gloria Elena Espinosa-Ayala; Gloria Maria Molina-Salinas; Gloria Vazquez; Hector Esteban Paz-Juarez; Hector Montoya-Fuentes; Helen Haydee Fernanda Ramirez-Plascencia; Irvin Gonzalez-Lopez; Jean Pierre Gonzalez; Jesus Hernandez; Joel Armando Vazquez-Perez.; Jorge Salas-Hernandez; Jose Antonio Enciso-Moreno; Jose Arturo Martinez-Orozco; Jose Esteban Muñoz-Medina; Jose de Jesus Nuñez-Contreras; Juan Bautista Chale-Dzul; Julissa Enciso-Ibarra; Luis Alberto Ochoa-Carrera; Margarita Matias-Florentino; Maria Guadalupe Santiago-Mauricio; Maria Guadalupe de Jesus Mireles-Rivera; Mario Mujica-Sanchez; Marissa Perez-Garcia; Nelly Selem-Mojica; Pavel Isa; Ricardo Ciria Merce; Ricardo Grande; Rosa Maria Gutierrez Rios; Santiago avila-Rios; Selene Zarate; Susana Lopez; Veronica Mata-Haro; Victor Eduardo Garcia-Arias; Victor Hugo Borja-Aburto |
| EPI_ISL_3155550, EPI_ISL_3155602 | Laboratorio Central de Epidemiología (LCE) | Centro de Investigación en Enfermedades Infecciosas (CIENI), Instituto Nacional de Enfermedades Respiratorias (INER) | ; Alejandra García-Gasca; Alejandra Hernández-Terán; Alejandro Sánchez-Flores; Alfredo Herrera-Estrella; Alicia Ocaña-Mondragón; Andreu Comas-García; Angel Gustavo Salas-Lais; Antonio Loza Román; Bernardo Martínez-Miguel; Blanca Taboada; Brenda Irasema Maldonado-Meza; Bruno Gómez-Gil; Carla Ivón Herrera-Najera; Carlos F. Arias; Celia Boukadida; Clara Esperanza Santacruz-Tinoco; Concepción Grajales-Muñiz; Consorcio Mexicano de Vigilancia Genómica (CoViGen-Mex). Authors (in alphabetical order): Julio Elias Alvarado-Yaah; Cristóbal Cháidez-Quiróz; Célida Duque Molina; Célida Martínez- Rodríguez; Daniel Fregoso-Rueda; Daniel Lira Morales; Eduardo Becerril-Vargas; Fernando Fontove-Herrera; Fidencio Mejía-Nepomuceno; Francisco Pulido; Gloria Elena Espinosa-Ayala; Gloria María Molina-Salinas; Gloria Vazquez; Hector Esteban Paz-Juárez; Hector Montoya-Fuentes; Helen Haydee Fernanda Ramírez-Plascencia; Irvin González-López; Jean Pierre González; Jesús Hernández; Joel Armando Vázquez-Pérez.; Jorge Salas-Hernández; José Antonio Enciso-Moreno; José Arturo Martínez-Orozco; José Esteban Muñoz-Medina; José de Jesús Nuñez-Contreras; Juan Bautista Chale-Dzul; Julissa Enciso-Ibarra; Kathia Elizabeth Tapia-Diaz; Luis Alberto Ochoa-Carrera; Margarita Matías-Florentino; Mario Mújica-Sánchez; Marissa Perez-Garcia; María Guadalupe Santiago-Mauricio; María Guadalupe de Jesús Mireles-Rivera; Nelly Sélem-Mojica; Pavel Isa; Ricardo Ciria Merce; Ricardo Grande; Rosa María Gutiérrez Rios; Santiago Ávila-Ríos; Selene Zárate; Susana Lopez; Verónica Mata-Haro; Victor Eduardo García-Arias; Víctor Hugo Borja-Aburto |
| EPI_ISL_2801850, EPI_ISL_2801876, EPI_ISL_3347552 | Laboratorio Central de Epidemiología (LCE) | Instituto de Biotecnología de la UNAM | ; Alejandra García-Gasca; Alejandra Hernández-Terán; Alejandro Sánchez-Flores; Alfredo Herrera-Estrella; Alicia Ocaña-Mondragón; Andreu Comas-García; Angel Gustavo Salas-Lais; Antonio Loza Román; Bernardo Martínez-Miguel; Blanca Taboada; Brenda Irasema Maldonado-Meza; Bruno Gómez-Gil; Carla Ivón Herrera-Najera; Carlos F. Arias; Celia Boukadida; Clara Esperanza Santacruz-Tinoco; Concepción Grajales-Muñiz; Consorcio Mexicano de Vigilancia Genómica (CoViGen-Mex). Authors (in alphabetical order): Julio Elias Alvarado-Yaah; Cristóbal Cháidez-Quiróz; Célida Duque Molina; Célida Martínez- Rodríguez; Daniel Fregoso-Rueda; Daniel Lira Morales; Eduardo Becerril-Vargas; Fernando Fontove-Herrera; Fidencio Mejía-Nepomuceno; Francisco Pulido; Gloria Elena Espinosa-Ayala; Gloria María Molina-Salinas; Gloria Vazquez; Hector Esteban Paz-Juárez; Hector Montoya-Fuentes; Helen Haydee Fernanda Ramírez-Plascencia; Irvin González-López; Jean Pierre González; Jesús Hernández; Joel Armando Vázquez-Pérez).; Joel Armando Vázquez-Pérez.; Jorge Salas-Hernández; José Antonio Enciso-Moreno; José Arturo Martínez-Orozco; José Esteban Muñoz-Medina; José de Jesús Nuñez-Contreras; Juan Bautista Chale-Dzul; Julissa Enciso-Ibarra; Kathia Elizabeth Tapia-Diaz; Luis Alberto Ochoa-Carrera; Margarita Matías-Florentino; Mario Mújica-Sánchez; Marissa Perez-Garcia; María Guadalupe Santiago-Mauricio; María Guadalupe de Jesús Mireles-Rivera; Nelly Sélem-Mojica; Pavel Isa; Ricardo Ciria Merce; Ricardo Grande; Rosa María Gutiérrez Rios; Santiago Ávila-Ríos; Selene Zárate; Susana Lopez; Verónica Mata-Haro; Victor Eduardo García-Arias; Víctor Hugo Borja-Aburto |
| EPI_ISL_3434753, EPI_ISL_3434754, EPI_ISL_3434756, EPI_ISL_3434757 | Laboratorio Central de Saude Publica do Estado Maranhao (LACEN-MA) | Laboratory of Respiratory Viruses and Measles, Oswaldo Cruz Institute, FIOCRUZ | Alice Sampaio Rocha; Ana Carolina Mendonca; Anna Carolina Paixao; Elisa Cavalcante Pereira; Fernando Motta; Lidio Gonçalves Lima Neto; Luciana Appolinario; Marilda Siqueira on behalf of the Fiocruz COVID-19 Genomic Surveillance Network; Paola Resende; Renata Serrano Lopes; Taina Venas |
| EPI_ISL_3235256, EPI_ISL_3434771, EPI_ISL_3434781, EPI_ISL_3827897 | Laboratorio Central de Saude Publica do Estado da Paraiba (LACEN/PB) | Laboratory of Respiratory Viruses and Measles, Oswaldo Cruz Institute, FIOCRUZ | Alice Sampaio Rocha; Ana Carolina Mendonca; Anna Carolina Paixao; Dalane Loudal Florentino Teixeira; Elisa Cavalcante Pereira; Fernando Motta; Joao Felipe Bezerra; Luciana Appolinario; Marilda Siqueira on behalf of the Fiocruz COVID-19 Genomic Surveillance Network; Paola Resende; Renata Serrano Lopes; Taina Venas |
| EPI_ISL_3190309, EPI_ISL_3190330, EPI_ISL_3434830, EPI_ISL_3434837, EPI_ISL_3434849, EPI_ISL_3802912, EPI_ISL_3802920, EPI_ISL_3802934, EPI_ISL_3802935, EPI_ISL_3802949, EPI_ISL_3802952 | | | |
| see above | Laboratorio Central de Saude Publica do Estado de Alagoas (LACEN/AL) | Laboratory of Respiratory Viruses and Measles, Oswaldo Cruz Institute, FIOCRUZ | Agatha Cristinne Prudencio; Agatha Soares; Alice Sampaio Rocha; Ana Carolina Mendonca; Anderson Brandao Leite; Anna Carolina Paixao; Elisa Cavalcante Pereira; Fernando Motta; Ighor Arantes; Ighor Leonardo Arantes Gomes; Luciana Appolinario; Marilda Siqueira on behalf of the Fiocruz COVID-19 Genomic Surveillance Network; Paola Resende; Renata Serrano Lopes; Taina Venas |
| EPI_ISL_3434791, EPI_ISL_3434795, EPI_ISL_3434798, EPI_ISL_3434799, EPI_ISL_3827907, EPI_ISL_3827974 | Laboratorio Central de Saude Publica do Estado de Santa Catarina (LACEN/SC) | Laboratory of Respiratory Viruses and Measles, Oswaldo Cruz Institute, FIOCRUZ | Alice Sampaio Rocha; Ana Carolina Mendonca; Anna Carolina Paixao; Darcita Buerger Rovaris; Elisa Cavalcante Pereira; Fernando Motta; Luciana Appolinario; Marilda Siqueira on behalf of the Fiocruz COVID-19 Genomic Surveillance Network; Paola Resende; Renata Serrano Lopes; Sandra Bianchini Fernandes; Taina Venas |
| EPI_ISL_3803005, EPI_ISL_3803013, EPI_ISL_3803017 | Laboratorio Central de Saude Publica do Estado de Sergipe (LACEN/SE) | Laboratory of Respiratory Viruses and Measles, Oswaldo Cruz Institute, FIOCRUZ | Agatha Soares; Alice Sampaio Rocha; Ana Carolina Mendonca; Anna Carolina Paixao; Cliomar Alves dos Santos; Elisa Cavalcante Pereira; Fernando Motta; Ighor Arantes; Luciana Appolinario; Marilda Siqueira on behalf of the Fiocruz COVID-19 Genomic Surveillance Network; Paola Resende; Renata Serrano Lopes; Tainá Moreira Martins Venas |
| EPI_ISL_3434982, EPI_ISL_3434983, EPI_ISL_3539749, EPI_ISL_3539932 | Laboratorio Central de Saude Publica do Estado do Amapa (LACEN/AP) | Laboratory of Respiratory Viruses and Measles, Oswaldo Cruz Institute, FIOCRUZ | Agatha Cristinne Prudencio; Alice Sampaio Rocha; Ana Carolina Mendonca; Andreia Santos Costa; Anna Carolina Paixao; Anne Caroline da Silva Soledade; Elisa Cavalcante Pereira; Fernando Motta; Ighor Leonardo Arantes Gomes; Lindomar dos Anjos Silva; Luciana Appolinario; Marcia Socorro Pereira Cavalcante; Marilda Siqueira on behalf of the Fiocruz COVID-19 Genomic Surveillance Network; Paola Resende; Renata Serrano Lopes; Taina Venas |
| EPI_ISL_3434742, EPI_ISL_3434743, EPI_ISL_3434750, EPI_ISL_3435042, EPI_ISL_3435043, EPI_ISL_3435044, EPI_ISL_3435045, EPI_ISL_3435046, EPI_ISL_3435047, EPI_ISL_3827857, EPI_ISL_3827900, EPI_ISL_3827932, EPI_ISL_3827996 | | | |
| see above | Laboratorio Central de Saude Publica do Estado do Espirito Santo (LACEN/ES) | Laboratory of Respiratory Viruses and Measles, Oswaldo Cruz Institute, FIOCRUZ | Alice Sampaio Rocha; Ana Carolina Mendonca; Anna Carolina Paixao; Elisa Cavalcante Pereira; Fernando Motta; Luciana Appolinario; Marilda Siqueira on behalf of the Fiocruz COVID-19 Genomic Surveillance Network; Paola Resende; Renata Serrano Lopes; Rodrigo Ribeiro Rodrigues; Taina Venas |
| EPI_ISL_3190262, EPI_ISL_3190263 | Laboratorio Central de Saude Publica do Estado do Maranhao (LACEN-MA) | Laboratory of Respiratory Viruses and Measles, Oswaldo Cruz Institute, FIOCRUZ | Agatha Soares; Alice Sampaio Rocha; Ana Carolina Mendonca; Anna Carolina Paixao; Elisa Cavalcante Pereira; Fernando Motta; Ighor Arantes; Lidio Gonçalves Lima Neto; Luciana Appolinario; Marilda Siqueira on behalf of the Fiocruz COVID-19 Genomic Surveillance Network; Paola Resende; Renata Serrano Lopes; Tainá Venas |
| EPI_ISL_3190172, EPI_ISL_3190173 | Laboratorio Central de Saude Publica do Estado do Maranhao (LACEN/MA) | Laboratory of Respiratory Viruses and Measles, Oswaldo Cruz Institute, FIOCRUZ | Agatha Soares; Alice Sampaio Rocha; Ana Carolina Mendonca; Anna Carolina Paixao; Elisa Cavalcante Pereira; Fernando Motta; Ighor Arantes; Lidio Gonçalves Lima Neto; Luciana Appolinario; Marilda Siqueira on behalf of the Fiocruz COVID-19 Genomic Surveillance Network; Paola Resende; Renata Serrano Lopes; Tainá Venas |
| EPI_ISL_3913672, EPI_ISL_3913767, EPI_ISL_3913789, EPI_ISL_3913806, EPI_ISL_3913815, EPI_ISL_3913836, EPI_ISL_3913858, EPI_ISL_3913870, EPI_ISL_3913925, EPI_ISL_3914163, EPI_ISL_3914184, EPI_ISL_3914211, EPI_ISL_3914232, EPI_ISL_3914267, EPI_ISL_3914275, EPI_ISL_3914282 | | | |
| see above | Laboratorio Central de Saude Publica do Estado do Parana (Instituto de Biologia | Instituto Carlos Chagas - Fiocruz | Alessandra De Melo Aguiar; Andreia Akemi Suzukawa; Andréa Rodrigues Ávila; Bruno Dallagiovanna; Dalila Zanette; Eduardo Balsanelli; Emanuel Maltempi de Souza; Fabio Passetti; Fabricio Klerynton Marchini; Fábio de Oliveira Pedrosa; Guilherme Becker; Helisson Faoro; Hellen Geremias dos Santos; Irina Nastassja Riediger; Letusa Albrecht; Lucas Blanes; Luis Gustavo Morello; Lysangela Ronalte Alves; Maria do Carmo Debur; Mauro de Medeiros Oliveira; Michelle Orane Schemberger; Paola Cristina Resende; Sheila Cristina Nardeli; Tiago Gräf; Valter Antônio de Baura |

| | | | |
|---|---|---|---|
| | Molecular do Paraná (LACEN-PR) | | |
| EPI_ISL_2983432, EPI_ISL_3190368, EPI_ISL_3190370, EPI_ISL_3190386, EPI_ISL_3832475 | Laboratorio Central de Saude Publica do Estado do Parana (LACEN/PR) | Laboratory of Respiratory Viruses and Measles, Oswaldo Cruz Institute, FIOCRUZ | Agatha Soares; Alice Sampaio Rocha; Ana Carolina Mendonca; Anna Carolina Paixao; Elisa Cavalcante Pereira; Fernando Motta; Ighor Arantes; Ighor Leonardo Arantes; Irina Riediger; Luciana Appolinario; Marilda Siqueira on behalf of the Fiocruz COVID-19 Genomic Surveillance Network; Paola Resende; Renata Serrano Lopes; Taina Venas |
| EPI_ISL_3539838, EPI_ISL_3832405 | Laboratorio Central de Saude Publica do Estado do Rio Grande do Norte (LACEN/RN) | Laboratory of Respiratory Viruses and Measles, Oswaldo Cruz Institute, FIOCRUZ | Agatha Soares; Alice Sampaio Rocha; Ana Carolina Mendonca; Ana Paula Ferreira Costa; Anna Carolina Paixao; Antonnyo Palmielly Diogenes Lima; Aurélio de Oliveira Bento; Elisa Cavalcante Pereira; Fernando Motta; Gessika Brenna Costa Alves; Heglayne Pereira Vital da Silva; Iago de Souza Gomes; Ighor Arantes; Isabelle Cristina Clemente dos Santos; Janaina Sonale Cavalcante Nogueira de Oliveira; Jayra Juliana Paiva Alves Abrantes; Jonas José da Silva; Luciana Appolinario; Marilda Siqueira on behalf of the Fiocruz COVID-19 Genomic Surveillance Network; Paola Resende; Renata Serrano Lopes; Taina Venas; Themis Rocha de Souza; Vítor Gabriel Saldanha Fernandes |
| EPI_ISL_2982804, EPI_ISL_3832395, EPI_ISL_3832467 | Laboratorio Central de Saude Publica do Estado do Rio Grande do Sul (LACEN-RS) | Laboratory of Respiratory Viruses and Measles, Oswaldo Cruz Institute, FIOCRUZ | Agatha Cristinne Soares; Alice Sampaio Rocha; Ana Carolina Mendonca; Anderson Brandao Leite; Anna Carolina Paixao; Elisa Cavalcante Pereira; Fernando Motta; Ighor Leonardo Arantes; Luciana Appolinario; Marilda Siqueira on behalf of the Fiocruz COVID-19 Genomic Surveillance Network; Paola Resende; Renata Serrano Lopes; Taina Venas |
| EPI_ISL_3235310, EPI_ISL_3235311, EPI_ISL_3235312, EPI_ISL_3235314 | Laboratorio Central de Saude Publica do Estado do Tocantins (LACEN/TO) | Laboratory of Respiratory Viruses and Measles, Oswaldo Cruz Institute, FIOCRUZ | Agatha Soares; Alice Sampaio Rocha; Ana Carolina Mendonca; Anna Carolina Paixao; Elisa Cavalcante Pereira; Fernando Motta; Ighor Arantes; Jucimaria Dantas Galvao; Luciana Appolinario; Marilda Siqueira on behalf of the Fiocruz COVID-19 Genomic Surveillance Network; Paola Resende; Renata Serrano Lopes; Taina Venas |
| EPI_ISL_3260098 | Laboratorio Central de la Ciudad de Santa Fe | Grupo de Genómica y Bioinformática del Instituto de Investigación de la Cadena Láctea CONICET-INTA on behalf of 'Proyecto Argentino Interinstitucional de genomica de SARS-CoV-2' (PAIS Consortium) | AF; Amadio; C; Eberhardt; G; Irazoqui; JM; MF; Mugna; Ojeda; Pastor; Rompato; V |
| EPI_ISL_3050574, EPI_ISL_3050616 | Laboratorio de Ecologia de Doencas Transmissiveis na Amazonia, Instituto Leonidas e Maria Deane - Fiocruz Amazonia | Laboratorio de Ecologia de Doencas Transmissiveis na Amazonia, Instituto Leonidas e Maria Deane - Fiocruz Amazonia | André Corado; Felipe Naveca; Fernanda Nascimento; George Silva; Karina Pessoa; Luciana Gonçalves; Maria Júlia Brandão; Matilde Mejía; Valdinete Nascimento; Victor Souza; Ágatha Costa |
| EPI_ISL_3707440 | Laboratorio de Genómica Microbiana, Universidad Peruana Cayetano Heredia | cov0937 | Alejandra Dávila-Barclay; Diego Cuicapuza; Guillermo Salvatierra; Janet Huancachoque; Luis González; Pablo Tsukayama; Pedro E. Romero; Pool Marcos |
| EPI_ISL_3049022, EPI_ISL_3761521, EPI_ISL_3761527, EPI_ISL_3761530, EPI_ISL_3761535, EPI_ISL_3761547, EPI_ISL_3761550, EPI_ISL_3761554, EPI_ISL_3761557, EPI_ISL_3761565, EPI_ISL_3761577, EPI_ISL_3761578, EPI_ISL_3761590, EPI_ISL_3761593, EPI_ISL_3761603, EPI_ISL_3761604, EPI_ISL_3761608, EPI_ISL_3761618, EPI_ISL_3761623, EPI_ISL_3761629, EPI_ISL_3761631, EPI_ISL_3761633, EPI_ISL_3761635, EPI_ISL_3761658, EPI_ISL_3761660, EPI_ISL_3761692, EPI_ISL_3761698, EPI_ISL_3761702, EPI_ISL_3761716, EPI_ISL_3761765, EPI_ISL_3761766 | | | |
| see above | Laboratorio de Pesquisa em Virologia, FAMERP, SJRP | Laboratorio de Pesquisa em Virologia, FAMERP, SJRP | Beatriz de Carvalho Marques; Cecília Artico Banho; Cíntia Bittar; Fábio Sossai Possebon; Guilherme Campos; Helena Lage Ferreira; Jorge A. Petroli Marchesi; João Pessoa Araújo Jr.; Leila Sabrina Ullmann; Lívia Sacchetto; Maisa C. Pereira Parra; Marília Moraes; Maurício L. Nogueira.; Paula Rahal; Paulo Inacio da Costa |
| EPI_ISL_3672407, EPI_ISL_3672408, EPI_ISL_3672424, EPI_ISL_3672426, EPI_ISL_3672429, EPI_ISL_3672431, EPI_ISL_3826386, EPI_ISL_3826387, EPI_ISL_3826390, EPI_ISL_3826391, EPI_ISL_3826414, EPI_ISL_3826419, EPI_ISL_3826421, EPI_ISL_3826423, EPI_ISL_3826424, EPI_ISL_3826428, EPI_ISL_3826435, EPI_ISL_3826438, EPI_ISL_3826442 | | | |
| see above | Laboratorio de Referencia Nacional de Virus Respiratorios. Centro Nacional de Salud Publica. Instituto Nacional de Salud Peru. | Laboratorio de Referencia Nacional de Virus Respiratorios. Centro Nacional de Salud Publica. Instituto Nacional de Salud Peru. | Carlos Padilla Rojas; Henri Bailon Calderon; Iris Silva Molina; Joseph Huayra Niquen; Lely Solari Zerpa; Luis Barcena Flores; Marco Galarza Perez; Nancy Rojas Serrano; Nieves Sevilla Castañeda; Omar Caceres Rey; Orson Mestanza Millones; Princesa Medrano Alhuay; Priscila Lope Pari; Sandra Morales Ruiz; Sara Gordillo Vilchez; Steve Acedo Lazo; Veronica Hurtado Vela; Victor Jimenez Vasquez; Wendy Lizarraga Olivares |
| EPI_ISL_2765554, EPI_ISL_2975218, EPI_ISL_2975220, EPI_ISL_3026010, EPI_ISL_3128484, EPI_ISL_3128491, EPI_ISL_3128492 | | | |
| see above | Laboratorio di Riferimento Regionale della Sicilia Occidentale per l'Emergenza COVID-19 | Laboratorio di Riferimento Regionale della Sicilia Occidentale per l'Emergenza COVID-19 | Carmelo Massimo Maida; Claudio Costantino; Daniela Di Naro; Fabio Tramuto; Francesco Vitale; Giorgio Graziano; Giulia Randazzo; Vincenzo Restivo; Walter Mazzucco |
| EPI_ISL_2611338, EPI_ISL_2872328, EPI_ISL_2926732, EPI_ISL_2926733, EPI_ISL_3017581, EPI_ISL_3111969, EPI_ISL_3112892, EPI_ISL_3113387, EPI_ISL_3326419, EPI_ISL_3326531, EPI_ISL_3329086, EPI_ISL_3511781, EPI_ISL_3514551, EPI_ISL_3518963, EPI_ISL_3687444, EPI_ISL_3687994, EPI_ISL_3747707 | | | |
| see above | Laboratory Corporation of America | Centers for Disease Control and Prevention Division of Viral Diseases, Pathogen Discovery | Adrian Paskey; Amanda Douglas; Amanda Suchanek; Andrea Throop; Ayla Burns; Benjamin Rambo-Martin; Bobbi Croy; Brian Krueger; Brian Norvell; Christopher Gulvick; Christos Petropoulos; Clinton Paden; Clinton R. Paden; Craig Lukasik; Dakota Howard; Darlene Wagner; Debbie Boles; Dhwani Batra; Duncan MacCannell; Eyad Almasri; Goran Stevovic; Howard Engler; Hrushikesh Deshmukh; Jake Humphrey; Jana Schroth; Jason Caravas; Joe Voshell; John Pruitt; Jonathan Meltzer; Jonathan Williams; Kara Moser; Kimberly Wagner; Lax Iyer; Lisa Pfefferle; Lyndon Tilson; Manoj Jain; Marcia Eisenberg; Mary Ann Cristobal; Mary Cristobal; Mary Williamson; Matthew Robinson; Matthew Schmerer; Michael Levandoski; Mike Sapeta; Mindy Nye; Minoo Agarwal; Mohan Kolli; Nuthawin Charoensri; Oren Cohen; Peter Cook; Peter W. Cook; Prashant Gupta; Qian Zeng; Rama Ghatti; Scott Parker; Scott Ryan; Scott Sammons; Shatavia Morrison; Stanley Letovsky; Steven Ragan; Suresh Babu Selvaraju; Suresh Selvaraju; Susan Countryman; Susan Hicks; Suzanne Dale; Thomas Urban; Tim Kuphal; Tricia Zwiefelhofer; Vincent Drouillon; Yvette Unoarumhi |
| EPI_ISL_3050310, EPI_ISL_3050313, EPI_ISL_3050314, EPI_ISL_3050320, EPI_ISL_3050321, EPI_ISL_3050324, EPI_ISL_3050330, EPI_ISL_3050333, EPI_ISL_3050335, EPI_ISL_3050336, EPI_ISL_3050339, EPI_ISL_3050340, EPI_ISL_3050343, EPI_ISL_3050345, EPI_ISL_3050347, EPI_ISL_3050348, EPI_ISL_3050355, EPI_ISL_3050360, EPI_ISL_3050364, EPI_ISL_3050366, EPI_ISL_3050372, EPI_ISL_3050373, EPI_ISL_3050374, EPI_ISL_3050377, EPI_ISL_3050378, EPI_ISL_3050382, EPI_ISL_3050386, EPI_ISL_3050389, EPI_ISL_3050398, EPI_ISL_3050399, EPI_ISL_3050400, EPI_ISL_3050405, EPI_ISL_3050410, EPI_ISL_3050411, EPI_ISL_3050412, EPI_ISL_3050413, EPI_ISL_3050414, EPI_ISL_3050417, EPI_ISL_3050419, EPI_ISL_3050422, EPI_ISL_3050425, EPI_ISL_3050426, EPI_ISL_3050427, EPI_ISL_3050428, EPI_ISL_3050429, EPI_ISL_3050430, EPI_ISL_3050433, EPI_ISL_3050439, EPI_ISL_3050443, EPI_ISL_3050444, EPI_ISL_3050445, EPI_ISL_3050446, EPI_ISL_3050447, EPI_ISL_3050448, EPI_ISL_3050449, EPI_ISL_3050450, EPI_ISL_3050451, EPI_ISL_3050458, EPI_ISL_3050465, EPI_ISL_3050466, EPI_ISL_3050473, EPI_ISL_3050476, EPI_ISL_3050477, EPI_ISL_3050479, EPI_ISL_3050488, EPI_ISL_3050492, EPI_ISL_3050504, EPI_ISL_3050507, EPI_ISL_3050508, EPI_ISL_3050512, EPI_ISL_3050513, EPI_ISL_3050520, EPI_ISL_3050526, EPI_ISL_3050532, EPI_ISL_3050534, EPI_ISL_3050535, EPI_ISL_3050539, EPI_ISL_3050541, EPI_ISL_3050542, EPI_ISL_3050546, EPI_ISL_3050548, EPI_ISL_3050549, EPI_ISL_3050550, EPI_ISL_3050553, EPI_ISL_3050555, EPI_ISL_3050556, EPI_ISL_3050562, EPI_ISL_3050563, EPI_ISL_3050564, EPI_ISL_3050567, EPI_ISL_3050568, EPI_ISL_3050572, EPI_ISL_3050573, EPI_ISL_3050576, EPI_ISL_3050577, EPI_ISL_3050578, EPI_ISL_3050579, EPI_ISL_3050583, EPI_ISL_3050585, EPI_ISL_3050596, EPI_ISL_3050598, EPI_ISL_3050610, EPI_ISL_3050623, EPI_ISL_3050635, EPI_ISL_3050647, EPI_ISL_3050648, EPI_ISL_3050650 | | | |
| see above | Laboratório Central de Saúde Pública do Amazonas - LACEN-AM | Laboratorio de Ecologia de Doencas Transmissiveis na Amazonia, Instituto Leonidas e Maria Deane - Fiocruz Amazonia | André Corado; Felipe Naveca; Fernanda Nascimento; George Silva; Karina Pessoa; Luciana Gonçalves; Maria Júlia Brandão; Matilde Mejía; Valdinete Nascimento; Victor Souza; Ágatha Costa |
| EPI_ISL_3610357 | Laboratório Central de Saúde Pública do Estado do Tocantins (LACEN/TO) | Bioinformatics and Biotechnology Laboratory (Labinftec/Federal University of Tocantins, Campus of Gurupi) | Bergmann Morais Ribeiro; Fabrício Souza Campos; Fernando Lucas Melo; Jucimária Dantas Galvão; Raíssa Nunes dos Santos; Ueric José Borges de Souza |
| EPI_ISL_3539791, EPI_ISL_3539792, EPI_ISL_3539793 | Laboratório de Biotecnologia Aplicada (LBA) - Laboratório de Biologia Molecular - Hospital das Clínicas, Faculdade de Medicina de Botucatu, Departamento de Bioprocessos e Biotecnologia - Faculdade de Ciências Agronômicas, UNESP – Botucatu/SP | Laboratory of Respiratory Viruses and Measles, Oswaldo Cruz Institute, FIOCRUZ | Alice Sampaio Rocha; Ana Carolina Mendonca; Anna Carolina Paixao; Elisa Cavalcante Pereira; Felipe Allan da Silva Costa; Fernando Motta; Jayme Augusto de Souza Neto; Leonardo Nazario de Moraes; Luciana Appolinario; Marilda Siqueira on behalf of the Fiocruz COVID-19 Genomic Surveillance Network; Paola Resende; Patricia Akemi Assato; Rejane Maria Tommasini; Renata Serrano Lopes; Taina Venas |
| EPI_ISL_3456065, EPI_ISL_3461099, EPI_ISL_3492497, EPI_ISL_3492499, EPI_ISL_3492512, EPI_ISL_3806780 | Laboratório de Microbiologia Molecular - Universidade FEEVALE | Molecular Microbiology Laboratory | Alana Witt Hansen; Fernando Rosado Spilki; Fágner Henrique Heldt; Juliana Schons Gulart; Juliana Schons Gularte; Juliane Deise Fleck; Mariana Soares da Silva; Matheus Nunes Weber; Meriane Demoliner; Michele Filippi.; Micheli Filippi; Paula Rodrigues de Almeida; Vyctoria Malayhka de Abreu Góes Pereira. |
| EPI_ISL_2863595, EPI_ISL_2863598, EPI_ISL_2983228, EPI_ISL_2983229, EPI_ISL_2983416, EPI_ISL_2983417, EPI_ISL_2983421, EPI_ISL_3190409, EPI_ISL_3190410, EPI_ISL_3190411 | | | |
| see above | Labortorio Central de Saude Publica do Estado do Tocantins (LACEN/TO) | Laboratory of Respiratory Viruses and Measles, Oswaldo Cruz Institute, FIOCRUZ | Agatha Soares; Alice Sampaio Rocha; Ana Carolina Mendonca; Anna Carolina Paixao; Elisa Cavalcante Pereira; Fernando Motta; Ighor Arantes; Jucimaria Dantas Galvao; Luciana Appolinario; Marilda Siqueira on behalf of the Fiocruz COVID-19 Genomic Surveillance Network; Paola Resende; Renata Serrano Lopes; Taina Venas |
| EPI_ISL_2809112 | Lighthouse Lab in Milton Keynes | Wellcome Sanger Institute for the COVID-19 Genomics UK (COG-UK) Consortium | Cordelia Langford; David K. Jackson; Dominic Kwiatkowski; Ewan Harrison; Ian Johnston; Jeffrey Barrett; John Sillitoe on behalf of the Wellcome Sanger Institute COVID-19 Surveillance Team; Roberto Amato; Sonia Goncalves; The Lighthouse Lab in Milton Keynes and Alex Alderton |
| EPI_ISL_3912101, EPI_ISL_3912235, EPI_ISL_3912237 | MATERNIDADE ESCOLA ASSIS CHATEAUBRIAND | ACME Lab, Oswaldo Cruz Foundation, FIOCRUZ/CE | Cleber Furtado Aksenen; Fabio Miyajima; Fernando Braga Stehling; Francisco Eder de Moura Lopes; Jamille Maria Mendes Bezerra; Joaquim Cesar do Nascimento Sousa Junior; Pedro Miguel Carneiro Jeronimo; Suzana Porto Almeida & Lucas Delerino on behalf of COVID-19 FIOCRUZ Genomic Network; Thais Ferreira de Oliveira; Thais de Oliveira Costa; Ticiane Cavalcante de Souza; Veridiana Pessoa Miyajima |
| EPI_ISL_3491530, EPI_ISL_3491531 | MD DOH Laboratories Administration | Centers for Disease Control and Prevention Division of Viral Diseases, Pathogen Discovery | Alex Burgin; Ben Rambo-Martin; Clinton Paden; Dakota Howard; Dave Wentworth; Dhwani Batra; Jasmine Padilla; Justin Lee; Krista Queen; Kristen Knipe; Kristine Lacek; Mark Burroughs; Matthew Schmerer; Meghan Bentz; Mili Sheth; Peter Cook; Sam Shepard; Sarah Nobles; Suxiang Tong; Vivien Dugan; Yvette Unoarumhi |
| EPI_ISL_3152288, EPI_ISL_3152291, EPI_ISL_3235571, EPI_ISL_3235641 | MD PHL | Maryland Department of Health Laboratories Administration | Ami Patel; Eric N. Keller; Jillian Loomis; Kwang Low; Terence L. Moore; and Robert Myers |
| EPI_ISL_3355449 | MI - Michigan Department of Health and Human Services - Bureau of Laboratories | Centers for Disease Control and Prevention Division of Viral Diseases, Pathogen Discovery | Alex Burgin; Ben Rambo-Martin; Clinton Paden; Dakota Howard; Dave Wentworth; Dhwani Batra; Jasmine Padilla; Justin Lee; Krista Queen; Kristen Knipe; Kristine Lacek; Mark Burroughs; Matthew Schmerer; Meghan Bentz; Mili Sheth; Peter Cook; Sam Shepard; Sarah Nobles; Suxiang Tong; Vivien Dugan; Yvette Unoarumhi |
| EPI_ISL_3549791 | Maryland Genomics, Institute for Genome Sciences, University of Maryland School of Medicine | Maryland Genomics, Institute for Genome Sciences, University of Maryland School of Medicine | Claire M; Fraser; Hazen; Holly; Humphrys; Ivette; Jacques; Jonathan; Kranthi; Lim; Lisa D; Luke J; Mike; Ott; Ravel; Roussey; Sadzewicz; Sandra; Santana-Cruz; Tallon; Tracy; Vavikolanu |
| EPI_ISL_3877468 | Medizinisch-Diagnostisches Labor Kempten | Robert Koch Institute | |

| | | | |
|---|---|---|---|
| | allgäulab | | |
| EPI_ISL_3713461 | Medizinische Laboratorien Düsseldorf | Robert Koch Institute | |
| EPI_ISL_3030663, EPI_ISL_3908105 | Microbiology Department & Molecular Biology CORE \| CDB \| Hospital Clinic \| Barcelona | Microbiology Department & Molecular Biology CORE \| CDB \| Hospital Clinic \| Barcelona | Andrea Vergara Gomez; Andrea Vergara Gomez \| Jose Luis Villanueva Canas \| Miguel Julian Martinez Yoldi \| M. Mar Mosquera Gutierrez \| M. Angeles Marcos Maeso; Jose Luis Villanueva Cañas; Miguel Julian Martínez Yoldi; Mª Angeles Marcos Maeso; Mª del Mar Mosquera Gutierrez |
| EPI_ISL_2718555, EPI_ISL_2718571, EPI_ISL_2718604, EPI_ISL_2812448, EPI_ISL_2812455, EPI_ISL_2812464, EPI_ISL_2812499, EPI_ISL_2812522, EPI_ISL_2812531, EPI_ISL_2812534, EPI_ISL_2832140, EPI_ISL_2832177, EPI_ISL_2832189, EPI_ISL_2832201, EPI_ISL_2897044, EPI_ISL_2897048, EPI_ISL_2926114, EPI_ISL_2926152, EPI_ISL_3320176, EPI_ISL_3320286, EPI_ISL_3320298 | | | |
| see above | Microbiology Department, Laboratori Clínic Metropolitana Nord. Hospital Universitari Germans Trias i Pujol | Can Ruti SARS-CoV-2 Sequencing Hub (HUGTiP/IrsiCaixa/IGTP) | Alba Sánchez; Alexia París; Anna Not; Antoni E Bordoy; Bonaventura Clotet; Cristina Casañ; Cristina Esteban; David Panisello; Francesc Catala-Moll; Gemma Clara; Ignacio Blanco; Laia Soler; Lauro Sumoy; Marc Noguera-Julian; Maria Casadellà; Mariona Parera; Mercedes Guerrero; Montserrat Giménez; Pere-Joan Cardona; Pilar Armengol; Roger Paredes; Verónica Saludes; and Elisa Martró on behalf of the Can Ruti SARS-CoV-2 Sequencing Hub |
| EPI_ISL_3216654, EPI_ISL_3216657, EPI_ISL_3216658, EPI_ISL_3260231, EPI_ISL_3260239, EPI_ISL_3260245, EPI_ISL_3260247, EPI_ISL_3260249, EPI_ISL_3260250, EPI_ISL_3448027, EPI_ISL_3459457, EPI_ISL_3803928, EPI_ISL_3803938, EPI_ISL_3803944, EPI_ISL_3803945 | | | |
| see above | Minnesota Department of Health, Public Health Laboratory | Minnesota Department of Health, Public Health Laboratory | Alexandra Lorentz; Jacob Garfin; Matt Plumb; and Xiong Wang |
| EPI_ISL_3317920 | NJDOH, Public Health and Environmental Laboratories | NJ_PHEL | Allison Roder; Byeong Jeong; Chelsea San Filippo; Dana Woell; Jacquelyn Deverell; Lindsey Bodnar; Ryan Pachucki; Shiv K. Verma |
| EPI_ISL_3134777, EPI_ISL_3134782, EPI_ISL_3134809, EPI_ISL_3134810, EPI_ISL_3134811, EPI_ISL_3134812, EPI_ISL_3447413, EPI_ISL_3447499, EPI_ISL_3447509, EPI_ISL_3447520, EPI_ISL_3447532, EPI_ISL_3447544, EPI_ISL_3703820, EPI_ISL_3703896, EPI_ISL_3703905, EPI_ISL_3703909, EPI_ISL_3703965, EPI_ISL_3704090, EPI_ISL_3704174, EPI_ISL_3704178, EPI_ISL_3704192, EPI_ISL_3704212, EPI_ISL_3704232, EPI_ISL_3704392, EPI_ISL_3704404, EPI_ISL_3704405, EPI_ISL_3835345 | | | |
| see above | NUPIT/UFPE | WallauLab on behalf of Fiocruz COVID-19 Genomic Surveillance Network | Alexandre Freitas da Silva; Cassia Docena; Constância Flávia Junqueira Ayres; Filipe Zimmer Dezordi; Gabriel Luz Wallau; Gustavo Barbosa de Lima; Lais Ceschini Machado; Lilian Caroliny Amorim Silva; Maira Galdino da Rocha Pitta; Marcelo Henrique dos Santos Paiva; Matheus Filgueira Bezerra; Michelly Cristiny Pereira; Rômulo Pessoa e Silva; Sinval Pinto Brandão Filho |
| EPI_ISL_3058135, EPI_ISL_3058136 | Nastavni zavod za javno zdravstvo Splitsko- Dalmatinske Županije | Hrvatski zavod za javno zdravstvo | Irena Tabain; Ivana Ferenčak |
| EPI_ISL_2600834 | OSPEDALE CIVILE TERAMO | Istituto Zooprofilattico Sperimentale dell'Abruzzo e Molise "G. Caporale" | Ancora M; Calistri P; Cammà C; Caporale M; Curini V; Delli Compagni E; Di Domenico M; Di Lollo Valeria; Di Pasquale A; Lorusso A; Mangone I; Marcacci M; Puglia I; Rinaldi A; Savini G; Scialabba S |
| EPI_ISL_3189593 | Orebro University Hospital, Dept Laboratory Medicine, Clinical Microbiology | Orebro University Hospital | Sundqvist M et al |
| EPI_ISL_3536146 | POSTO DE SAUDE DE SANTA FELICIA | Oswaldo Cruz Institute, FIOCRUZ/CE | Cleber Furtado Aksenen; Fabio Miyajima; Fernando Braga Stehling; Francisco Eder de Moura Lopes; Jamille Maria Mendes Bezerra; Joaquim César do Nascimento Sousa Junior; Pedro Miguel Carneiro Jeronimo; Suzana Porto Almeida e Lucas Delerino; Thais Ferreira de Oliveira; Thais de Oliveira Costa; Ticiane Cavalcante de Souza; Veridiana Pessoa Miyajima |
| EPI_ISL_3922275, EPI_ISL_3922298 | PRONTO SOCORRO MUNICIPAL TAMBAU | Instituto Butantan | Antonio Jorge Martins; Claudia Renata dos Santos Barros; David Schlesinger; Debora Botequio Moretti; Dimas Tadeu Covas; Elaine Cristina Marqueze; Elaine Vieira Santos; Evandra Strazza Rodrigues; Heidge Fukumasu; Jayme Augusto de Souza-Neto; José Salvatore Leister Patané; Luiz Alcantara; Luiz Lehmann Coutinho; Maria Carolina Elias; Maurício Lacerda Nogueira; Rafael dos Santos Bezerra; Raul Machado Neto; Rejane Maria Tommasini Grotto; Ricardo Haddad; Sandra Coccuzzo Sampaio Vessoni; Simone Kashima; Svetoslav Nanev Slavov; Vincent Louis Viala |
| EPI_ISL_3394308 | Pandemic Response Lab - NYC | Pandemic Response Lab, R&D | Alex Carpio; Cybill del Castillo; Dylan Law; Haiping Hao; Henry Lee; Isabel Fernandez Escapa; Jon Laurent; Melissa Hopkins; Michael Hammerling; Pradeep Bugga; Shinyoung Clair Kang; Sol Rey; William Ward |
| EPI_ISL_3448507, EPI_ISL_3448545 | Public Health Ontario Laboratory | Public Health Ontario Laboratory | Aimin Li; Alireza Eshaghi; Andre Villegas; Ashleigh Sullivan; Christine Frantz; Dean Maxwell; Esha Joshi; Jared Simpson; Jennifer L Guthrie; Jonathan B Gubbay; Karthikeyan Sivaraman; Lawrence Heisler; Matthew Watson; Michael CY Li; Michael Laszloffy; Nahuel Fittipaldi; Philip Banh; Richard de Borja; Samir N Patel; Sandeep Nagra; Sandra Zittermann; Sarah Teatero; Vanessa G Allen; Yao Chen; Yogi Sundaravadanam |
| EPI_ISL_2649977, EPI_ISL_3085834, EPI_ISL_3303877, EPI_ISL_3460812 | Quest Diagnostics Incorporated | Centers for Disease Control and Prevention Division of Viral Diseases, Pathogen Discovery | A. Gerasimova; A. Perez; Adrian Paskey; B. Anderson; Benjamin Rambo-Martin; Christopher Gulvick; Clinton R. Paden; Dakota Howard; Darlene Wagner; Dhwani Batra; Duncan MacCannell; F. Lacbawan; I. A. Shlyakhter; Jason Caravas; K.E. Livingston; Kara Moser; L.E. Bernstein; M. Hua; Matthew Schmerer; P. Tanpaiboon; Peter W. Cook; R. M. Kagan; R. Owen; R. V. Rolando; S. H. Rosenthal; Scott Sammons; Shatavia Morrison; Y. Liu; Yvette Unoarumhi |
| EPI_ISL_3216707 | Regions Hospital | Minnesota Department of Health, Public Health Laboratory | Alexandra Lorentz; Jacob Garfin; Matt Plumb; and Xiong Wang |
| EPI_ISL_2904392 | Respiratory Virus Unit, Microbiology Services Colindale, Public Health England | COVID-19 Genomics UK (COG-UK) Consortium | PHE Covid Sequencing Team |
| EPI_ISL_3912240, EPI_ISL_3912242 | SECRETARIA MUNICIPAL DE SAUDE DE TIANGUA | ACME Lab, Oswaldo Cruz Foundation, FIOCRUZ/CE | Cleber Furtado Aksenen; Fabio Miyajima; Fernando Braga Stehling; Francisco Eder de Moura Lopes; Jamille Maria Mendes Bezerra; Joaquim Cesar do Nascimento Sousa Junior; Pedro Miguel Carneiro Jeronimo; Suzana Porto Almeida & Lucas Delerino on behalf of COVID-19 FIOCRUZ Genomic Network; Thais Ferreira de Oliveira; Thais de Oliveira Costa; Ticiane Cavalcante de Souza; Veridiana Pessoa Miyajima |
| EPI_ISL_3391234, EPI_ISL_3393387 | SELAS MEDILYS | Department of Virology, Henri Mondor University Hospital, Assistance Publique Hôpitaux de Paris, Université Paris-Est Créteil, INSERM U955 | Alexandre Soulier; Christophe Rodriguez; Elisabeth Trawinski; Guillaume Gricourt; Jean-Michel Pawlotsky; Melissa N'Debi; Slim Fourati; Vanessa Demontant |
| EPI_ISL_2673672 | SIESP CHIETI - DRIVE IN ORTONA | Istituto Zooprofilattico Sperimentale dell'Abruzzo e Molise "G. Caporale" | Ancora M; Calistri P; Cammà C; Caporale M; Curini V; Delli Compagni E; Di Domenico M; Di Lollo Valeria; Di Pasquale A; Lorusso A; Mangone I; Marcacci M; Puglia I; Rinaldi A; Savini G; Scialabba S |
| EPI_ISL_2600822 | SIESP DIPARTIMENTO DI PREVENZIONE CHIETI | Istituto Zooprofilattico Sperimentale dell'Abruzzo e Molise "G. Caporale" | Ancora M; Calistri P; Cammà C; Caporale M; Curini V; Delli Compagni E; Di Domenico M; Di Lollo Valeria; Di Pasquale A; Lorusso A; Mangone I; Marcacci M; Puglia I; Rinaldi A; Savini G; Scialabba S |
| EPI_ISL_3836178 | STONY BROOK UNIVERSITY HOSPITAL | Wadsworth Center, New York State Department of Health | Alexis Russell; Catharine Prussing; Daryl M. Lamson; Erasmus Schneider; Erica Lasek-Nesselquist; John Kelly; Jonathan Plitnick; Kirsten St. George; Matthew Shudt; Melissa A Leisner; Navjot Singh |
| EPI_ISL_2844964, EPI_ISL_2844971, EPI_ISL_2844982, EPI_ISL_2844991 | SYNLAB MVZ Weiden | Robert Koch Institute | |
| EPI_ISL_2616916, EPI_ISL_2692617, EPI_ISL_2810148, EPI_ISL_2978363 | Salud Digna | Instituto Nacional de Medicina Genomica | Abraham Campos-Romero; Cedro-Tanda A; Escobar-Arrazola; Gonzalez-Barrera D; Herrera-Montalvo LA.; Hidalgo-Miranda A; Luna-Ruiz Marco; M; Mendoza-Vargas A; Moreno-Camacho José Luis; Munguia-Garza P; Ramirez-Vega O; Rangel-DeLeon D; Reyes-Grajeda JP; Rodriguez-Gallegos Jorge |
| EPI_ISL_3373984, EPI_ISL_3373996, EPI_ISL_3374015 | Salud Digna, A.C | Andersen lab at Scripps Research | Abraham Garcia Gil; Jose Luis Moreno Camacho; Marco Antonio Luna Ruiz-Esparza; Miguel A. Fernandez Rojas; SEARCH Alliance with Abraham Campos Romero |
| EPI_ISL_3374121 | San Diego County Public Health Laboratory | Andersen lab at Scripps Research | Brett Austin; Jovan Shephard; SEARCH Alliance San Diego with Ashleigh Murphy |
| EPI_ISL_3503243, EPI_ISL_3708858 | San Diego County Public Health Laboratory | San Diego County Public Health Laboratory | Ashleigh Murphy Schafer; Brett Austin |
| EPI_ISL_3373825 | Sharp HealthCare Laboratory | Andersen lab at Scripps Research | Art Mendoza; Cathy Woerle; Jacquelyn Berumen; Liam McGinnis; Omid Bakhtar; SEARCH Alliance San Diego with Aaron Harding |
| EPI_ISL_3460522 | TGen North | TGen North | "Jolene Bowers; Brett Van Tassel; Chris French; Darrin Lemmer; Dave Engelthaler"; Hayley Yaglom; Heather Centner |
| EPI_ISL_3215225, EPI_ISL_3215284, EPI_ISL_3266303 | U.O. Microbiologia Laboratorio Unico Centro Servizi - AUSL della Romagna | U.O. Microbiologia, Laboratorio Unico Centro Servizi - AUSL della Romagna | Giorgio Dirani |
| EPI_ISL_3536337 | UBS RAFAEL ALVES BEZERRA | Oswaldo Cruz Institute, FIOCRUZ/CE | Cleber Furtado Aksenen; Fabio Miyajima; Fernando Braga Stehling; Francisco Eder de Moura Lopes; Jamille Maria Mendes Bezerra; Joaquim César do Nascimento Sousa Junior; Pedro Miguel Carneiro Jeronimo; Suzana Porto Almeida e Lucas Delerino; Thais Ferreira de Oliveira; Thais de Oliveira Costa; Ticiane Cavalcante de Souza; Veridiana Pessoa Miyajima |
| EPI_ISL_3912318 | UNIDADE DE PRONTO ATENDIMENTO DE EUSEBIO | ACME Lab, Oswaldo Cruz Foundation, FIOCRUZ/CE | Cleber Furtado Aksenen; Fabio Miyajima; Fernando Braga Stehling; Francisco Eder de Moura Lopes; Jamille Maria Mendes Bezerra; Joaquim Cesar do Nascimento Sousa Junior; Pedro Miguel Carneiro Jeronimo; Suzana Porto Almeida & Lucas Delerino on behalf of COVID-19 FIOCRUZ Genomic Network; Thais Ferreira de Oliveira; Thais de Oliveira Costa; Ticiane Cavalcante de Souza; Veridiana Pessoa Miyajima |
| EPI_ISL_3912059, EPI_ISL_3912108, EPI_ISL_3912335, EPI_ISL_3912336, EPI_ISL_3912340, EPI_ISL_3912346, EPI_ISL_3912347, EPI_ISL_3912349, EPI_ISL_3912350, EPI_ISL_3912351, EPI_ISL_3912352, EPI_ISL_3912353, EPI_ISL_3912354, EPI_ISL_3912356, EPI_ISL_3912357, EPI_ISL_3912360, EPI_ISL_3912361, EPI_ISL_3912368, EPI_ISL_3912374, EPI_ISL_3912375, EPI_ISL_3912376 | | | |
| see above | UNIDADE REFERENCIA COVID 19 | ACME Lab, Oswaldo Cruz Foundation, FIOCRUZ/CE | Cleber Furtado Aksenen; Fabio Miyajima; Fernando Braga Stehling; Francisco Eder de Moura Lopes; Jamille Maria Mendes Bezerra; Joaquim Cesar do Nascimento Sousa Junior; Pedro Miguel Carneiro Jeronimo; Suzana Porto Almeida & Lucas Delerino on behalf of COVID-19 FIOCRUZ Genomic Network; Thais Ferreira de Oliveira; Thais de Oliveira Costa; Ticiane Cavalcante de Souza; Veridiana Pessoa Miyajima |
| EPI_ISL_3536199, EPI_ISL_3536200, EPI_ISL_3536202, EPI_ISL_3536203 | UNIDADE REFERENCIA COVID 19 | Oswaldo Cruz Institute, FIOCRUZ/CE | Cleber Furtado Aksenen; Fabio Miyajima; Fernando Braga Stehling; Francisco Eder de Moura Lopes; Jamille Maria Mendes Bezerra; Joaquim César do Nascimento Sousa Junior; Pedro Miguel Carneiro Jeronimo; Suzana Porto Almeida e Lucas Delerino; Thais Ferreira de Oliveira; Thais de Oliveira Costa; Ticiane Cavalcante de Souza; Veridiana Pessoa Miyajima |
| EPI_ISL_2942471 | Unidad de Investigacion Biomedica de Zacatecas (UIBZ) | Unidad de Genomica Avanzada | ; Alejandra Garcia-Gasca; Alejandra Hernandez-Teran; Alejandro Sanchez-Flores; Alfredo Herrera-Estrella; Alicia Ocaña-Mondragon; Andreu Comas-Garcia; Angel Gustavo Salas-Lais; Antonio Loza Roman; Bernardo Martinez-Miguel; Blanca Taboada; Brenda Irasema Maldonado-Meza; Bruno Gomez-Gil; Carla Ivon Herrera-Najera; Carlos F. Arias; Celia Boukadida; Celida Duque Molina; Celida Martinez- Rodriguez; Clara Esperanza Santacruz-Tinoco; Concepcion Grajales-Muñiz; Consorcio Mexicano de Vigilancia Genomica (CoViGen-Mex). Authors (in alphabetical order): Julio Elias Alvarado-Yaah; Cristobal Chaidez-Quiroz; Daniel Fregoso-Rueda; Daniel Lira Morales; Eduardo Becerril-Vargas; Fernando Fontove-Herrera; Fidencio Mejia-Nepomuceno; Francisco Pulido; Gloria Elena Espinosa-Ayala; Gloria Maria Molina-Salinas; Gloria Vazquez; Hector Esteban Paz-Juarez; Hector Montoya-Fuentes; Helen Haydee Fernanda Ramirez-Plascencia; Irvin Gonzalez-Lopez; Jean Pierre Gonzalez; Jesus Hernandez; Joel Armando Vazquez-Perez.; Jorge Salas-Hernandez; Jose Antonio Enciso-Moreno; Jose Arturo Martinez-Orozco; Jose Esteban Muñoz-Medina; Jose de Jesus Nuñez-Contreras; Juan Bautista Chale-Dzul; Julissa Enciso-Ibarra; Luis Alberto Ochoa-Carrera; Margarita Matias-Florentino; Maria Guadalupe Santiago-Mauricio; Maria Guadalupe de Jesus Mireles-Rivera; Mario Mujica-Sanchez; Marissa Perez-Garcia; Nelly Selem-Mojica; Pavel Isa; Ricardo Ciria Merce; Ricardo Grande; Rosa Maria Gutierrez Rios; Santiago avila-Rios; Selene Zarate; Susana Lopez; Veronica Mata-Haro; Victor Eduardo Garcia-Arias; Victor Hugo Borja-Aburto |
| EPI_ISL_2681274, | Unidad de Investigación Médica de | Instituto de Biotecnología de la | ; Alejandra García-Gasca; Alejandra Hernández-Terán; Alejandro Sánchez-Flores; Alfredo Herrera-Estrella; Alicia Ocaña-Mondragón; Andreu Comas-García; Angel Gustavo Salas-Lais; Antonio Loza Román; Bernardo Martínez-Miguel; Blanca Taboada; Brenda Irasema Maldonado-Meza; Bruno Gómez-Gil; |

| | | | |
|---|---|---|---|
| EPI_ISL_2681364 | Yucatán (UIMY) | UNAM | Carla Ivón Herrera-Najera; Carlos F. Arias; Celia Boukadida; Clara Esperanza Santacruz-Tinoco; Concepción Grajales-Muñiz; Consorcio Mexicano de Vigilancia Genómica (CoViGen-Mex). Authors (in alphabetical order): Julio Elias Alvarado-Yaah; Cristóbal Cháidez-Quiróz; Célida Duque Molina; Célida Martínez- Rodríguez; Daniel Fregoso-Rueda; Daniel Lira Morales; Eduardo Becerril-Vargas; Fernando Fontove-Herrera; Fidencio Mejía-Nepomuceno; Francisco Pulido; Gloria Elena Espinosa-Ayala; Gloria María Molina-Salinas; Gloria Vazquez; Hector Esteban Paz-Juárez; Hector Montoya-Fuentes; Helen Haydee Fernanda Ramírez-Plascencia; Irvin González-López; Jean Pierre González; Jesús Hernández; Joel Armando Vázquez-Pérez.; Jorge Salas-Hernández; José Antonio Enciso-Moreno; José Arturo Martínez-Orozco; José Esteban Muñoz-Medina; José de Jesús Nuñez-Contreras; Juan Bautista Chale-Dzul; Julissa Enciso-Ibarra; Luis Alberto Ochoa-Carrera; Margarita Matías-Florentino; Mario Mújica-Sánchez; Marissa Perez-Garcia; María Guadalupe Santiago-Mauricio; María Guadalupe de Jesús Mireles-Rivera; Nelly Sélem-Mojica; Pavel Isa; Ricardo Ciria Merce; Ricardo Grande; Rosa María Gutiérrez Rios; Santiago Ávila-Ríos; Selene Zárate; Susana Lopez; Verónica Mata-Haro; Victor Eduardo García-Arias; Víctor Hugo Borja-Aburto |
| EPI_ISL_2837278, EPI_ISL_3072175, EPI_ISL_3072325, EPI_ISL_3072539, EPI_ISL_3072624, EPI_ISL_3245417, EPI_ISL_3245422, EPI_ISL_3921410 | | | |
| see above | Unidade de apoio ao diagnostico da COVID - UNADIG | Bioinformatics Laboratory / LNCC | Alessandra P Lamarca; Alexandra L Gerber; Amilcar Tanuri; Ana Paula de C Guimaraes; Ana Tereza R Vasconcelos; Andrea Cony Cavalcanti; Caio Luiz Pereira Ribeiro; Cassia Alves; Cintia Policarpo; Claudia Maria Braga de Mello; Cristiane Gomes da Silva; Diana Mariani; Douglas Terra Machado; Erica Ramos dos Santos Nascimento; Fernanda Leitao dos Santos; Flavio Dias da Silva; Gleidson da Silva de Oliveira; Leandro Magalhaes de Souza; Liliane Cavalcante; Luiz G P de Almeida; Marcio Henrique de Oliveira Garcia; Mario Sergio Ribeiro; Ricardo Jose Barbosa Salviano; Ronaldo da Silva F Jr; Silvia Carvalho |
| EPI_ISL_3666911 | Unilabs Ticino (Breganzona) | Laboratorio di Microbiologia | Martinetti Lucchini Gladys; Valeria Spina |
| EPI_ISL_2981937, EPI_ISL_2981938 | University Hospitals of Geneva, Laboratory of Virology | HUG, Laboratory of Virology and the Health2030 Genome Center | Ana Rita Goncalves; Deborah Penet; Emmanouil Dermitzakis; Henri Pegeot; Ioannis Xenarios; Keith Harshman; Laurent Kaiser; Lorenzo Cerutti; Melyssa Elies; Samuel Cordey |
| EPI_ISL_3014243, EPI_ISL_3014244, EPI_ISL_3014271 | Università Federico II - Dipartimento di scienze mediche traslazionali - Napoli | TIGEM | Antonio Grimaldi Patrizia Annunziata Francesco Panariello Teresa Giuliano Michele Cennamo Valentina Bouche Chiara Colantuono Lucio Di Filippo Mariano Fiorenza Anna Manfredi Marcello Salvi Giuseppe Portella Andrea Ballabio Davide Cacchiarelli |
| EPI_ISL_3270706 | Università Federico II - Dipartimento di scienze mediche traslazionali - Napoli | Telethon Institute of Genetics and Medicine (TIGEM) | Antonio Grimaldi Patrizia Annunziata Francesco Panariello Teresa Giuliano Michele Cennamo Valentina Bouche Chiara Colantuono Lucio Di Filippo Mariano Fiorenza Anna Manfredi Marcello Salvi Giuseppe Portella Andrea Ballabio Davide Cacchiarelli |
| EPI_ISL_2600742, EPI_ISL_2686030, EPI_ISL_2686039, EPI_ISL_2686043, EPI_ISL_2686045, EPI_ISL_2686048, EPI_ISL_2686049, EPI_ISL_2686050, EPI_ISL_2790250, EPI_ISL_2840651 | | | |
| see above | Università degli Studi di Perugia | Istituto Zooprofilattico Sperimentale dell'Abruzzo e Molise "G. Caporale" | Ancora M; Biagetti M; Calistri P; Camilloni B; Cammà C; Curini V; Delli Compagni E; Di Domenico M; Di Pasquale A; Giammarioli M; Lorusso A; Mangone I; Marcacci M; Mencacci A; Puglia I; Rinaldi A; Savini G; Scialabba S |
| EPI_ISL_3578428 | Universität Zürich | Institute of Medical Virology | Alexandra Trkola; Annette Audigé; Cyril Shah; Gabriela Ziltener; Guido Bloemberg; Jon Huder; Jürg Böni; Kevin Steiner; Maria Grünberg; Maryam Zaheri; Michael Huber; Riccarda Capaul; Stefan Schmutz; Verena Kufner |
| EPI_ISL_2761735 | Universitätsklinikum Münster Institut für Virologie | Robert Koch Institute | |
| EPI_ISL_3805216 | Utah Public Health Laboratory | Utah Public Health Laboratory | Erin L. Young; Kelly F. Oakeson; Olinto Linares-Perdomo; Pooja Gupta |
| EPI_ISL_3331654 | Valais Hospital, Central Institute | Valais Hospital, Central Institute | Alexis Dumoulin; Cedric Howald; Deborah Penet; Henri Pegeot; Ioannis Xenarios; Keith Harshman; Lorenzo Cerutti; Melyssa Elies |
| EPI_ISL_4000335 | Virginia Division of Consolidated Laboratory Services | Virginia Division of Consolidated Laboratory Services | Virginia Division of Consolidated Laboratory Services |
| EPI_ISL_2884605 | Virology Laboratory, Scientific Department, Army Medical Center | Virology Laboratory, Scientific Department, Army Medical Center | Anella Monte; Anna Anselmo; Antonella Fortunato; Filippo Molinari; Florigio Lista; Francesco Giordani; Giancarlo Petralito; Giandomenico Cerreto; Giulia Campoli; Lucia Nicosia; Marzia Cavalli; Riccardo De Sanctis; Rossella Brandi; Silvia Fillo; Vanessa Vera Fain |
| EPI_ISL_3841134 | Yale Clinical Virology Lab | Grubaugh Lab - Yale School of Public Health | Anderson Brito; Annie Watkins; Chaney Kalinich; Chantal Vogels; Isabel Ott; Jessica Rothman; Joseph Fauver; Kendall Billig; Mallery Breban; Marie L. Landry; Mary Petrone; Nathan Grubaugh; Tara Alpert; Tobias Koch |
| EPI_ISL_2617446 | laboratorio Microbiologia PO Cardarelli | laboratorio Microbiologia PO Cardarelli | Felice V.; Niro G. Scutella' M. |
